# Break-induced replication underlies formation of inverted triplications and generates unexpected diversity in haplotype structures

**DOI:** 10.1101/2023.10.02.560172

**Authors:** Christopher M. Grochowski, Jesse D. Bengtsson, Haowei Du, Mira Gandhi, Ming Yin Lun, Michele G. Mehaffey, KyungHee Park, Wolfram Höps, Eva Benito-Garagorri, Patrick Hasenfeld, Jan O. Korbel, Medhat Mahmoud, Luis F. Paulin, Shalini N. Jhangiani, Donna M. Muzny, Jawid M. Fatih, Richard A. Gibbs, Matthew Pendleton, Eoghan Harrington, Sissel Juul, Anna Lindstrand, Fritz J. Sedlazeck, Davut Pehlivan, James R. Lupski, Claudia M.B. Carvalho

## Abstract

**Background:** The duplication-triplication/inverted-duplication (DUP-TRP/INV-DUP) structure is a type of complex genomic rearrangement (CGR) hypothesized to result from replicative repair of DNA due to replication fork collapse. It is often mediated by a pair of inverted low-copy repeats (LCR) followed by iterative template switches resulting in at least two breakpoint junctions *in cis*. Although it has been identified as an important mutation signature of pathogenicity for genomic disorders and cancer genomes, its architecture remains unresolved and is predicted to display at least four structural variation (SV) haplotypes.

**Results:** Here we studied the genomic architecture of DUP-TRP/INV-DUP by investigating the genomic DNA of 24 patients with neurodevelopmental disorders identified by array comparative genomic hybridization (aCGH) on whom we found evidence for the existence of 4 out of 4 predicted SV haplotypes. Using a combination of short-read genome sequencing (GS), long-read GS, optical genome mapping and StrandSeq the haplotype structure was resolved in 18 samples. This approach refined the point of template switching between inverted LCRs in 4 samples revealing a DNA segment of ∼2.2-5.5 kb of 100% nucleotide similarity. A prediction model was developed to infer the LCR used to mediate the non-allelic homology repair.

**Conclusions:** These data provide experimental evidence supporting the hypothesis that inverted LCRs act as a recombinant substrate in replication-based repair mechanisms. Such inverted repeats are particularly relevant for formation of copy-number associated inversions, including the DUP-TRP/INV-DUP structures. Moreover, this type of CGR can result in multiple conformers which contributes to generate diverse SV haplotypes in susceptible *loci*.

## Introduction

DNA rearrangements can take many forms in a diploid genome, including deletions, duplications, inversions, and translocations, and can occur on a scale ranging from a few base pairs (bp) to several million base pairs (Mb) [1]. Among these diverse forms, complex genomic rearrangements (CGRs) are particularly intriguing due to their mostly unpredicted genomic architecture and potential impact on gene dosage and the consequences for human health.

CGRs represent a subset of genomic rearrangements that involve more than one breakpoint junction *in cis*, often resulting in the formation of highly complex genomic structures within a chromosome [2,3].

The duplication-triplication/inversion-duplication (DUP-TRP/INV-DUP) structure is one such CGR perturbation of genome integrity. This genomic instability can be incited by a given pair of inverted low copy repeats (LCRs) and result from two template switches (TS) during the process of DNA break repair [4]. This recurring DNA rearrangement end product structure is increasingly recognized for its significant roles in human disease, including neurodevelopmental disorders of childhood and adult onset neurodegenerative diseases, as well as its occurrence in cancer genomes [1,4–7].

In 2013 a genome-wide computational analysis of the GRCh37 human reference build elucidated 1,551 inverted LCRs that may predispose a region to local genomic instability generating a DUP-TRP/INV-DUP structure with 1,445 potentially disease-associated genes located within intervals of CNV formation due to the potential mutational event [8]. The variability in the gene copy number generated as a result of this structure and gene dosage effects, i.e., whether mapping to the duplicated or triplicated genomic interval, has been shown to influence disease severity and subsequent clinical heterogeneity [4,9–12].

The DUP-TRP/INV-DUP structure also contributes to X-linked diseases. It was originally described in the *MECP2* duplication syndrome (MDS) (MIM: 300260), a developmental disorder affecting boys caused by CNVs spanning the dosage sensitive gene *MECP2* at Xq28.

Approximately 26% of MDS have been reported to harbor a DUP-TRP/INV-DUP mediated by inverted LCRs downstream of the gene [4,13,14]. A more severe clinical phenotype is observed in patients with *MECP2* triplication [4,15,16]. Copy number events not spanning the *MECP2* gene but mediated by the same LCRs have also been implicated in duplication syndromes sometimes with incomplete penetrance [17].

Upstream of *MECP2* on the X chromosome, a different pair of inverted LCRs at Xq22.2 *PLP1* locus can also generate a DUP-TRP/INV-DUP CGR structure causing Pelizaeus-Merzbacher disease (PMD) (MIM: 312080); the CGR occurring with a prevalence of up to 20% in a combined cohort of 134 PMD subjects [18–20]. Triplications of *PLP1* are associated with a more severe phenotype in patients [21]. A majority of the pathogenetic effects for DUP-TRP/INV-DUP seem to be due to higher gene expression, i.e. gain of function. However, loss of function effect of this type of variant has also been reported; e.g. DUP-TRP/INV-DUP generated by LCRs within Xp21.1,disrupting exons 45-60 in the gene *DMD,* causes Duchenne muscular dystrophy (DMD) (MIM: 310200) [22].

Pathogenic DUP-TRP/INV-DUP structures in autosomes have been reported in multiple studies although at lower frequencies. In a cohort of 27 individuals with 17p13.3 duplication syndrome, 10% were found to have a DUP-TRP/INV-DUP structure formed by inverted *Alu* elements within that genomic region [23]. Amplifications of the gene *SCNA* within a DUP-TRP/INV-DUP structure at 14q22.1 has been associated as a causal factor in the progression of Parkinson disease (MIM: 168601) with duplications of the gene leading to a late-onset of the disease versus triplications which leads to an early-onset [24–26]. Triplications of the gene *CHRNA7* as result of a DUP-TRP/INV-DUP structure on chromosome 15 have been associated with neuropsychiatric phenotypes and other cognitive impairments including autism spectrum disorder (ASD) and attention-deficit disorder (ADHD) [27,28]. A pair of inverted LCRs on the long arm of chromosome 7 have been shown to generate the DUP-TRP/INV-DUP structure disrupting the gene *VIPR2* potentially impacting neurodevelopment and behavior [29]. In addition, errors in imprinting due to template switching in the formation of DUP-TRP/INV-DUP events can underlie cases of Temple Syndrome (MIM: 616222) and be associated with patients harboring multiple congenital malformations [30,31].

As the number of rare Mendelian disease traits and genomic disorders associated with this CGR continue to increase, our understanding of its implications in somatic cell mutagenesis and cancer genome evolution and progression is just beginning. Genomic instability is a hallmark of cancer cells and recent investigations into the role of structural variation in cancer genomes have begun to uncover the breath and complexity of SVs that are possible; the DUP- TRP/INV-DUP structure being identified as one of the twelve most prevalent SV mutational signatures found in that cohort [5]. The altered copy number state generated by the formation of this structure may lead to tumor-level selection pressures from aberrant gene dosage as well as the activation of oncogenes or inactivation of tumor suppressor genes [32–34]. Notably, both genomic disorders and cancer genome studies provided insights into the recombinant junctions and structural haplotype possibilities that may be formed.

Whether found within the constitutional or cancer genome, the DUP-TRP/INV-DUP structure is formed through genomic instability generated by a given pair of inverted repetitive elements in the genome [4,8]. This process begins during replication due to a fork collapse in an inverted LCR and subsequent strand invasion to resume replication mediated by homology of the inverted LCR pair [35]. DNA replication is continued in the reverse direction until a second fork collapse occur. Repair of the original strand may be accomplished by non-homologous end-joining (NHEJ) or microhomology mediated break-induced replication which resolves the second break [4,36].

Until recently, genomic sequencing technology limitations within large segments with high nucleotide sequence similarity stymied our ability to identify break-induced replication (BIR) breakpoints. Furthermore, we were previously unable to investigate the haplotype structure of the large sized (kb or Mb) segments involved in DUP-TRP/INV-DUP events in the context of a personal genome. Herein we sought to fully resolve those CGR structures and establish the structural variant haplotypes utilizing multimodal genomic analyses experimental and computational tools. We also define the molecular features of the inverted repeats that serve as substrates for recurrent pathogenic BIR at a specific Xq28 locus.

## Results

### Inverted LCR Pairs Generate Recurring DUP-TRP/INV-DUP Patterns

The current study includes 24 individuals who all harbor a DUP-TRP/INV-DUP genomic structure as initially identified by high-resolution aCGH. Out of the 24 samples within this cohort, 23 are males with a duplication (n=19) or triplication (n=4) spanning the *MECP2* gene causing MDS (Figure 1A, Table1). There is one female subject harboring a large (approximately 7.3 Mb) DUP-TRP/INV-DUP at Xq21 without overlapping *MECP2* (Figure 1B). who presented with developmental delay.

**Figure 1:**
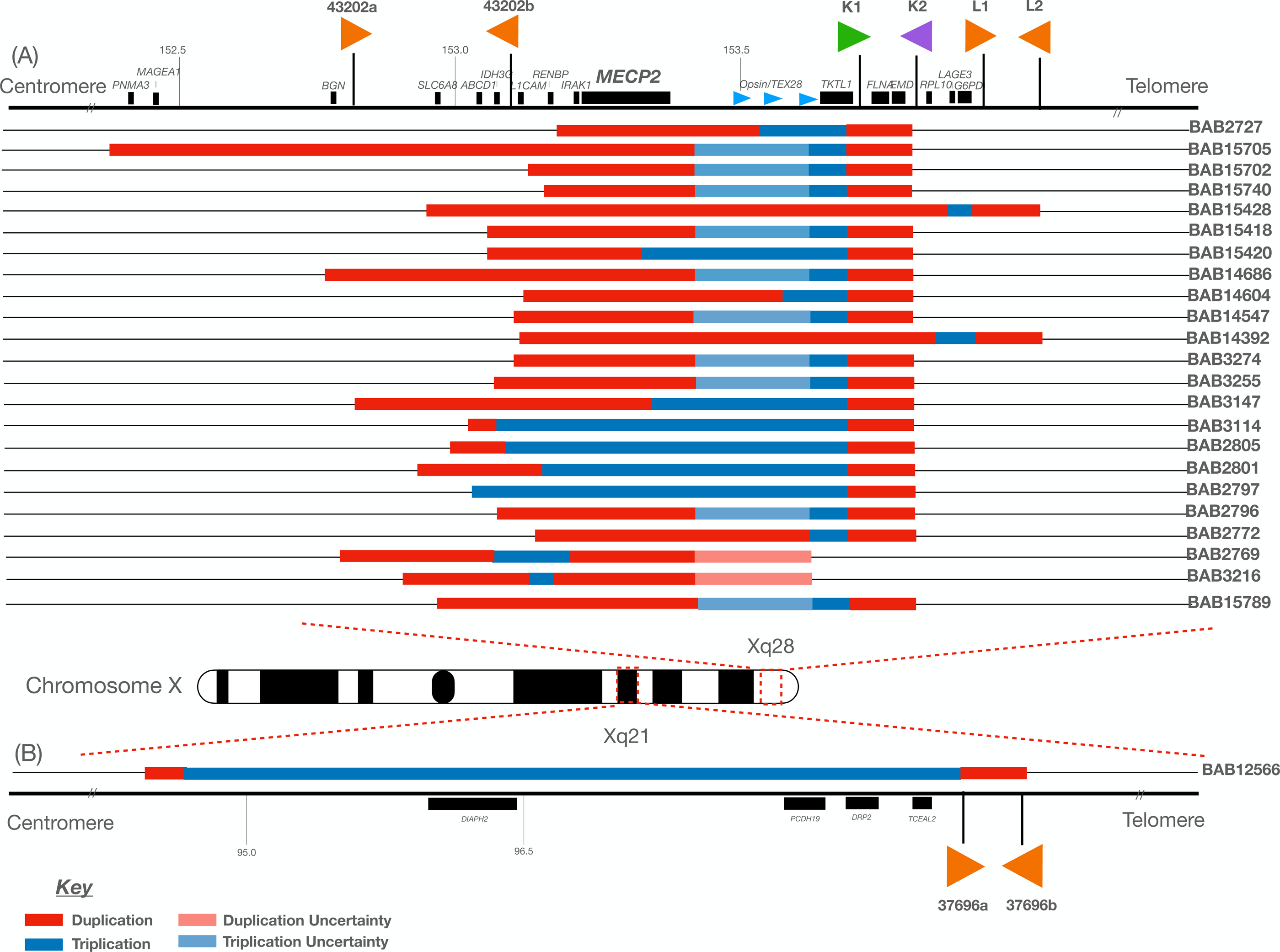
Individuals Carrying a DUP-TRP/INV-DUP Genomic Structure. A) Genomic region spanning Xq28 including the *MECP2* critical region is shown including the location of inverted low copy repeats (43202a/43202a; 43221a/43221b (K1/K2) (green and purple arrows); 43231a/43231b (L1/L2). The relative genomic locations of the duplication (red) and triplication (blue) are shown. Uncertainty as to the precise location of the start/end of either the duplication or triplication due to lack of probes on aCGH or low mapping quality in short-read GS within a given interval are depicted in light red and light blue respectively. B) A single individual female (BAB12566) is shown with a DUP-TRP/INV-DUP within Xq21 along with the relative position of inverted low copy repeat pairs (37696a/37696b). The naming scheme for 43221a/43221b (K1/K2) and 43231a/43231b (L1/L2) are derived from previous work detailing the DUP-TRP/INV-DUP structures at the *MECP2* locus on the X chromosome [4,13,68].

Based on customized aCGH data (hg19), the genomic rearrangements range in size from 417 kb to 7.4 Mb (from the beginning of the first duplication to end of the second duplication) (Additional File 1: Supplemental Table 1). The size of the initial duplication as well as triplication are variable ranging from 18.5 kb (BAB2797) to 854 kb (BAB15705) for the duplication region and 8.2 kb (BAB3216) to 7 Mb for the triplicated region (BAB12566). The size of the duplication and triplication events are dependent on the location of the second template switch forming junction 2. The size of the second duplication is dependent on the distance between the two initiating LCRs within a given genomic loci. In contrast, the size of the second duplication tends to be constant for the same *loci* since that CNV is often mediated by inverted repeats. In this cohort, the second duplication varies from 16 kb to 575 kb, but 18 out of 24 CGRs show the same 47 kb duplicated segment.

In this cohort four different pairs of inverted LCRs were identified to initiate the formation of the CGR event; three pairs are located at Xq28 (43202a/43202a; 43221a/43221b (K1/K2); 43231a/43231b (L1/L2)) whereas the fourth pair located at Xq22.1 (37696a/ 37696b). The size of each inverted LCR pair included in this study as well as the distance between the pairs varied. The smallest pair (43202a/43202b) was 926 and 917 bp in length with 98.12% similarity, separated by 317,810 bp. The next smallest (43221a/43221b(K1/K2)) was 11,455 and 11,446 bp in size with 99.23% similarity, separated by 37,614 bp. The next largest pair (43231a/43231b(L1/L2)) had both repeats approximately 35,968 bp in size and 99.92% similarity with 21,624 bp separating the two. The largest repeat pair identified in this cohort (37696a/37696b) was 140,562 bp and 140,621 bp in size; with a distance of 10,767 bp apart with 99.89% similarity (Additional File 1: Supplemental Table 2).

The breakpoint junction alignments for junction 2 in the structure for each sample were determined through either short-read GS, long-read PacBio HiFi sequencing, traditional Sanger dideoxy sequencing or a combination of methods. Out of a total of 24 samples, 14 samples showed a 1-9 bp microhomology at the breakpoint junction, one showing 7 bp of microhomeology, one with a blunt junction, two showed 1 bp to 2 bp insertion, one sample displayed an *Alu/Alu* fusion at the breakpoint and five displayed additional complexities such as templated insertions or microhomology-mediated deletions (Table 1). BAB15428 showed a chimeric fusion of *AluY* and *AluSx1*; these *Alu* repetitive elements were present in an inverted orientation on the reference genome and share 83% nucleotide sequence similarity. Although we have not obtained the breakpoint junction at the nucleotide level for this junction, we hypothesize that is an *Alu-Alu* mediated event (AAMR) [37]. Junction 2 in BAB15705 was found to be mediated by *AluYa8* and *AluJo* which were also in an inverted orientation and shared 36% sequence similarity when aligned to each other using the NCBI BLAST tool [38]. (Additional File 2) [4,13,14].

**Table 1:** Haplotype and junction resolution among DUP-TRP/INV-DUP patients.

| Patient Identifier | BH Identifier | Haplotype Structure | Jct1/Jct2 Single Molecule | MECP2 Inverted (Y/N) | Junction 1 | Inverted Repeat Pair | Junction 2 | Previous Studies |
| --- | --- | --- | --- | --- | --- | --- | --- | --- |
| BAB2727 | BH16106_1 | N/D | N/D | N/D | PB<br>(ChrX:153,613, 143-153,615,342) | 43221a/4<br>3221b<br>(K1/K2) | Microhomology 2 bp + templated insertion | Carvalho *et al.*, 2009 |
| BAB2769 | N/A | 3 | No | No | OGM, Sanger | 43202a/4<br>3202b | Microhomology: 2 bp + deletion | Carvalho *et al.*, 2011 |
| BAB2772 | N/A | 3 | Yes | Yes | OGM | 43221a/4<br>3221b<br>(K1/K2) | Microhomology: 3 bp | Carvalho *et al.*, 2011 |
| BAB2796 | BH16110_1 | 2 | Yes | No | OGM | 43221a/4<br>3221b<br>(K1/K2) | 2 bp insertion | Carvalho *et al.*, 2011 |
| BAB2797 | N/A | N/D | N/D | N/D | N/D | 43221a/4<br>3221b<br>(K1/K2) | 1 bp insertion | Carvalho *et al.*, 2011 |
| BAB2801 | BH15649_1 | 4 | Yes | No | OGM | 43221a/4<br>3221b<br>(K1/K2) | Microhomeology: 7 bp | Carvalho *et al.*, 2011 |
| BAB2805 | N/A | N/D | N/D | Yes | N/D | 43221a/4<br>3221b<br>(K1/K2) | Blunt junction | Carvalho *et al.*, 2011 |
| BAB3114 | BH14245_1 | 1 | No | Yes | OGM, PB<br>(ChrX:153,613, 143-153,615,342) | 43221a/4<br>3221b<br>(K1/K2) | Microhomology: 2 bp | Carvalho *et al.*, 2011 |
| BAB3147 | BH16111_1 | 6 | Yes | No | OGM | 43221a/4<br>3221b<br>(K1/K2) | Microhomology: 2<br>bp | Carvalho *et al.*,<br>2013 |
| BAB3216 | N/A | N/D | N/D | N/D | N/D | 43202a/4<br>3202b | Microhomology: 4<br>bp + templated<br>insertion | Carvalho *et al.*,<br>2013 |
| BAB3255 | BH16108_1 | 1 or 3 | No | Yes | OGM | 43221a/4<br>3221b<br>(K1/K2) | Microhomology: 2<br>bp | Carvalho *et al.*,<br>2013 |
| BAB3274 | BH16112_1 | 1 or 3 | No | Yes | OGM | 43221a/4<br>3221b<br>(K1/K2) | Microhomology: 3<br>bp + 9 bp + 5 bp<br>deletions | Carvalho *et al.*,<br>2013 |
| BAB12566 | BH13842_1 | 1 | No | N/A | OGM | 37696a/3<br>7696b | Microhomology: 2<br>bp | This Study |
| BAB14392 | BH15645_1 | N/D | N/D | N/D | N/D | 43231a/4<br>3231b(L1/<br>L2) | Microhomology: 1<br>bp | This Study |
| BAB14547 | BH15700_1 | 3 | Yes | Yes | OGM, PB<br>(ChrX:153,613,<br>143-153,615,342) | 43221a/4<br>3221b<br>(K1/K2) | Microhomology: 1<br>bp | This Study |
| BAB14604 | BH15701_1 | 3 | Yes | Yes | OGM, PB<br>(ChrX:153,613,<br>143-153,618,666) | 43221a/4<br>3221b<br>(K1/K2) | Microhomology: 1<br>bp | This Study |
| BAB14686 | BH15640_1 | 2 | Yes | No | OGM | 43221a/4<br>3221b<br>(K1/K2) | Microhomology: 2<br>bp | This Study |
| BAB15418 | BH16300_1 | 4 | Yes | No | OGM | 43221a/4<br>3221b<br>(K1/K2) | Microhomology:<br>2bp | This Study |
| BAB15428 | BH16301_1 | 1 or 3 | No | Yes | OGM | 43231a/4<br>3231b(L1/<br>L2) | *AluY/AluSx1* at<br>Breakpoint | This Study |
| BAB15702 | BH16609_1 | 6 | Yes | No | OGM | 43221a/4<br>3221b<br>(K1/K2) | Microhomology:<br>9bp | This Study |
| BAB15705 | BH16611_1 | 2 | Yes | No | OGM | 43221a/4<br>3221b<br>(K1/K2) | Microhomology:<br>6bp<br>(*AluYa8/AluJo*) | This Study |
| BAB15740 | BH16610_1 | 2 | Yes | No | OGM | 43221a/4<br>3221b<br>(K1/K2) | Microhomology:<br>2bp | This Study |
| BAB15789 | N/A | 2 | Yes | No | OGM | 43221a/4<br>3221b<br>(K1/K2) | Microhomology:<br>2bp | This Study |
| BAB15420 | BH16299_2 | N/D | N/D | N/D | N/D | 43221a/4<br>3221b<br>(K1/K2) | Complexities | This Study |

Majority of patients contains *MECP2* in duplicated segment A/A'; except BAB3147, BAB15420 (truncated *MECP2* – B); BAB2801, BAB2805, BAB3114, BAB2797 (triplicated *MECP2* – B); BAB2769, BAB3216 – C.
N/A: Not available; N/D: Not determined; OGM: Optical Genome Mapping; PB: PacBio HiFi

### Modeling SV Haplotype Conformers of Mutational Events

The previous identification of triplications being inverted and embedded within duplicated sequence provided evidence for two breakpoint junctions that occur *in cis* forming the DUP-TRP/INV-DUP structure with inverted LCRs acting as a recombinant substrate through BIR to generate this type of structure [4]. The model for formation of each haplotype conformer is predicated on which inverted LCR was used to generate the event and the distance and location that the template then switches back to the reference strand continuing through replication and resolved by extended replication, repair by non-homologous end joining or formation of half-crossover [5].

We developed a prediction model based on a LCR-pair within the *MECP2* locus 43221a (K1) and 43221b (K2) which can be used to infer if the template switch occurred from the first LCR (K1) to the second LCR (K2) on the sister chromatid via homologous recombination and re-initiation of the replication fork to resume replication in the opposite orientation (haplotype conformer 1 and 2) or a template switch from the second LCR (K2) to the first LCR (K1) on the opposite sister chromatid (haplotype conformers 3 and 4) (Figure 2). Both form a chimeric LCR, i.e. recombinant representing junction 1, and a recurrent duplication (DUP2) spanning the genomic segment in between the inverted repeats. Junction 2 results from a second template-switch triggered by double-stranded break (DSB) or replication fork stalling/collapse which will produce the inverted triplication segment and DUP1. The size of the inverted triplication and DUP1 depend on the location of the second template switch The linearized final structure thus allows inferences as to the temporal replication fork jumps, i.e. iterative TS of the progressing replication fork, forming junctions 1 and 2 with the formation of a chimeric LCR. This same prediction model can be inferred for all inverted LCRs detailed in this study (43202a, 43202a; 43221a/43221b (K1/K2); 43231a/43231b (L1/L2), 37696a, 37696b) as well as other pairs that generate additional DUP-TRP/INV-DUP events at other positions genome-wide.

**Figure 2:**
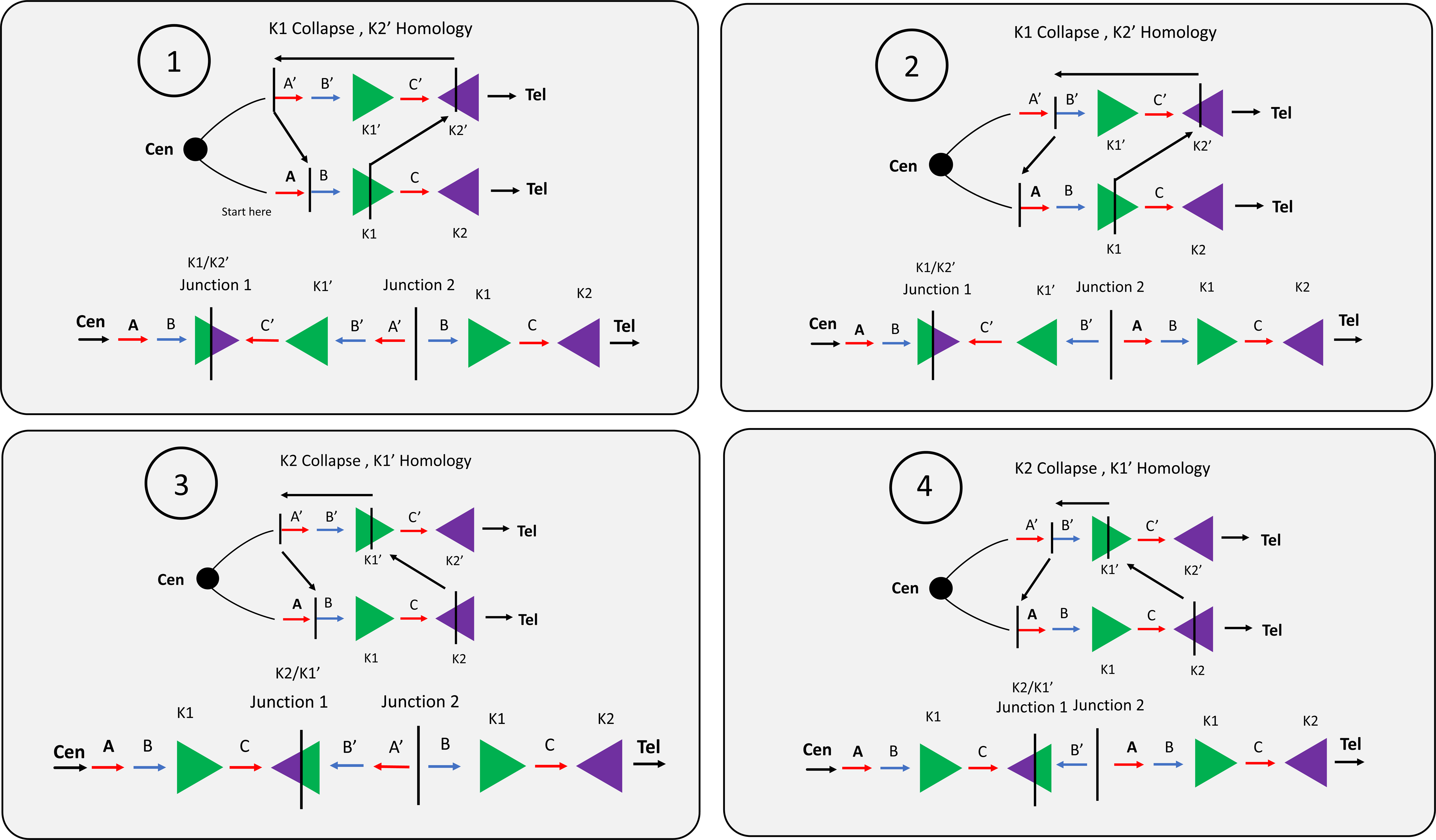
Predictive Model for DUP-TRP/INV-DUP Formation At least four haplotypes sub-structures can be derived from rearrangement involving a pair of inverted low-copy repeats. This figure depicts the LCRs K1 and K2 (green and purple arrowheads) within the *MECP2* locus during an intrachromosomal event. The same model can be applied to all DUP-TRP/INV-DUPs formed through inverted LCRs. The formation of the DUP- TRP/INV-DUP event may start due to a replication fork stall and collapse at or nearby the LCR (K1, denoted as a green arrowhead. Homology drives strand invasion at the inverted LCR (K2’) on the opposite strand (denoted in purple), producing junction 1. DNA replication continues in the opposite direction until a second replication fork collapse and repair on the original strand through either microhomology-mediated break-induced replication (MMBIR) or non-homologous end-joining (NHEJ) resolves the second junction. . The four conformer possibilities shown here are determined by the replication fork collapsing and jumping (template switch denoted by dashed black arrows) from either K1 to K2’ or K2 to K1’.

### Structural Variant Haplotype Conformers within DUP-TRP/INV-DUP Events

In samples for which cell lines or whole blood was previously frozen and available, we utilized ultra-high molecular weight DNA and OGM to phase genomic fragments in the context of the larger structure and diploid genome through the visualization of single DNA molecules containing genomic segments in the structure (Figure 3). Out of 19 samples on which OGM was performed we could phase the DUP-TRP-DUP CGR into 4 distinct and predicted substructures that are possible through two template switches (Figure 2, Additional File 3: Supplemental Figure 1). The four identified conformers include i) the initial duplication, triplication and final duplication all in an inverted orientation (haplotype structure 1) (BAB3114, BAB12566) ii) the triplication and final duplication in an inverted orientation (haplotype structure 2) (BAB2796, BAB14686, BAB15705, BAB15740, BAB15789) iii) the triplication and initial duplication in an inverted orientation (haplotype structure 3) (BAB2772, BAB14547, BAB14604) (Figure 3) and finally iv) just the triplication in an inverted orientation (haplotype structure 4) (BAB2801, BAB15418) (Figure 3, Table 1). Two samples (BAB3147 and BAB15702) were found to harbor an additional structure (haplotype structure 6), that is formed through the same mechanism as haplotype structure 2 but leads to an appearance of an inversion of only the triplication due to a potential ancestral inversion of the segment C relative to reference (Additional File 3: Supplemental Figure 1). For the LCR K1 and K2, a polymorphic inversion is known to be present in approximately 18% of the population of European descent [39].

**Figure 3:**
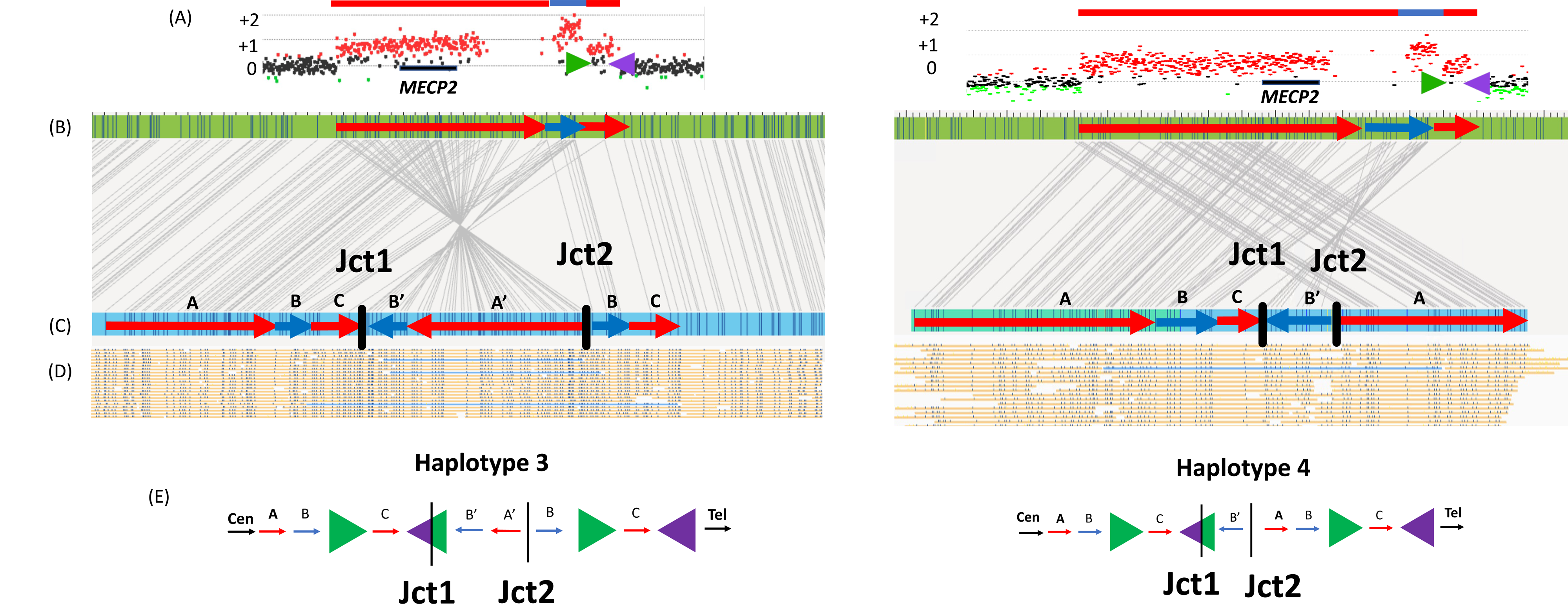
Haplotype Resolution using with Optical Genome Mapping Structural haplotype determination and conformer configuration was established based upon single molecule support through junction 1 and junction 2 *in cis.* Two samples in the cohort are highlighted, BAB14604 (left) and BAB15418 (right). A) ArrayCGH plots for each sample show a similarly sized DUP-TRP-DUP event both mediated by inverted low-copy repeats (shown as green and purple arrows) downstream of *MECP2* (black rectangle). B) Optical genome mapping (OGM) reference (green rectangle) shows *in silico* motifs throughout the *MECP2* locus. The red arrows correspond to the duplicated segments whereas blue arrow corresponds to the triplicated segments, length of the CNVs are proportional to the aCGH CNV. C) Proband OGM de novo assembly is shown in blue rectangle. Sequence motifs are aligned to the reference shown as connecting gray lines enable ‘restriction fragment genome mapping’ and pattern recognition. Red and blue arrows are overlayed to represent the position and orientation of each amplified genomic fragment within the DUP-TRP-DUP structure. The connection point forming junctions 1 and 2 are shown as black vertical dashed lines/bars. D) Single DNA molecules that span both junctions 1 and 2 are highlighted in blue confirming that both junctions are present *in cis*. E) Hypothesized resolved haplotypes based on CNV and *in cis* junction analysis. Although both samples show nearly identical aCGH patterns, BAB14604 has conformer haplotype 3 and BAB15418 shows conformer haplotype 4. All 4 predicted conformer haplotype structures were identified (Additional File 2).

The two breakpoint junctions (Jct1 and Jct2) are identical within all sub-haplotype structures, only the orientations of genomic fragments in the structure differ between distinct haplotype conformers when the structure is visualized in a linear fashion (Figure 2, Additional File 3: Supplemental Figure 1). Single DNA molecule resolution *in cis* through both breakpoint junctions 1 and 2 (Figure 3D) enables an interpretation of each individual haplotype structure given the rearrangements occur in a male on the X chromosome and not on an autosome. For samples that we did not have a single DNA molecule that spans both junctions 1 and 2 (BAB2769, BAB3255, BAB3274, BAB15428) we could not definitively refine the structure to a single haplotype but could refine it to either haplotype 3 or 1.

Within DUP-TRP/INV-DUP, all samples with triplications encompassing the entire *MECP2* gene have *MECP2* an inverted orientation (BAB3114, BAB2805, BAB2801, BAB2797). Additionally, haplotype structures 1 and 3 have the initial duplication (including the *MECP2* gene) in an inverted orientation on the amplified genomic fragment (BAB15428, BAB14604, BAB14547, BAB3274, BAB3255, BAB2772) (Table 1). The remainder of the samples with an identified haplotype structure include the amplified copy of *MECP2* that appears to be present in the structure in a proposed haploid human genome reference orientation.

### Long-read sequencing facilitated breakpoint mapping within Inverted Repeats

PacBio “HiFi” facilitated the ability to generate highly accurate reads though repetitive sequences that were not possible using previous short-read technologies due to low mapping quality within a given region. For the *MECP2* region, LCRs K1 and K2 are approximately 11 kb in length and 99.23% similar (Hg19) (Additional File 1: Supplemental Table 2). The CGR generates a hybrid K1/K2 resulting from copy-number event in addition to additional copies of either K1 or K2. All the long-reads that span the LCRs are mapped to the reference, but we can refine the exact reads that map to the K1/K2 hybrid as they will present soft-clipping junctions flanking the genomic border of the LCRs (Figure 4). This approach enabled the re-mapping of the hybrid reads which revealed the recombinant breakpoint junctions within the LCRs in 7 samples (BAB2727, BAB2801, BAB3114, BAB3147, BAB3255, BAB14547, BAB14604). These reads indicate the connection of LCR K1 and K2 forming Jct1 within the DUP-TRP/INV-DUP structure that as a result form a recombinant or chimeric LCR.

**Figure 4:**
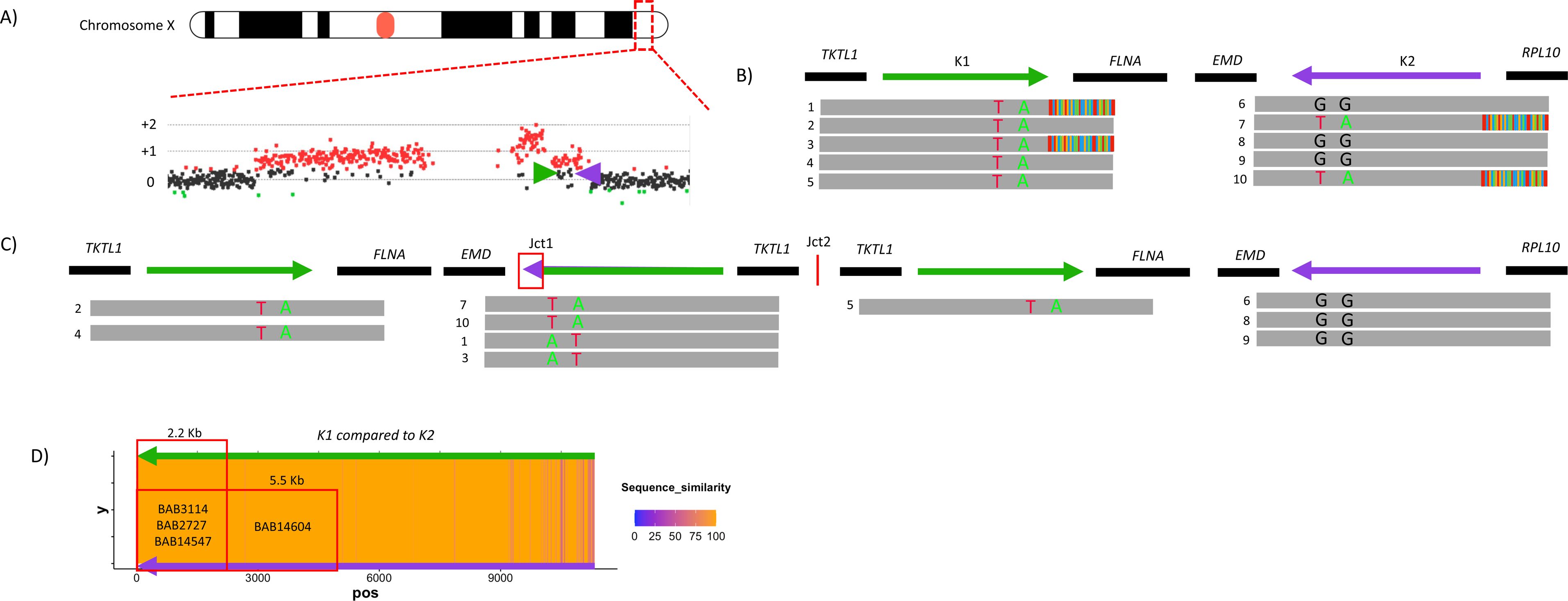
Refined K1/K2 Position of LCR Template Switch Visualization of paralogous sequence variants through inverted LCRs allows for a determination of the relative position of replication fork collapse, subsequent 5’ -end resection, and 3’ -end strand invasion to the point of homology within the inverted LCRs for BAB2727. A) ArrayCGH showing a DUP-TRP-DUP structure with *MECP2* locus highlighted and zoomed in. The inverted LCRs K1 and K2 (shown as green and purple arrows) are located flanking the terminal / 3’ end duplication in the structure. B) Position of K1 and K2 are shown with representative HiFi data below, highlighting sequence reads that span the region. Ancestorial reads denotes HiFi reads are uniquely aligned with LCR (e.g., read 2, 4, and 5). Breakpoint reads denote HiFi reads that begin in unique sequence and show soft clipping as they exit the LCR (e.g., read 1, and 3). Paralogous sequence variants (PSVs) are visualized in LCR K2 with the green (A nucleotide) and red (T nucleotide) positions that are found breakpoint reads in K2 and are present within points of homology in K1. C) Linearized structure showing the reads found within each position. The chimeric K1/K2 shows the positioning of PSVs used to refine the position of Jct1. D) Percentage of uniquely aligned base in slide window of 20 base pair, i.e. sequence similarity were shown as a heatmap. A ‘hot’ color – orange, denotes to 100% match, while a ‘cold’ color- purpole, denotes reduced similarity. The position of the PSV can be used to estimate the distance of replication fork proceed before the template switch to K2 occurred. Samples in this cohort could be narrowed to a 2.2 kb or 5.5 kb region.

Moreover, the high accuracy rate of PacBio HiFi sequencing allows identification of single nucleotide changes even within highly similar sequences i.e. paralogous sequence variants or PSVs [40,41]. Single nucleotide variation between the LCRs enables one to refine the point from where BIR uses homology and the second LCR; i.e. the recombinant join point (Figure 4). We could further refine the ‘crossover uncertainty’ to approximately 5.5 kb in one sample (BAB14604) and 2.2 kb in three samples (BAB14547, BAB2727, BAB3114) (Figure 4D). The uncertainty range is based on the presence of informative PSV (i.e SNPs that are present in K2 but map to the same reference location in K1). If no informative SNPs were present within a given breakpoint spanning read, the uncertainty range as to where the homology-driven template switch occurred cannot be determined within a given sequence read.

### CRISPR-Cas9 Enrichment (ONT) and Strand-Seq Orthogonally Validate Fusion Junction Formation and Haplotype Structure

CRISPR-Cas9 enrichment and subsequent ONT sequencing for the *MECP2* critical region was performed on three individuals in family BH14245): including BAB3114 (proband), BAB3115 (carrier mother) as well as BAB3121 (maternal grandfather). Targeted nanopore sequencing and Cas9-guided adapter ligation on ultra-high molecular weight extracted DNA allowed for sequencing within the *MECP2* region through the LCRs K1 and K2 as well as through duplication-triplication-duplication event with single DNA molecule resolution of the structure.

ONT long-read sequencing through both K1 and K2 LCRs orthogonally validated the informative PSVs that were detected within K2 at position ChrX:153,615,342 and ChrX:153,615,645 on reads that span the chimeric LCR and that are present within the same position on K1 as independently visualized though HiFi sequencing data (Additional File 4).

Additionally, the presence of a large single 530 kb read that spanned (in a single molecule) the duplication and triplication regions enabled refinement of the haplotype structure in the context of the larger CGR (Additional File 4). This method allowed for an additional orthogonal confirmation of the haplotype structure that was observed in the OGM analysis of the same sample (BAB3114) (haplotype conformer 1).

The implementation of Strand-Seq for samples BAB3114 and BAB14547 provided an orthogonal confirmation of the haplotype structures 1 and 3, respectively. Strand-Seq was particularly important to validate the haplotype of BAB3114 because there were no molecules spanning both junctions 1 and 2 in the optical mapping data for BAB3114.

## Discussion

We studied 24 individuals harboring a DUP-TRP/INV-DUP structure mediated by inverted low copy repeats including three sets that reside at the *MECP2* critical region and an additional pair not involving *MECP2.* The size of the genomic fragments as well as the fact that breakpoint junctions may occur within repetitive regions of the genome previously obfuscated resolution of the structural variant haplotype. Utilizing data from high-resolution aCGH as well as short and long-read GS (ONT and PacBio HiFi), OGM and StrandSeq enabled elucidation of the recombinant events within each SV haplotype and visualized the individual conformers (Table 1, Additional File 1: Supplemental Table 3).

The formation of the DUP-TRP/INV-DUP structure was hypothesized to occur by a combination of BIR and MMBIR or NHEJ using inverted LCRs as the recombinant substrate during the process of generating junctions 1 and 2 in the CGR conformer structure [4]. A combination of Southern blot and Sanger dideoxy sequencing of these breakpoint junctions revealed the inverted orientation of the triplication and its connections to both the initial and final duplications [4]. Although the majority of individuals present with only two template switches forming two “breakpoint junctions”, there are four theoretical SV haplotypes that can occur resulting from genomic fragments included in the structure in either a reference or inverted orientation [5,7]. Previous sequencing methodologies did not identify structural haplotype differences in each sample harboring a DUP-TRP/INV-DUP partly due to technical limitations [4,19,30,31], although they were recently predicted [5].

Within this study, all four hypothesized SV haplotypes were detected (Figure 2) in addition to new ones (haplotype 6) (Table 1). Utilization of OM with long- read ONT and HiFi sequencing data allowed for i) identification of relative orientation of each genomic fragment in context of the chromosome haplotype and ii) breakpoint junction sequence to the base-pair level resolution. The formation of this genomic aberration may cause a pathogenic consequence through gene fusion, gene interruption and/or new gene formation events [42] (Additional File 3: Supplemental Figure 2). Identification of these SV haplotype structures presents a previously unknown level of complexity to SV mutagenesis. Specifically, the initial LCR used to mediate the rearrangement can now be proposed and the region within that LCR can be delineated.

Based on the experimental data provided by OGM, long-read GS and CRISPR-Cas9 targeted sequencing for this cohort we developed a predictive model that can be applied to any other susceptible loci in the genome [8]. The implementation of StrandSeq allowed for the refinement of haplotype structure 1 in BAB3114 due to molecule size limitations in OGM data that limited the ability to refine the haplotype from one data source alone (Additional File 5).

Additional derivations of these 4 observed haplotypes can be formed through this process occurring as interchromosomal versus intrachromosomal events as well if there are SV inversion alleles in a reference region as was observed with BAB3147 and BAB15702 (Additional File 3: Supplemental Figure 1).

We elucidated 4 pairs of inverted LCRs that act as recombinant substrates for the formation of this genomic event (Additional File 3: Supplemental Figure 3). Shared nucleotide similarity ranges from 99.12% to 99.92% whereas there is significantly differences in size and distance separating them (Additional File 1: Supplemental Table 1). Most of the samples within this cohort (19/24) have CGRs mapping to the LCR pairs 43221a/43221b (K1/K2), downstream of the *MECP2* locus. K1/K2 pair has the third largest separation distance (37,614 bp) and are the second smallest sized LCR pairs (11,446 bp). The average size of the duplication 1 was 365,269 bp with a median size of 320,848 bp. The distance from *MECP2* for the first LCR in 43221a/43221b (K1/K2) is 201,074 bp while the distance from *MECP2* to the LCRs 43231a/43231b(L1/L2) is 420,500 bp which is 96,118 bp larger than median size of the initial duplication. It is possible that an unstable replication fork is generated as a result of BIR within an inverted LCR moving in a reverse direction (generating the initial duplication) [43]. The instability possibly limits the distance replication can continue in the reverse direction before the lagging strand disengages and integrates sequence on the original strand in a MMBIR model or through DSB repair in a NHEJ model. The preference for 43221a/43221b (K1/K2) is likely an ascertainment bias within our cohort due to the distance the pairs sit from a dosage sensitive gene - in this case *MECP2*. Other inverted pairs on the X chromosome are known to mediate the same type of CGRs. For instance, the inverted LCR pairs A1a/A1b (38209a/38209b), downstream of the gene *PLP1* at Xq22, form DUP-TRP/INV-DUP structures causing Pelizaeus-Merzbacher disease MIM: 312080 in 20% of the cohort. A1a/A1b are 20349 and 20353 bp in size and share 99.27% sequence similarity with a distance of 60,043 bp apart. Importantly, inverted repetitive elements such as *Alu’*s with similarities as low as 85% (*Alu*Sg/*Alu*Sg, *Alu*Sx3/*Alu*Sz) have also been identified as mediators of the CGR event as seen in 17p13.3 [23]. However, contrary to K1/K2, L1/L2 at Xq28 or A1a/A1b at Xq22 that are responsible to multiple independent events involving those loci, 17p13.3 *Alu’s* have not been reported in more than a single DUP-TRP/INV-DUP event in a given loci. In aggregate, these data support a combined role for inverted repeat size (> 10 kb), shared nucleotide similarity (> 99%) and proximity (< 100 kb) in the formation of this type of CGR.

The region of uncertainty in which the recombination crossover takes place within the first or second LCR in 43221a/43221b (K1/K2) was narrowed to a 2.2-5.5 kb The segment defined in 4 crossovers shares 100% sequence similarity between the LCRs K1 and K2 supporting the hypothesis that large stretches of homology are substrates non-allelic recombination. While this length and sequence similarity is comparable to those that generate rearrangements through non-allelic homologous recombination (NAHR) process [44,45], the recombination formed within the LCR generate inversions accompanied by copy-number variation that are further resolved by a second mechanism (MMBIR or NHEJ) which are consistent with an unstable BIR event [46].

In summary, optical genome mapping and long-read GS approaches facilitated phasing and assembly of a clinically relevant recurrent SV structure– the DUP-TRP/INV-DUP CGR at Xq28 (Figure 5). Furthermore, this work provides insights into BIR, a relevant molecular mechanism contributing to formation of inversion alleles and CGRs. The ability to resolve SV haplotypes and to determine gene structure perturbations will guide our understanding of neomorphic allele and gene fusion formation. Complex structural variation may have pathogenic consequences as well as beneficial clinical ramifications [47], but also the potential of driving genome evolution [48,49].

**Figure 5:**
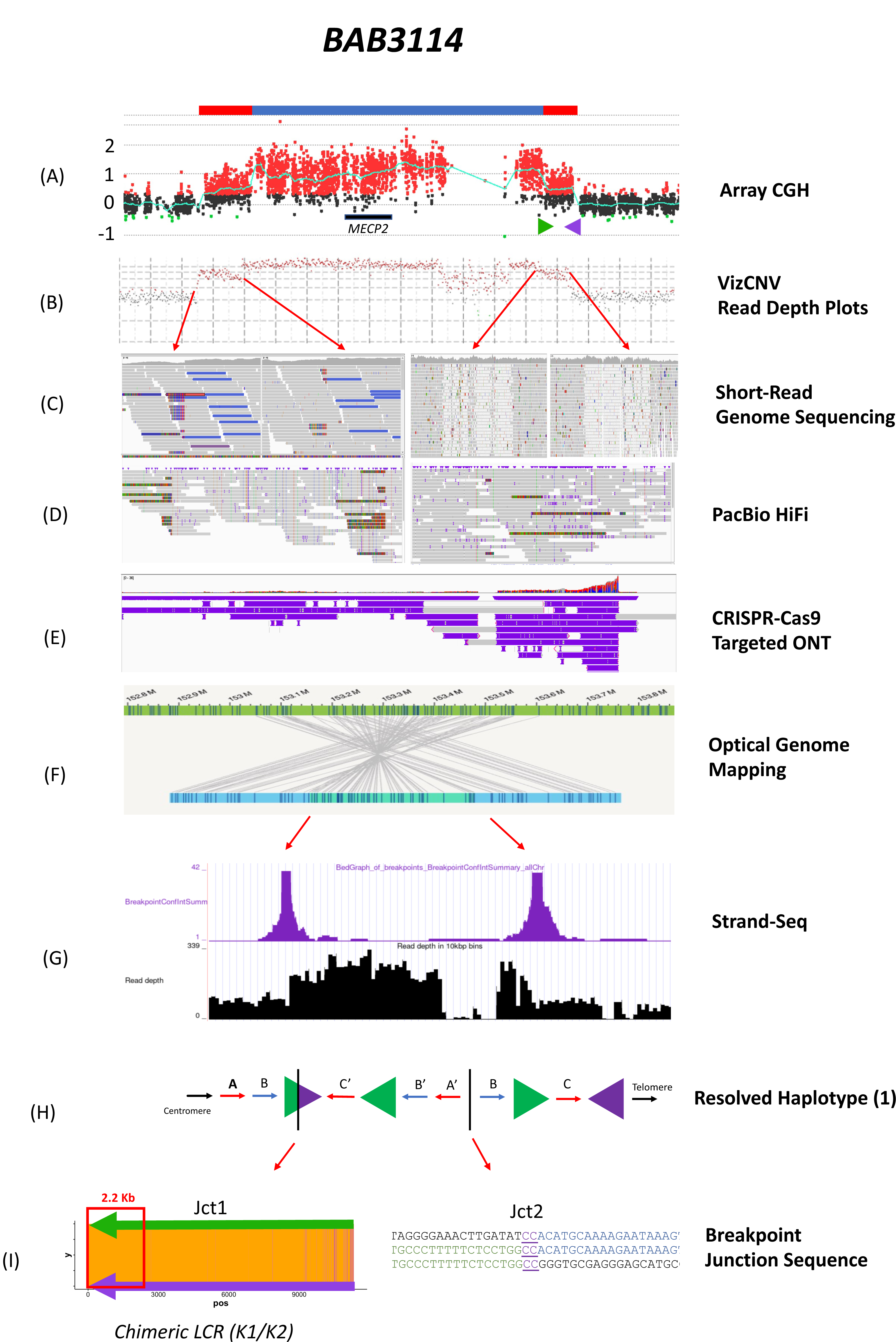
Multipronged Approach Resolving DUP-TRP/INV-DUP Events Sample BAB3114 including the methodology used to fully resolve the structural variant haplotype and breakpoint junctions 1 and 2. A) Array comparative genomic hybridization showing a duplication-triplication-duplication (DUP-TRP-DUP) structure at Xq28 including the *MECP2* gene and LCRs K1 (green arrow) and K2 (purple arrow). B) Illumina short-read genome sequencing showing the read depth for the region as visualized in the VizCNV plotting program [62]. C) Red arrows denote the regions of copy number change as seen in the short-read sequencing in IGV. Of note soft clipping can be seen in the regions of unique sequence (left) versus the unmapped reads at the region with K1 and K2 due to sequence similarity of the region. D) PacBio HiFi data shows the reads that include the breakpoint region (shown as soft clipping) within both junction 1 and junction 2. E) CRISPR-Cas9 targeted ONT facilitated ultra- long molecule (>500kb) sequencing to capture the haplotype structure within a single DNA molecule. F) Bionano OGM shows orientation and connection points of amplified genomic fragments forming junctions 1 and 2 within the structure. G) Strand-Seq data showing the points of breakpoint (purple peaks) with the inverted genomic sequence between. H) Resolved haplotype structure 1 for BAB3114 shows the triplication and initial duplication in an inverted orientation. I) Junction 1 shows a heatmap of K1/K2 similarity. The point of fork stall/collapse and strand invasion to the inverted LCR occurs within a 2.2 kb stretch of the LCR K1/K2 (as shown in the red arrow). Junction two can be determined to nucleotide level resolution and shows a 2 bp microhomology.

Untangling complex genomic events including the DUP-TRP/INV-DUP enables better comprehension of the underlying molecular basis of Mendelian disease but also enlightens the mechanisms leading to genomic instability and provides insights into cancer mutagenesis and evolution of genes and genomes. Drastic and rapid changes to the genome caused by complex structural variation such as that observed herein has the effect of generating changes beyond simple Watson-Crick single base-pair ‘editing’ [50]. Through the generation of CGRs, large portions of the genome are moved, reordered, inverted and connected in novel ways not previously seen, driving new and unknown possible outcomes and involve previously cryptic genomic complexities [51–53]. The subsequent gene expression and clinical effect(s) of such genomic perturbations must be further investigated.

## Methods

### Patient Enrollment

Study participants (N=24) (Table 1) were consented according to the Institutional Review Board for Human Subject Research at Baylor College of Medicine approved protocols: H-29697, H- 20268, and H-47127 or Pacific Northwest Research Institute WIRB #20202158. Whole blood samples (3-10mL) were collected via peripheral venous blood draw in Ethylenediaminetetraacetic acid (EDTA) and Acid Citrate Dextrose (ACD) vacutainer tubes from patients diagnosed with *MECP2* Duplication Syndrome. Genomic DNA from patients and family members was isolated from blood according to standard procedures.

### Array Comparative Genomic Hybridization

To evaluate copy-number changes in chromosomes X and Y, we designed a custom 4 x 180K tiling-path oligonucleotide microarray spanning the entirety of X and Y, including the *MECP2* region on Xq28 (Hg19). The custom 4x180K Agilent Technologies microarray (AMADID #086099) was designed using the Agilent Sure Design Website version 6.9.1.1 (https://earray.chem.agilent.com/suredesign/) on NCBI Build 37. We selected 143,860 probes interrogating chrX: 1-155,270,560 for a median probe spacing of 797 bp and 23,912 probes covering chrY: 1-59,373,566 for a median probe spacing of 425 bp. Arrays were run according to the manufacturer’s protocol (Agilent Oligonucleotide Array-Based CGH for Genomic DNA Analysis, version 7.2, Agilent Technologies) with modifications [4] on probands, mothers, and in select cases, fathers and maternal grandparents (if available) to determine inherited vs *de novo* rearrangements. Coordinates for each CNV observed along those chromosomes were annotated. The genomic context where breakpoint junctions occur were investigated using UCSC Genome Browser GRCh37/hg19 Assembly (http://genome.ucsc.edu) [54] for information about the presence of repeats, low-copy repeats, and genes or pseudogenes. To identity inverted and direct repeats that may mediate DUP-TRP/INV-DUP events, we mapped the breakpoints to genome-wide maps of highly-similar intrachromosomal repeats [55].

### Short-Read Genome Sequencing

Short-read libraries were prepared at the BCM Human Genome Sequencing Center (HGSC) with KAPA Hyper PCR-free library reagents and sequenced with an Illumina NovaSeq instrument using standard sequencing procedures which yielded 30X GS 150 bp paired-end sequence reads on average. Raw sequence read data was processed with the HGSC HgV analysis pipeline which applies BWA-MEM software to map the reads to NIH-compliant GRCh38 reference genome. A subset of samples (N=2) had sequencing performed at the National Genomics Infrastructure (NGI), in Stockholm, Sweden using an Illumina 30X PCR-free paired-end (PE) approach [56].

### Optical Genome Mapping

Ultra-high molecular weight (UHMW) DNA was isolated from frozen EDTA blood or cryopreserved cells following manufacturer instructions (documents 30246 rev F, 30268 rev D). In short, the frozen samples were first thawed in a 37 °C water bath. Then blood and cell samples were counted using a either a HemoCue WBC System (HemoCue AB) or hemocytometer, respectively. A volume containing 1.5 million cells was pelleted via centrifugation at 2,200x g for 5 minutes. The pellets were resuspended in a DNA stabilization buffer and treated with proteinase K in lysis and binding buffer. Cryopreserved cell samples were also treated with RNAse A at this step. After proteinase K digestion, samples were treated with PMSF, bound to a nanobind disk, washed, and eluted. DNA extracts were homogenized via end-over-end rotation and incubated at room temperature overnight before fluorescent labeling. For each sample, 750 ng of DNA was labeled at the recognition site CTTAAG using Direct Labeling Enzyme 1 (DLE-1) and counter-stained following manufacturer instructions (document 30206 Rev F). Labeled DNA was imaged on a Saphyr Gen2 platform, collecting 400X-1500X effective coverage for each dataset. *De novo* assembly and structural variant calling was performed using Solve version 3.7 as described by the Bionano Solve Theory of Operation: Structural Variant Calling (Document 30110 Rev J). To reduce the computation time for *de novo* assemblies, each dataset was down-sampled to 250X effective coverage by filtering for the longest molecules of each data set.

Structural variants (SV) were called against the human reference genome Hg19. Using the Variant Annotation Pipeline, SV calls were annotated and compared to the Bionano control sample database, which contains >600,000 SV calls from >150 phenotypically normal individuals from >26 populations. See the Bionano Solve Theory of Operation: Variant Annotation Pipeline (document 30190 revision H) for more details.

### Pacific Biosciences (PacBio HiFi)

Whole genome sequencing was performed at the HGSC at Baylor College of Medicine using long reads from the Pacific Biosciences sequencing platform. After DNA quality was assessed using Qubit and pulsed-field gel electrophoresis (PFGE), 15ug genomic DNA was used to construct a library using the SMRTbell Express Template Preparation Kit 2.0 with an average fragment length of 15 kb. Using the PacBio Sequel II instrument, two SMRTcells were sequenced per library for an average of 43Gb of HiFi reads per sample with an average coverage of 15-20x.

### Oxford Nanopore (Promethion)

Long read whole genome sequencing data was also generated using the Oxford Nanopore Technologies sequencing platform. After DNA quality was assessed using Qubit and pulsed-field gel electrophoresis (PFGE), A library was constructed with 15ug input genomic DNA using the SQK-LSK110 ligation sequencing kit with an average fragment length of 15Kb. Using the Oxford Nanopore Technologies Promethion instrument, one flowcell was sequenced per library with an average yield of 90Gb per sample. Basecalling was performed using Guppy version 4.3.4+ecb2805 and methylation analysis with Megalodon version 2.3.1 [https://github.com/nanoporetech/megalodon] using the default parameters of the program.

### Additional Data Processing and Analysis of Long Read Sequencing Data

Using PRINCESS version 1.0 [57], an in-house developed data processing workflow for long reads sequencing, reads were aligned to GRCh37 and phased variant calls were generated for SVs and SNVs. Briefly, PRINCESS will start by aligning reads using the appropriate parameters based on the type of sequencing technology using Minimap2 version 2.17 [58] followed by calling SVs using Sniffles version 1.12 [59] and will identify SNVs and indels using Clair3 [60]. Finally, PacBio HiFi data was processed using the same methods using PRINCESS with the read-option set to CCS (-r ccs). For SVs from both sequencing platforms, variants were filtered based on read support to require a maximum ∼25k SVs per sample.

### Nanopore Cas9 Enrichment and Sequencing for BH14245 Family

Patient derived immortalized lymphoblastoid cell lines were cultured in RPMI-1640 media (ATCC) supplemented with 10% FBS (ATCC) and 1% penicillin/streptomycin/amphotericin B (Thermo Fisher). Cells were maintained at 37°C in 5% CO2. *Genomic DNA Extraction and Purification:* Genomic DNA was extracted from 5 M cells using the Gentra Puregene Cell Kit (Qiagen) following the manufacturer’s instructions. Extracted DNA was further purified by isopropanol precipitation. For Adaptive Sampling experiments, DNA was sheared to approximately 20 kb using g-tube (Covaris). Ultra- high molecular weight (UHMW) DNA was purified using the Nanobind CBB Big DNA Kit (Circulomics) following the manufacturer’s instructions, eluted into 150 µL Circulomics EB containing 0.02% Triton-X100, and equilibrated overnight at room temperature. DNA was quantified using the Qubit fluorometer (Thermo Fisher). *Adaptive Sampling:* DNA was prepared for sequencing using the Ligation Sequencing Kit (Oxford Nanopore Technologies, catalog no. SQK-LSK109) and sequenced using the GridION sequencer (ONT) with readfish^1^ integration (minKNOW 20.10.6) or using MinKNOW Adaptive Sampling^2^ (minKNOW 19.16.6, guppy 3.4.5) with a target region of interest defined as chrX:141000000-156000000 in the GRCh38 reference. *Cas9 Sequencing:* Guide RNAs were designed and ordered using the Custom Alt-R® CRISPR-Cas9 guide RNA design tool (Integrated DNA Technologies) with a 1 kb reference input fasta target region from the human GRCh38 genome. Cas9 sequencing libraries were prepared using the Cas9 Sequencing Kit (Oxford Nanopore Technologies, catalog no. SQK-CS9109) with modifications as described^3^. Ultra-high molecular weight (UHMW) Cas9 sequencing libraries were prepared with the following additional modifications: adapter-ligated libraries were purified via Nanobind disk (Circulomics) with precipitation in NAF10 buffer (Circulomics); Nanobind disks were washed three times with magnetic separation in Long Fragment Buffer (Oxford Nanopore Technologies, catalog no. LFB); final elutions were carried out at room temperature overnight with 60 µL or 120 µL Elution Buffer for MinION or PromethION libraries, respectively. Samples were sequenced using the MinION, GridION, or PromethION sequencer (Oxford Nanopore Technologies) using R9.4.1 flow cells. Final libraries were combined with 30 µL or 120 µL of Sequencing Buffer for MinION or PromethION flow cells, respectively, and equilibrated for 30 minutes at room temperature prior to loading with a wide-bore pipette tip. Flow cells were flushed and reloaded as needed using the Flow Cell Wash Kit (Oxford Nanopore Technologies, catalog no. EXP-WSH004). *Analysis:* Due to each genome in this family containing a different mecp2 locus with different expected ploidy, each genome was examined using a different method. Adaptive Sampling reads for BAB3121 (flowcell: FAO74863) were aligned to GRCh38 with minimap2 (version 2.17), and SNVs were detected using the medaka_variant wrapper (version 1.0.3). Cas9 targeted reads for BAB3121 (flow cell: FAN49258) were analyzed using an identical workflow. BAB3114 was sequenced using either a pair of Cas9 targets that flank the region of interest (flowcell: PAG08429) or a single Cas9 target next to the region of interest (flowcell: FAO31820). Flanking target Cas9 reads were aligned with minimap2, and SNVs were detected with medaka using medaka’s default diploid method. Single-target Cas9 reads (flowcell: FAO31820) were aligned with minimap2 and then inspected manually for ultra-long reads.

BAB3115 was sequenced using two separate single-target Cas9 guide RNAs (flowcells: FAN40573 and PAG08038) in order to preferentially enrich for either the two original copies of *FLNA*, or the additional copy created in the rearrangement, which we call *FLNA*’. Reads for the two original *FLNA* copies were enriched by targeting five Cas9 guide RNAs to the *RPL10* gene, which is only found adjacent to these two copies of *FLNA*. Reads were aligned with minimap2, and SNVs were called using medaka_variant. To enrich for *FLNA*’ reads, three Cas9 guide RNAs targeting *TKTL*-1 were used, which flanks *FLNA*’ on both sides but is only on one side of the original *FLNA* gene copies. Reads were filtered to include reads producing either primary alignments or any supplementary alignments ending within 50 bp of the telomeric end of the *FLNA* flanking repeat (chrX:154384868-154396222 in GRCh38). All alignments from these reads were then subjected to variant calling with medaka before final analysis validation was performed.

### Oxford Nanopore (Minion)

In house nanopore sequencing used a minion R.10.4.1 flow cell, with the V14 ligation sequencing kit (LSK114) following the manufacturer’s directions with modifications. DNA was sheared to a N50 of 10 kb using a g-tube (Covaris), 2 µg of DNA was sheared by centrifugation at 5500 rpm in an Eppendorf 5424r centrifuge two times for one minute each. Shearing was confirmed by visualization on a 1% agarose gel. DNA ends were repaired using the NEBNext FFPE DNA Repair kit (NEB cat# E7180S) following the manufactures directions. DNA was purified by AMPure magnetic beads. Sequencing adapters ligation was carried out using NEB Quick Ligase (NEB cE7180S) and Oxford Nanpore’s ligation buffer. Following purification with AMPure magnetic beads. Fifteen femtomoles of library were loaded onto the R.10.4.1 flow cell following priming. Post run base calling used guppy 6.0.1. Reads were mapped with minimap2 to the hg19 reference genome.

### Read-Depth Plotting of Short-Read GS via VizCNV Platform

The depth of sequencing coverage was computed using mosdepth (version 0.3.4) [61] and subsequently visualized using our custom visualization tool, VizCNV (https://github.com/BCM-Lupskilab/VizCNV) [62]. This tool enables the plotting of normalized read depth for the individuals’ sequencing data, which facilitates manual assessment of CNVs exceeding 3 kilobases in size as well as the determination of B-allele frequency for a given genomic range.

### Strand-Seq

#### Strand-Seq data generation and data processing

Strand-Seq data were generated at the European Molecular Biology Laboratory using a modification of the the OP-Strand-Seq library preparation protocol [63]. Briefly, Lymphoblastoid cell lines derived from two patients (BAB3114, BAB14547) were first cultured in RPMI media and subjected to 40μM BrdU treatment for 18h and 24h. The cells were then lysed to release the nuclei and nuclei were digested with RNase and MNase as described in the original protocol. Cells were fixed with formaldehyde, and the crosslinked nuclei were stained with Hoechst to reveal the population of cells that had incorporated BrdU for a single cell division. The population of once-divided cell nuclei was used to sort single nuclei into 96 well plates using fluorescence-activated cell sorting (FACS). The individual nuclei were processed to produce a sequencing library using a robotic liquid handler. In brief, nuclei were first de-crosslinked, protease-digested, fragmented DNA ends “polished” and Illumina adapters ligated as described in the original OP protocol with the necessary volumetric adjustments according to the starting volume. A necessary deviation from the protocol in our hands was to introduce a bead-based cleanup after adapter ligation to remove adapter dimers prior to PCR amplification. This cleanup was done at a 0.8x bead: DNA ratio.

The adapter dimer free DNA was then exposed to Hoechst and UV light to ablate the BrdU-substituted strands. Finally, the libraries were PCR amplified for 15x cycles and simultaneously barcoded using a dual indexing strategy with iTru adapters. Amplified libraries were again subjected to a bead-based cleanup at a 0.8x bead ratio and pooled for size selection. Final, size-selected libraries were subjected to deep sequencing on the Illumina NextSeq500 platform (MID-mode, 75 bp paired-end protocol). The resulting raw read files were aligned to the GRCh38 reference assembly (GCA_000001405.15) using BWA aligner (version 0.7.17). Low-quality libraries were automatically flagged using ASHLEYS (version 1.0) [64], resulting in 41/96 (43%) and 52/96 (54%) viable, high-quality single-cell libraries for BAB14547 and BAB3114, respectively.

#### Strand-Seq data analysis

To detect SV breakpoint candidates, we initially flagged genomic regions which displayed a switch in read directionality, suggestive of inversion- or inverted duplication breakpoints, using the breakpointR tool (version 0.99.0) with default settings [65]. Using this procedure, we generated breakpoint estimates from each individual cell, with confidence intervals between 10 kbp (default minimum resolution) and >100 kbp for poorly covered regions. Under the assumption of clonality, we merged breakpoint estimates of cells from the same sample and extracted ‘peak’ regions in which at least 75% of cells predicted a breakpoint, yielding high-confidence consensus breakpoint regions of typically 10 kbp size. Genotypes for all regions were subsequently obtained using the ArbiGent tool [66], which estimates regional genotypes based on a directionality-specific read depth model and integrates this information across cells. Lastly, to confirm the obtained breakpoints and genotypes over long-range haplotype stretches visually, we generated a pseudo-bulk data track for each sample, which is conceptually similar to Strand-Seq based ‘composite files’ described previously [67]. In these tracks, we combined the reads from all cells and synchronised their read directionality in a way that ‘reference’ and ‘inverse’ orientation are encoded by reads mapping on the ‘W’ and ‘C’ strands, respectively. All previously obtained breakpoint regions and genotypes could be confirmed after visualising this pseudo-bulk track in the UCSC browser.

## Abbreviations

DUP-TRP/INV-DUP: duplication-triplication/inverted-duplication; CGR: complex genomic rearrangement; LCR: low copy repeat; TS: template switching; CNV: copy number variant; MDS: *MECP2* duplication syndrome; AOH: absence of heterozygosity; SV: structural variant; SNV: single nucleotide variant; NHEJ: non-homologous end-joining; BIR: break-induced replication; UHMW: ultra-high molecular weight; CCS: circular consensus sequencing; MMBIR: microhomology-mediated break-induced replication, PSV: paralogous sequence variant

## Declarations

### Ethics Approval and Consent to Participate

All study participants were consented according to the Institutional Review Board for Human Subject Research at Baylor College of Medicine approved protocols: H-29697, H-20268, and H- 47127 or Pacific Northwest Research Institute WIRB #20202158.

### Consent for Publication

All study participants have consented for publication.

### Availability of data and materials

Microarray data generated in previous studies [4,14] are available through the gene expression omnibus (GEO) under accessions GSE49440 and GSE49446; new microarray data within this study is available under accession XXX. Oxford nanopore datasets are available within SRA BioProject ID PRJNA953021 and dbGaP phs002999.v1.p1 with additional genomic data available within BioProject ID XXX.

### Authors contributions

CMG performed laboratory experiments, interpreted and analyzed data, and wrote the manuscript. JB, MG, KP, JMF and WH performed laboratory experiments as well as analyzed and interpreted data. JOK SNJ, DMM, RAG, AL, JRL interpreted and/or analyzed data. MYL, HD, MGM, MM, LFP, MP, EH, SJ, FJS performed bioinformatics analyses and interpretation. DP provided patient samples as well as clinical interpretation. CMBC conceptualized the study, analyzed and interpreted the data as well as contributed to writing the manuscript. All authors have read, edited and approved the final manuscript.

---

Additional File 1: Supplemental Tables and Legends

**Supplemental Table 1:** Genomic location and size of each genomic fragment in the DUP-TRP/INV-DUP structure for each individual within the cohort. All coordinates are based on array positions in Hg19 genome reference.

**Supplemental Table 2:** LCR summary table including the genomic location (Hg19) as well as nucleotide size (bp), distance between the inverted pair and similarity score (%) each of the 4 pairs identified in this cohort.

**Supplemental Table 3:** Sequencing methodology that was performed on each sample within this cohort.

**Additional File 2:** ArrayCGH showing the *MECP2* critical region, nucleotide-level resolution of junction 2 as well as resolved haplotype and OGM data for the region (where applicable) for each sample.

Additional File 3: Supplemental Figures and Legends

**Supplemental** Figure 1: Structural variant haplotype possibilities within a DUP-TRP/INV-DUP event during and interchromosmal and intrachromosomal template switch between inverted LCRs.

**Supplemental** Figure 2: Possible pathogenic effects of a DUP-TRP/INV-DUP event include gene dosage (A) gene interruption (B) as characterized by a disruption within the *DMD* gene [22] or gene fusion (C) events.

**Supplemental** Figure 3: Three pairs of inverted LCRs at Xq28 (43202a/43202a; 43221a/43221b (K1/K2); 43231a/43231b (L1/L2)) within the *MECP2* locus with the fourth pair being located at Xq22.1 (37696a/ 37696b). The corresponding location in the UCSC genome browser is shown.

**Additional File 4:** CRISPR-Cas9 targeted ONT data for BAB3114 family.

**Additional File 5:** StrandSeq data for samples BAB3114 and BAB14547.

