## Additional File 2 for "Break-induced replication underlies formation of inverted triplications and generates unexpected diversity in haplotype structures"

**Additional File 2:** ArrayCGH showing the *MECP2* critical region, nucleotide-level resolution of junction 2 as well as resolved haplotype and OGM data for the region (where applicable) for each sample.

### BAB2727

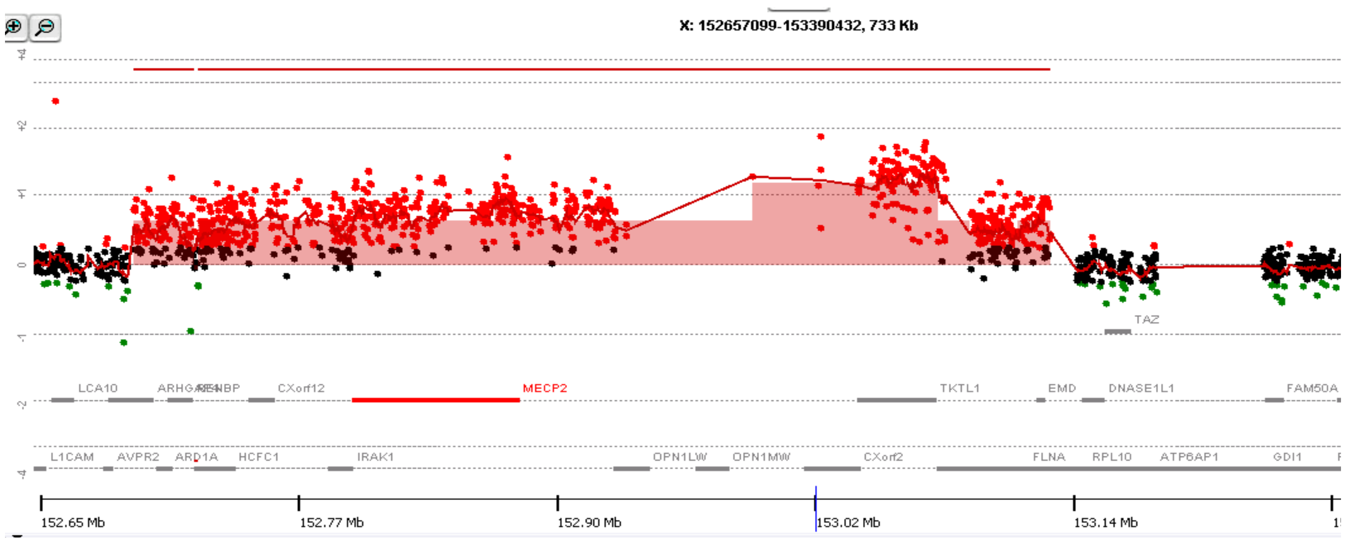


#### Breakpoint Junction 2:

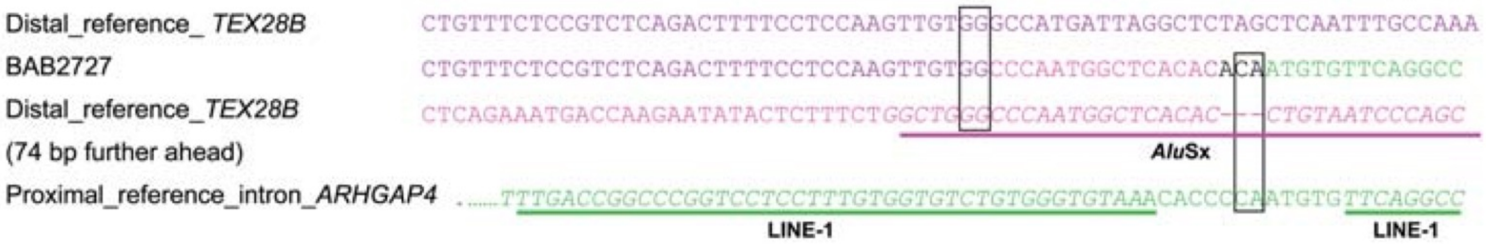


***Adapted From:***

Carvalho, C. M. B., Zhang, F., Liu, P., Patel, A., Sahoo, T., Bacino, C. A., Shaw, C., Peacock, S., Pursley, A., Tavyev, Y. J., Ramocki, M. B., Nawara, M., Obersztyn, E., Vianna-Morgante, A. M., Stankiewicz, P., Zoghbi, H. Y., Cheung, S. W., & Lupski, J. R. (2009). Complex rearrangements in patients with duplications of MECP2 can occur by fork stalling and template switching. *Human Molecular Genetics*, *18*(12), 2188–2203.

**Haplotype Structure: *Not Available***

### BAB2769

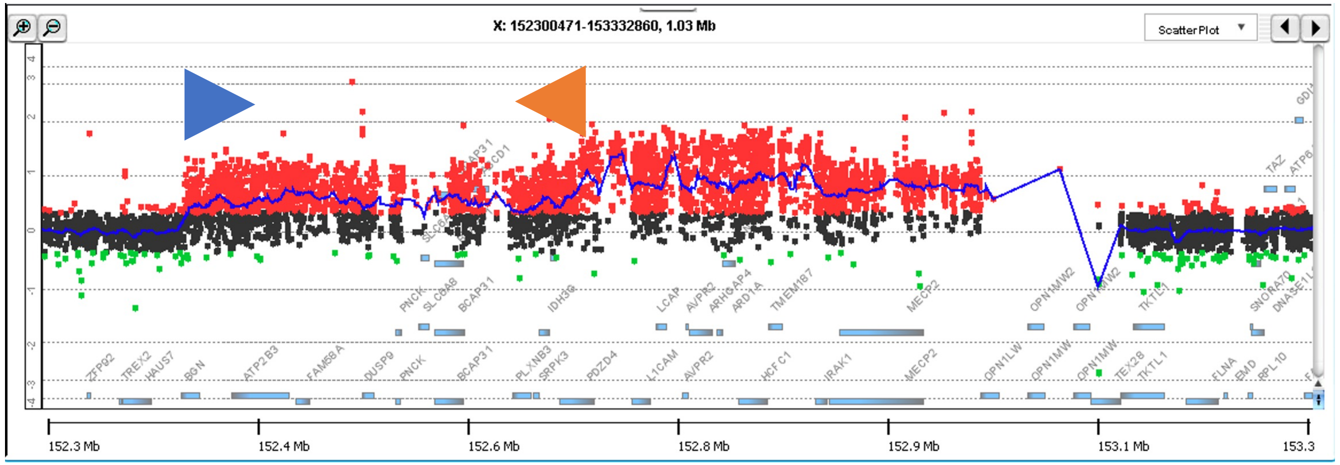


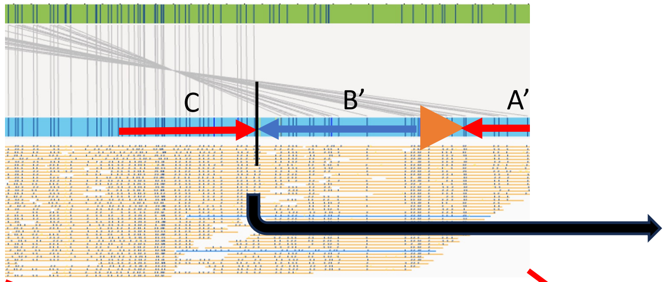


#### Junction 2

DUPd (+) AACCCCCGCAGGAAAGCCTGTGGGATCACTTCACACCCTGTGACCTTAACTTCACACCCTGTGACCTTAAT

BAB2769 AACCCCCGCTGGAGCCGGGCGCGGGGCCTAGGGCTTTCAGC-----------------------AGGCCAG

TRPd (–) ACACCTGGCTGGAGCCGGGCGCGGGGCCTAGGGCTTTCAGCCTGTGTGAGTGGGTCCTGCCAGCAGGCCAG

#### Junction 1

TRPp (–) TCTGTGGGCGGGAGA.........GGCTTTCTCAGGG

BAB2769 TCTGTGGGCGGGAGA.........GGCTTTCCCAGGG

DUPp (+) TCTGTGGGCCGGAGA.........GGCTTTCCCAGGG

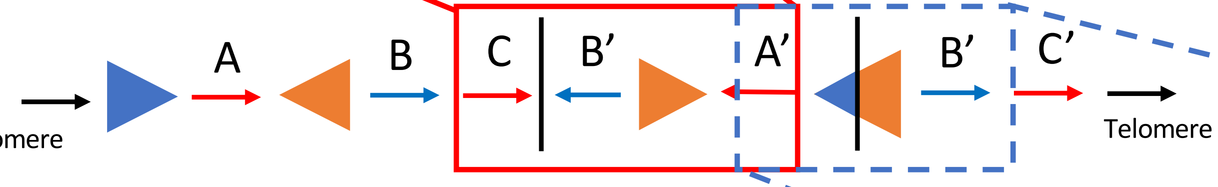


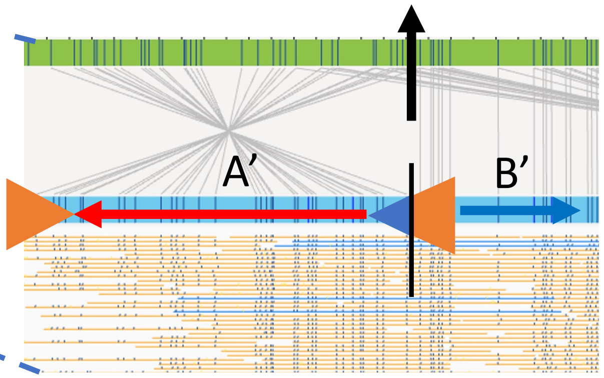


#### Haplotype Structure 3

***Adapted From:***
Carvalho, C. M. B., Ramocki, M. B., Pehlivan, D., Franco, L. M., Gonzaga-Jauregui, C., Fang, P., McCall, A., Pivnick, E. K., Hines-Dowell, S., Seaver, L. H., Friehling, L., Lee, S., Smith, R., Del Gaudio, D., Withers, M., Liu, P., Cheung, S. W., Belmont, J. W., Zoghbi, H. Y., … Lupski, J. R. (2011). Inverted genomic segments and complex triplication rearrangements are mediated by inverted repeats in the human genome. *Nature Genetics*, *43*(11), 1074–1081.

### BAB2772

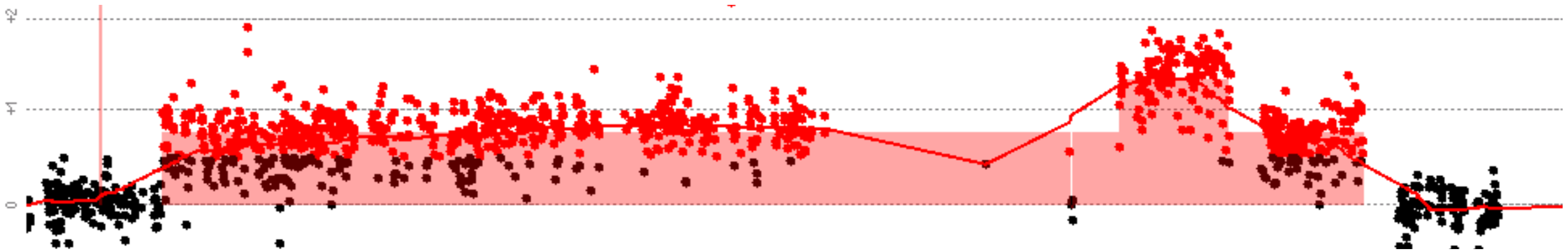

#### Breakpoint Junction 2:

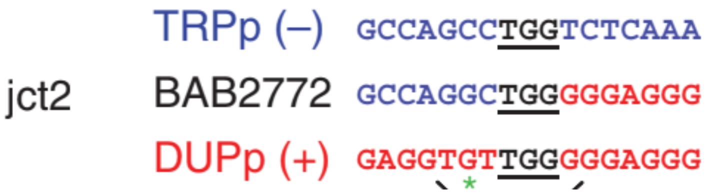


***Adapted From:***
Carvalho, C. M. B., Ramocki, M. B., Pehlivan, D., Franco, L. M., Gonzaga-Jauregui, C., Fang, P., McCall, A., Pivnick, E. K., Hines-Dowell, S., Seaver, L. H., Friehling, L., Lee, S., Smith, R., Del Gaudio, D., Withers, M., Liu, P., Cheung, S. W., Belmont, J. W., Zoghbi, H. Y., … Lupski, J. R. (2011). Inverted genomic segments and complex triplication rearrangements are mediated by inverted repeats in the human genome. *Nature Genetics*, *43*(11), 1074–1081.

#### Haplotype Structure 3:

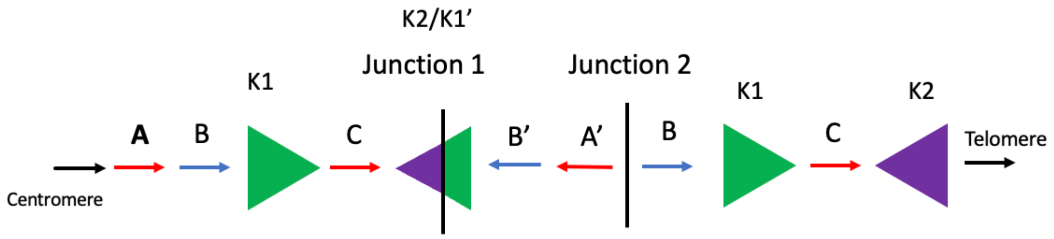


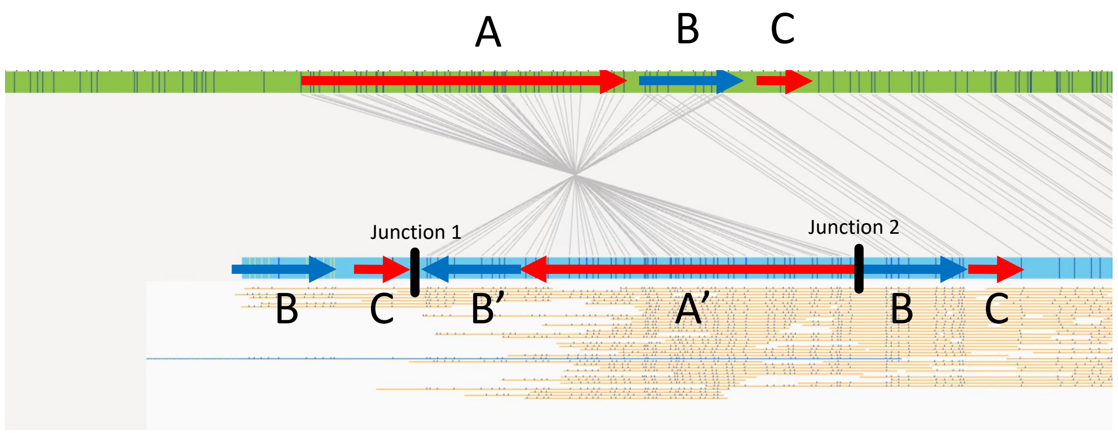

### BAB2796

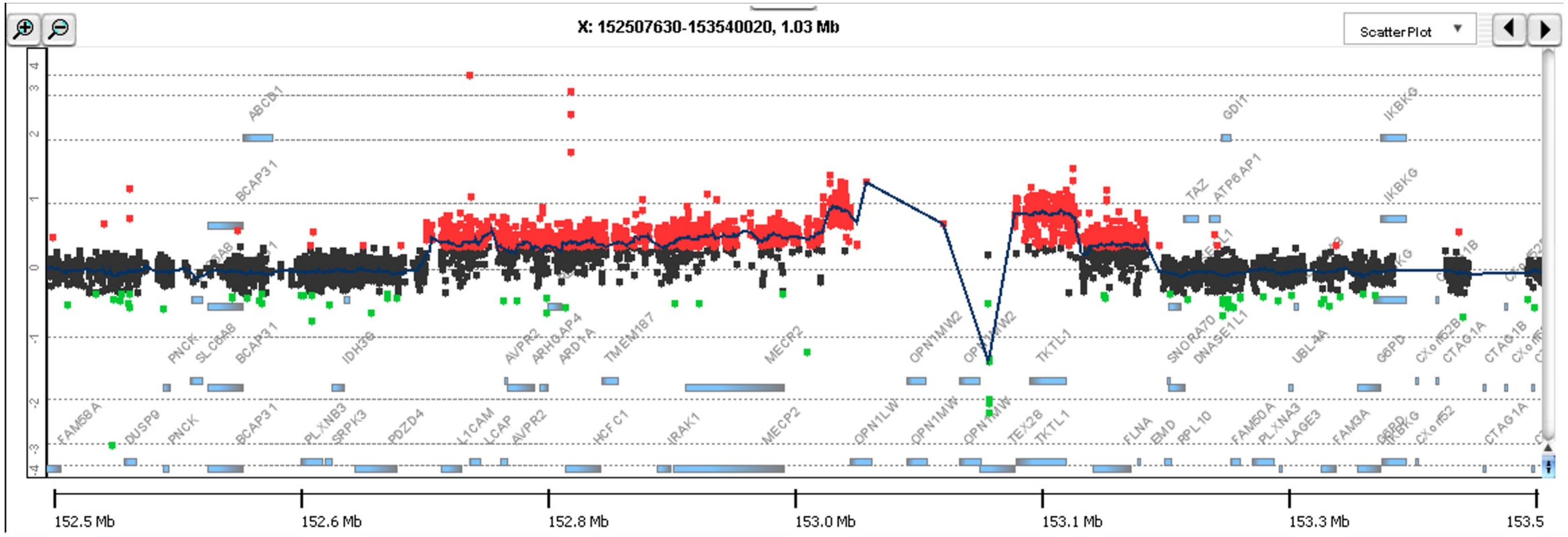


#### Breakpoint Junction 2:

| | | |
|---|---|---|
| | TRPp (–) | TCTCTTG--CCTCAGC |
| jct2 | BAB2796/BAB2980 | TCTCTTGAATTATAAC |
| | DUPp (+) | CTAATTCTTTTATAAC |

***Adapted From:***

Carvalho, C. M. B., Ramocki, M. B., Pehlivan, D., Franco, L. M., Gonzaga-Jauregui, C., Fang, P., McCall, A., Pivnick, E. K., Hines-Dowell, S., Seaver, L. H., Friehling, L., Lee, S., Smith, R., Del Gaudio, D., Withers, M., Liu, P., Cheung, S. W., Belmont, J. W., Zoghbi, H. Y., … Lupski, J. R. (2011). Inverted genomic segments and complex triplication rearrangements are mediated by inverted repeats in the human genome. *Nature Genetics*, *43*(11), 1074–1081.

#### Haplotype Structure 2

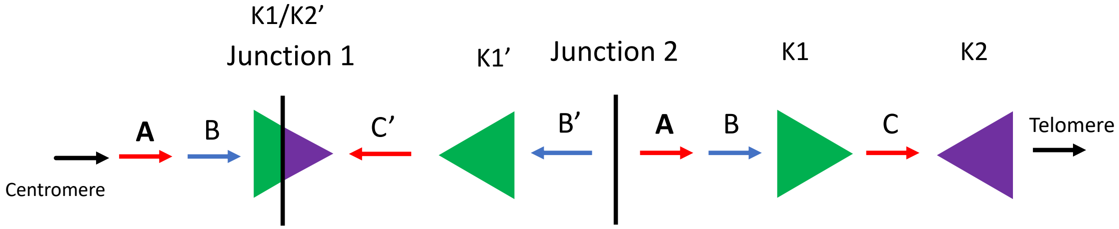


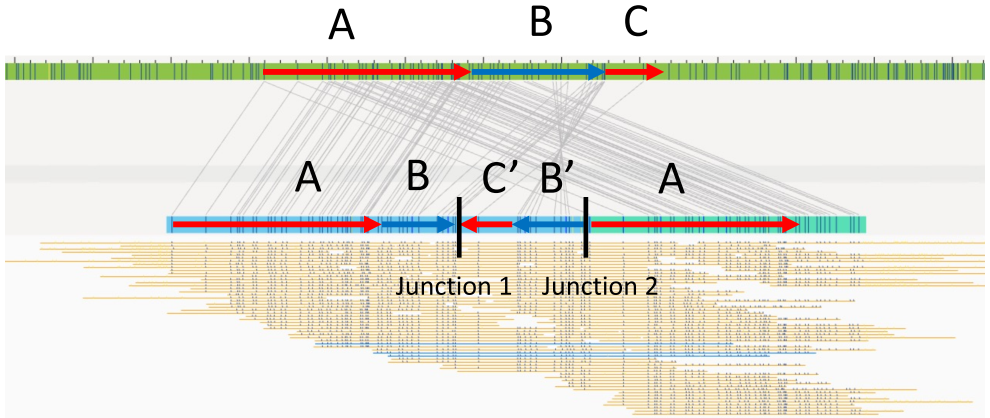

### BAB2797

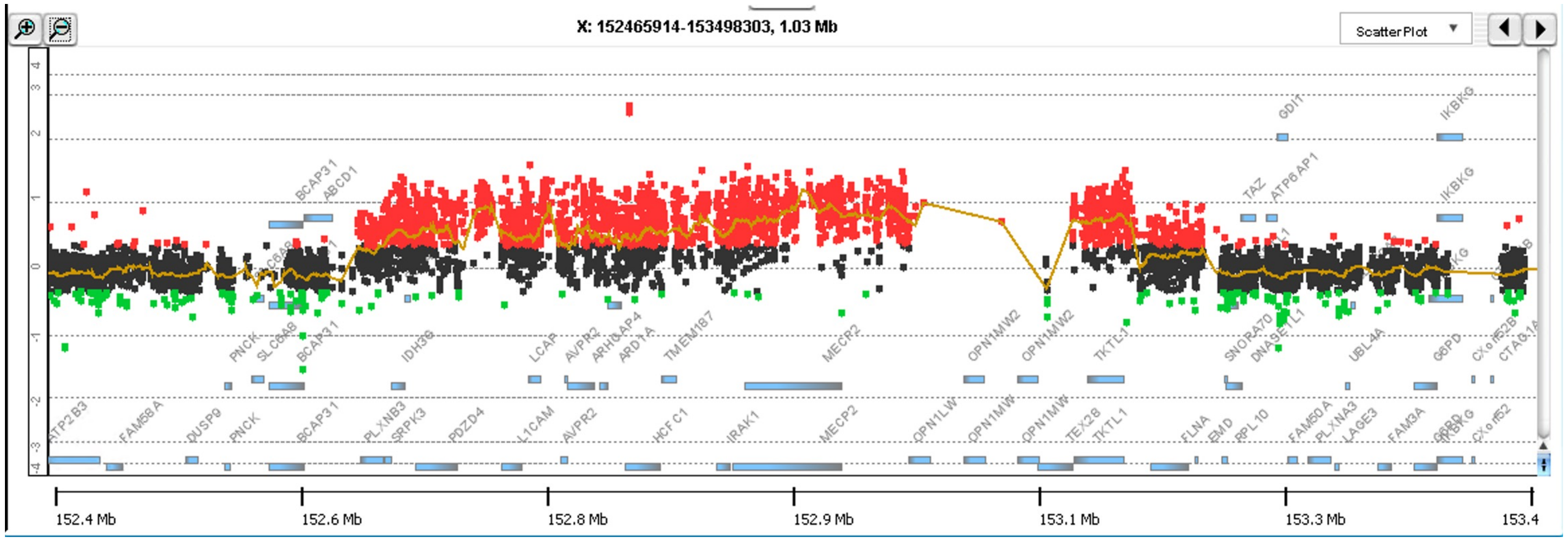


**Breakpoint Junction 2:**

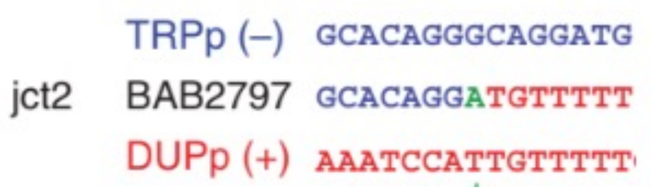


***Adapted from:***

Carvalho, C. M. B., Ramocki, M. B., Pehlivan, D., Franco, L. M., Gonzaga-Jauregui, C., Fang, P., McCall, A., Pivnick, E. K., Hines-Dowell, S., Seaver, L. H., Friehling, L., Lee, S., Smith, R., Del Gaudio, D., Withers, M., Liu, P., Cheung, S. W., Belmont, J. W., Zoghbi, H. Y., … Lupski, J. R. (2011). Inverted genomic segments and complex triplication rearrangements are mediated by inverted repeats in the human genome. *Nature Genetics*, *43*(11), 1074–1081.

**Haplotype Structure:** ***Not Available***

### BAB2801

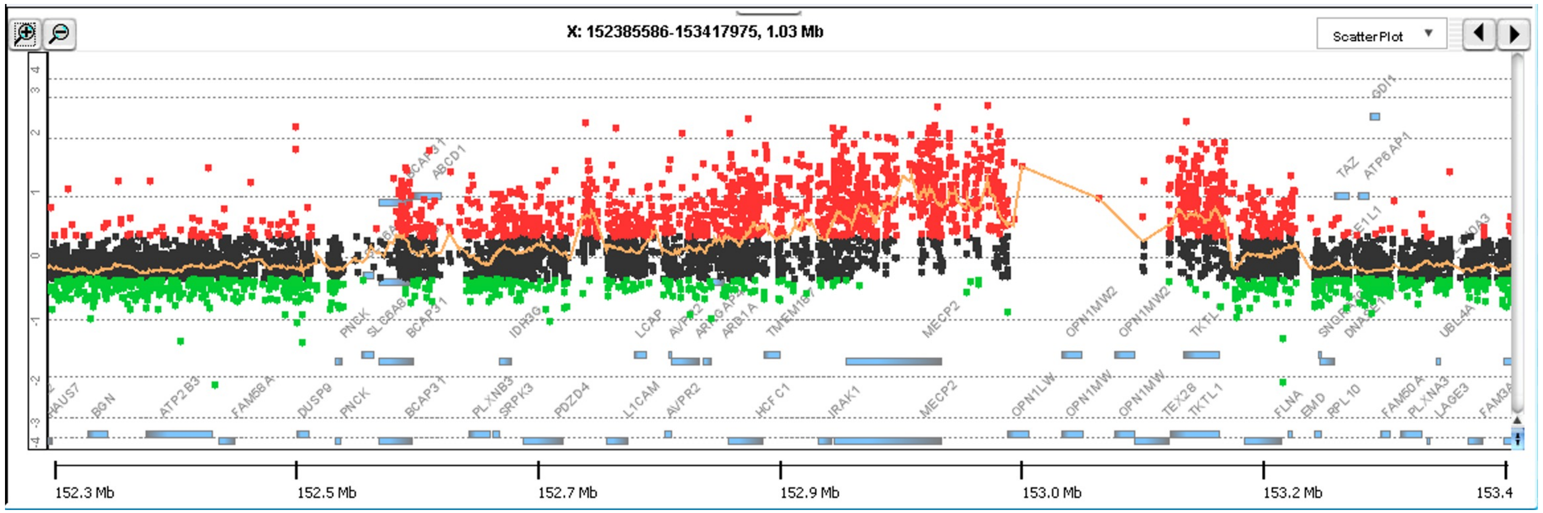


**Breakpoint Junction 2:**

chrX:152955672

chrX:152955637(+) TACTGTAGCCCCTCTTCAACACACTCAACCCACCCCCTCAAGACTCCACCTGGGCCTGAGTCAGTGGCC

Junction 2 ATAGTTTCACTGTGTTCCACGCATTCATCCCTCCCCCTCAAGACTCCACCTGGGCCTGAGTCAGTGGCC

chrX:153198729(-) ATAGTTTCACTGTGTTCCACGCATTCATCCCTCCCTCTCCTTAACCCGTGGCAACCACTGATTTTTACT

chrX:153198702

**Haplotype Structure 4**

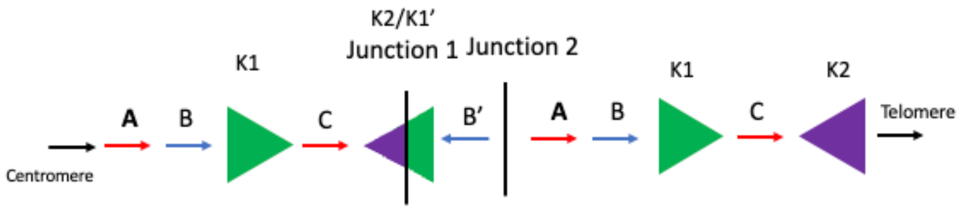


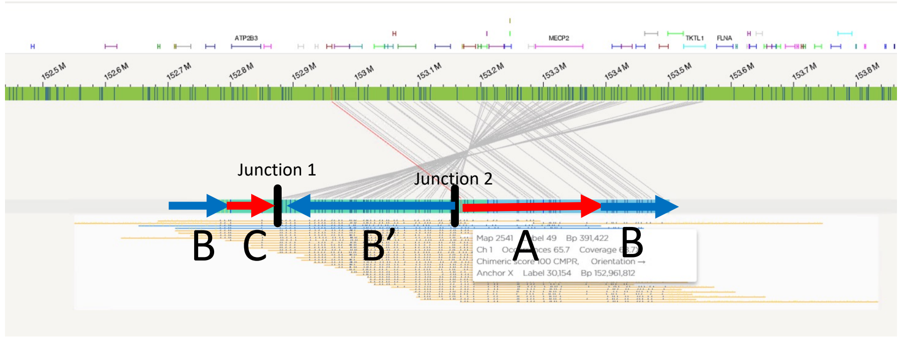

### BAB2805

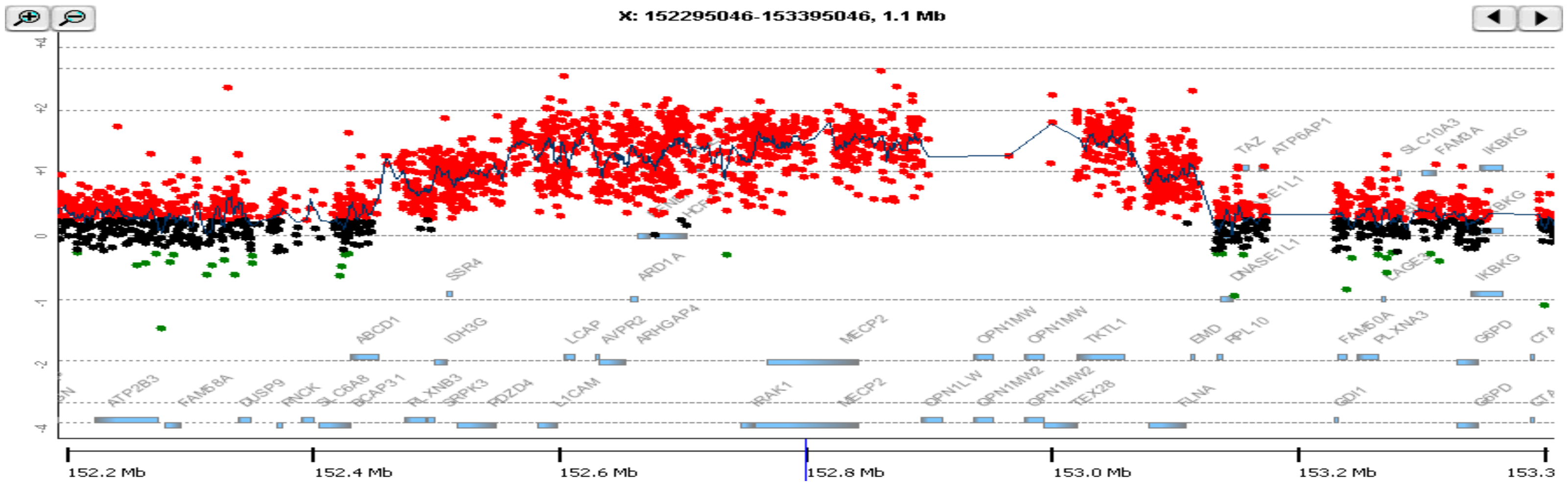


**Breakpoint Junction 2:**

| | | |
|---|---|---|
| | TRPp (–) | AACAGACAAAAAAAGGCCGGGC |
| jct2 | BAB2805 | AACAGACAAAAAAAGATG-AAA |
| | DUPp (+) | ATAAAATAAAAAAATATGGAAA |

***Adapted from:***

Carvalho, C. M. B., Ramocki, M. B., Pehlivan, D., Franco, L. M., Gonzaga-Jauregui, C., Fang, P., McCall, A., Pivnick, E. K., Hines-Dowell, S., Seaver, L. H., Friehling, L., Lee, S., Smith, R., Del Gaudio, D., Withers, M., Liu, P., Cheung, S. W., Belmont, J. W., Zoghbi, H. Y., … Lupski, J. R. (2011). Inverted genomic segments and complex triplication rearrangements are mediated by inverted repeats in the human genome. *Nature Genetics*, *43*(11), 1074–1081.

**Haplotype Structure: *Not Available***

### BAB3114

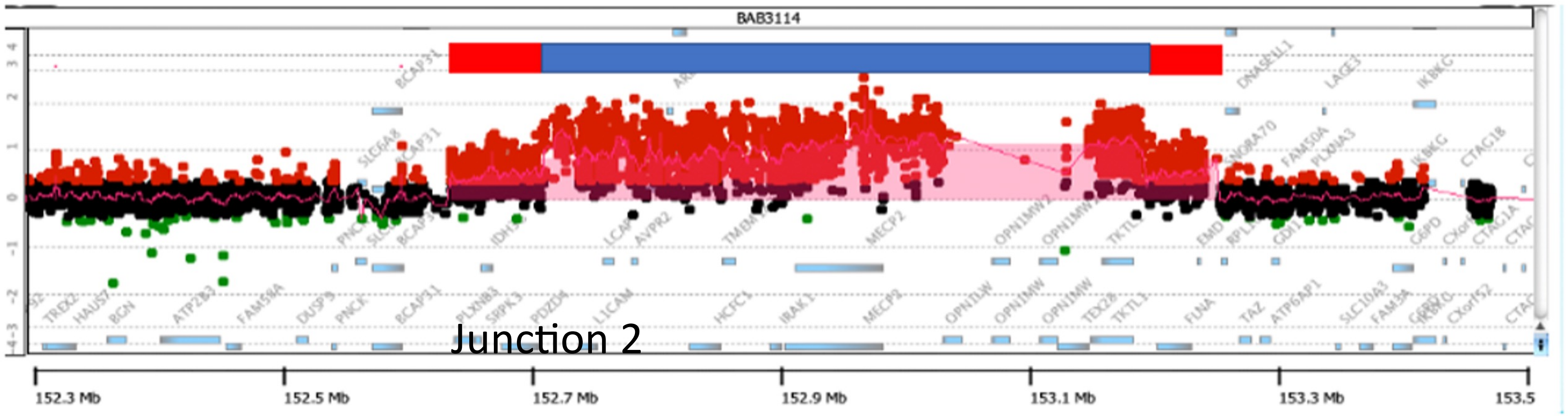


**Breakpoint Junction 2:**

```
                                          chrX:153024486
chrX:153024456(+) AACAAATGGTATAGGGGAAACTTGATATCCACATGCAAAAGAATAAAGTTGGAGGCTGGGCATGGTG
       Junction 2 CCATCCCCCGCTGCCCTTTTTCTCCTGGCCACATGCAAAAGAATAAAGTTGGAGGCTGGGCATGGTG
chrX:153096011(-) CCATCCCCCGCTGCCCTTTTTCTCCTGGCCGGGTGCGAGGGAGCATGCCCGCGCCTGGCCTCGGCCA
                                chrX:153095984
```

**Haplotype Structure 1**

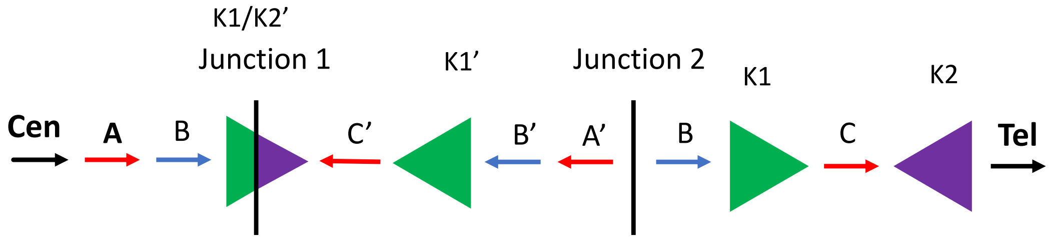


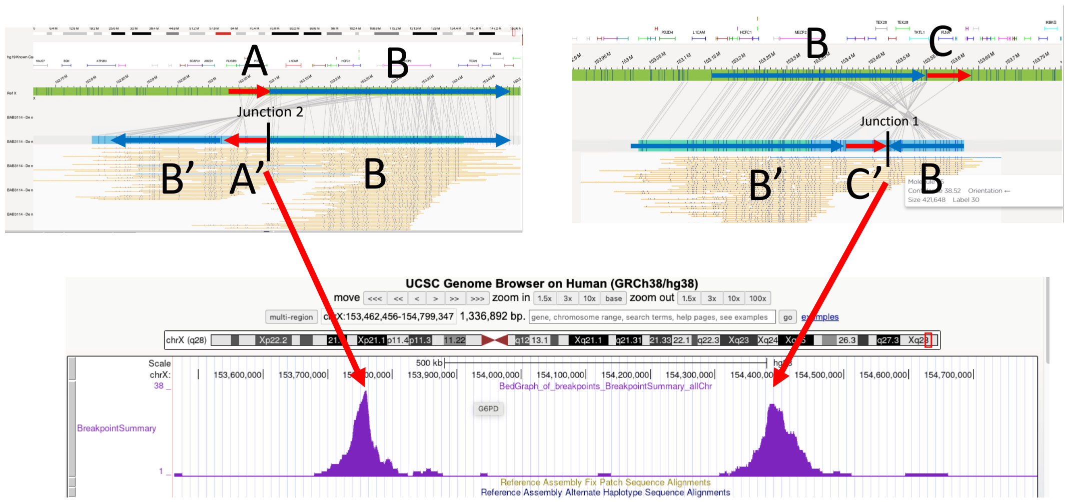

### BAB3147

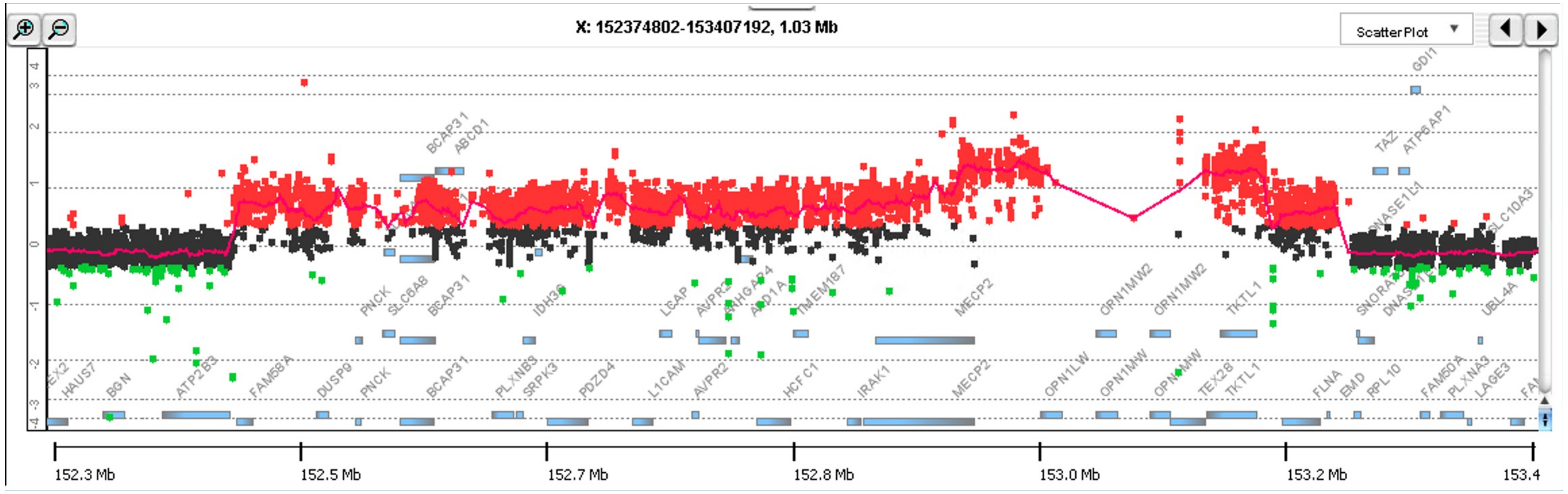


**Breakpoint Junction 2:**

```
                               chrX:152850819
chrX:152850786(+) GTTAGGTTCCAGAAGAAATATAGGCACGGAGGCTGTTTCTGGCTTGCAATTCTGTCACCTCAGAGT
       Junction 2 AGGTATAGCCCACTTAGTCAGTTGTGTCTAAGCTGTTTCTGGCTTGCAATTCTGTCACCTCAGAGT
chrX:153353826(-) AGGTATAGCCCACTTAGTCAGTTGTGTCTAAGCCACTTGATGAACCAGTCAATCCAGTCTTCACCT
                      chrX:153353796
```

**Haplotype Structure 6:**

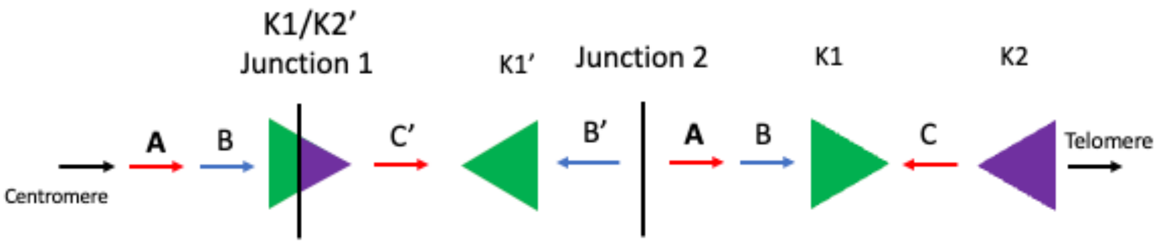


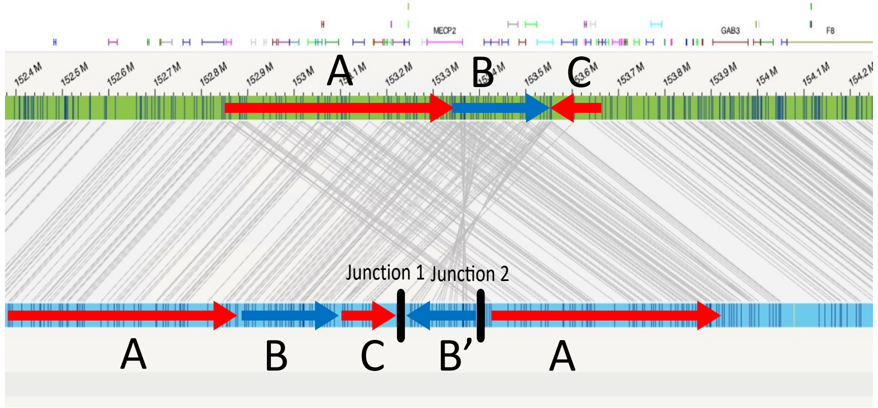

### BAB3216

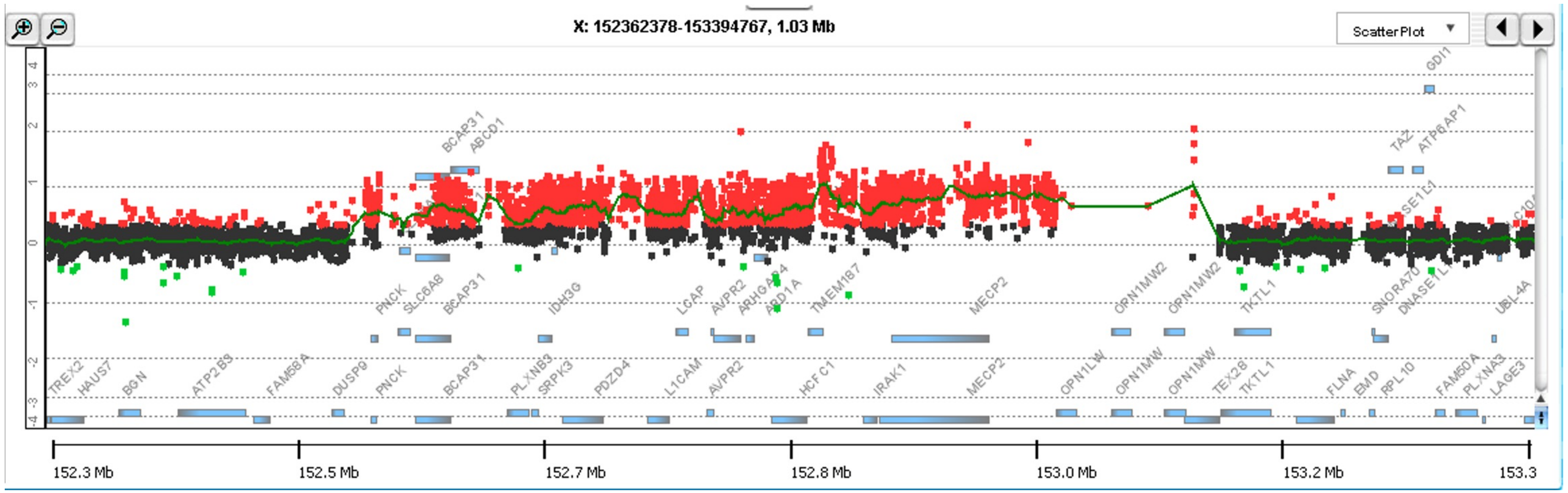


**Breakpoint Junction 2:**

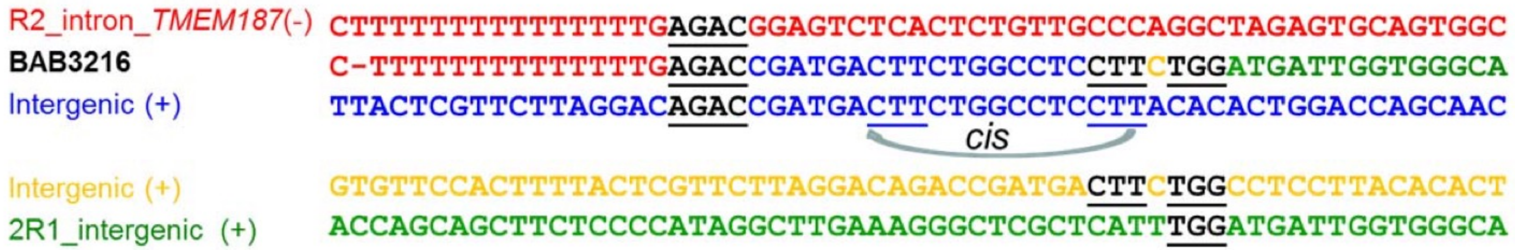


***Adapted from:***
Carvalho, C. M. B., Pehlivan, D., Ramocki, M. B., Fang, P., Alleva, B., Franco, L. M., Belmont, J. W., Hastings, P. J., & Lupski, J. R. (2013). Replicative mechanisms for CNV formation are error prone. *Nature Genetics*, *45*(11), 1319–1326.

**Haplotype Structure: *Not Available***

### BAB3255

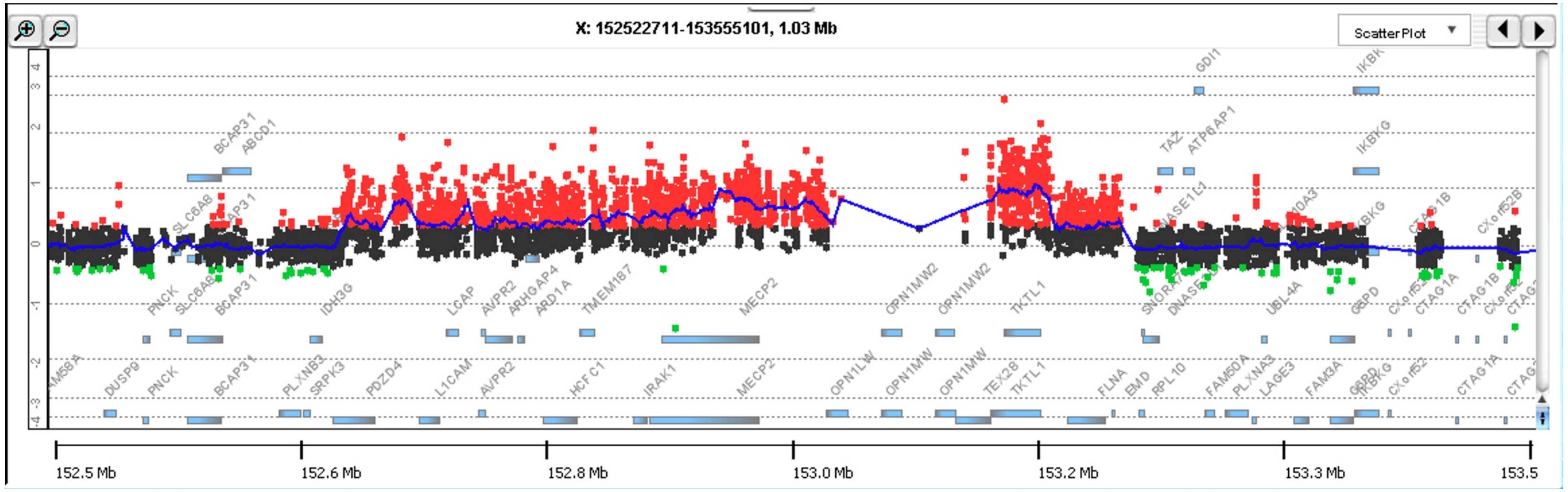


**Breakpoint Junction 2:**

chrX:153073620

chrX:153073584(+) CTCCCTCGGCCCTGGGGTAGATACAACGGGGCGCCTGCTGGAGCCCGGCCCAAGGCTAGAGGCCTGGAGGTTT

Junction 2 **TAAATTTCAACACGAGTTTTGGGGGGACATGTACCTGCTGGAGCCCGGCCCAAGGCTAGAGGCCTGGAGGTTT**

chrX:153433884(-) **TAAATTTCAACACGAGTTTTGGGGGGACATGTACC**CCATAGCAGTATGCTTAACTTTTTAAGAAAGAGGAAGG

chrX:153433851

**Haplotype Structure 3**

*Or*

**Haplotype Structure 1**

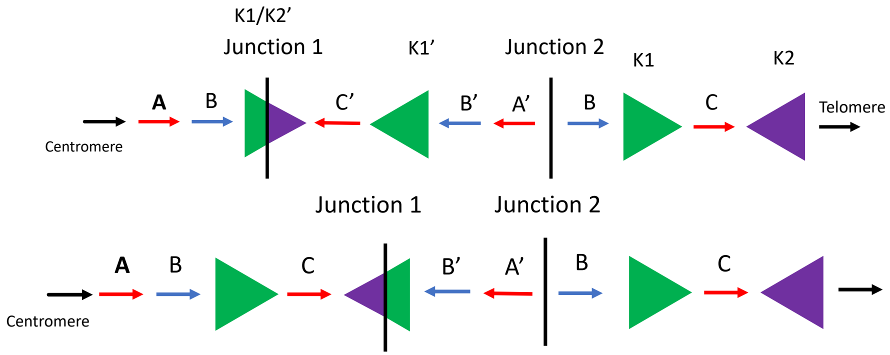


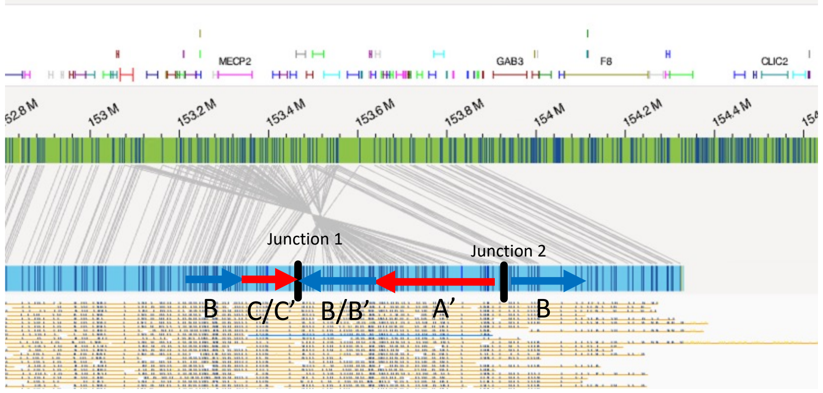

### BAB3274

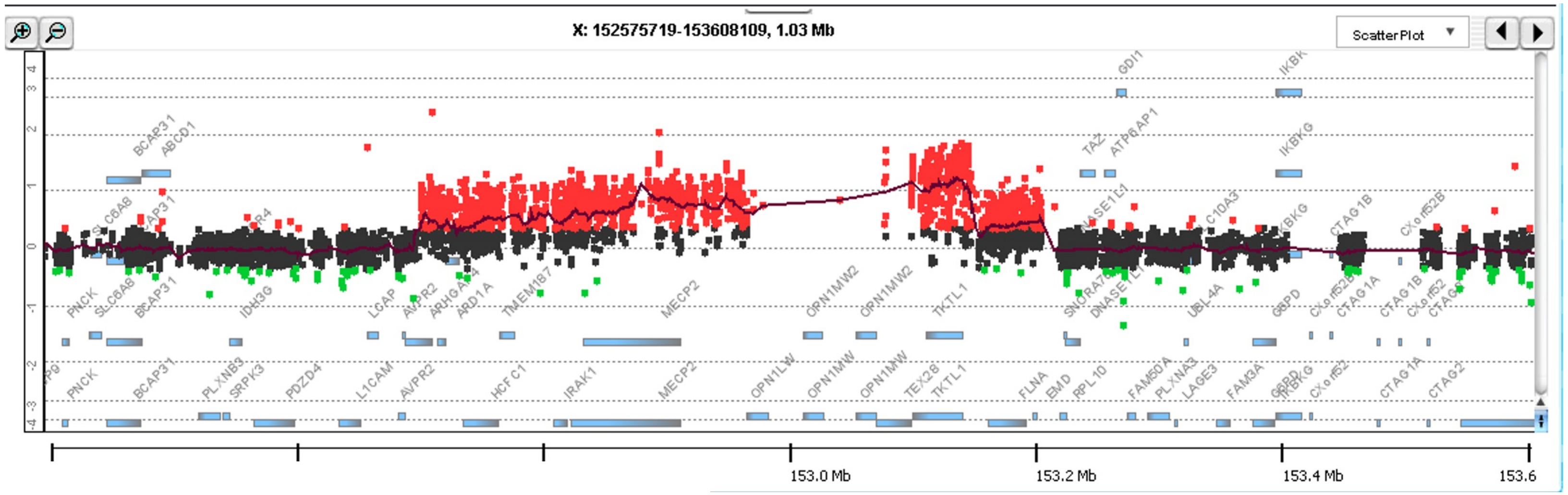


**Breakpoint Junction 2:**

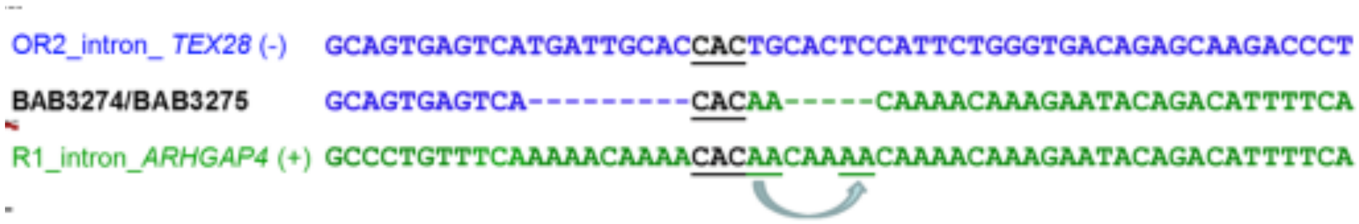


***Adapted from:***

Carvalho, C. M. B., Pehlivan, D., Ramocki, M. B., Fang, P., Alleva, B., Franco, L. M., Belmont, J. W., Hastings, P. J., & Lupski, J. R. (2013). Replicative mechanisms for CNV formation are error prone. *Nature Genetics*, *45*(11), 1319–1326.

**Haplotype Structure 3**

*Or*

**Haplotype Structure 1**

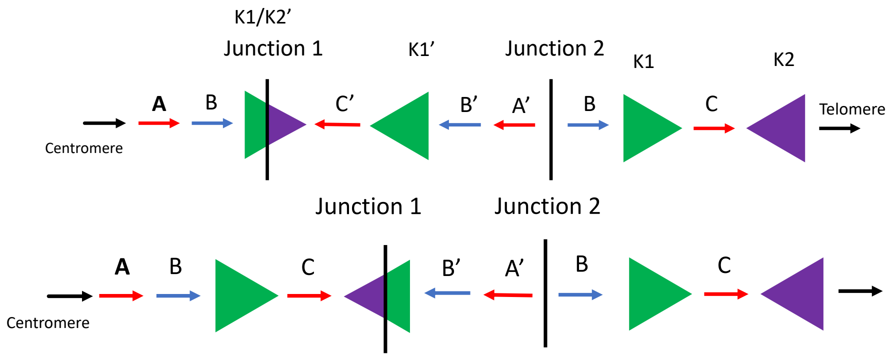


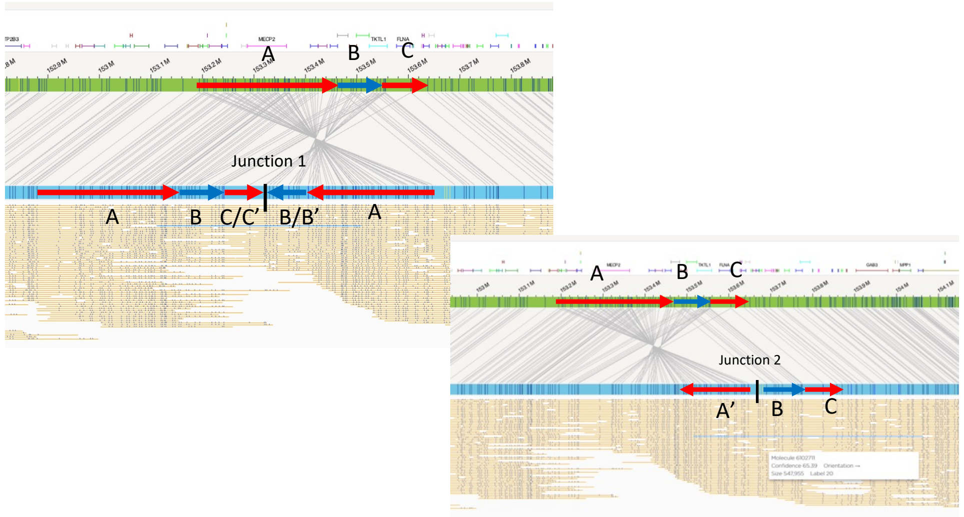

### BAB12566

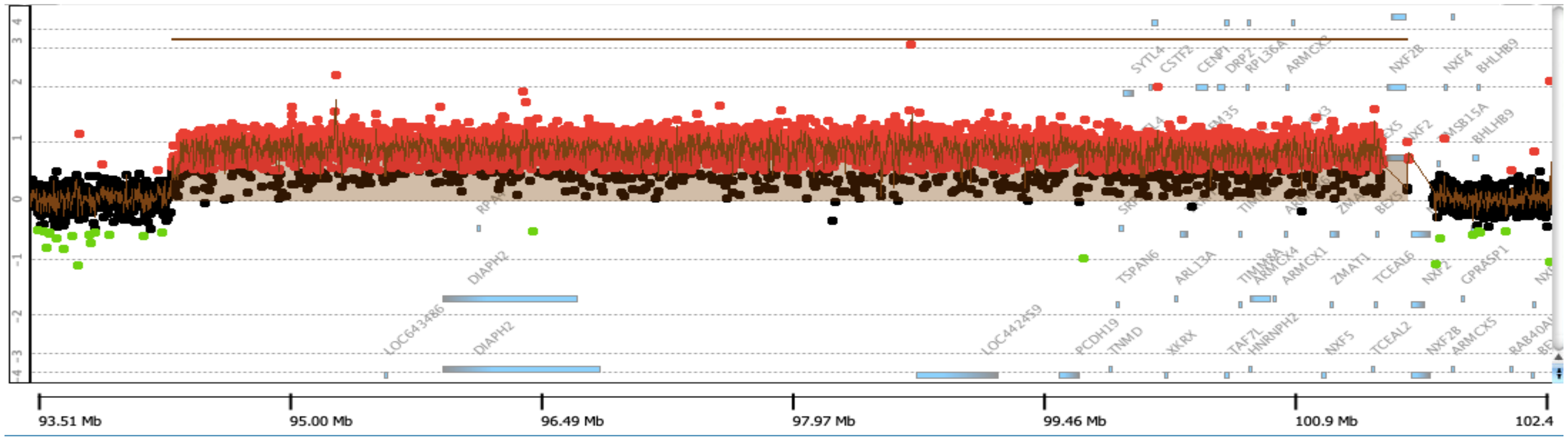


**Breakpoint Junction 2:**

```
                                                  chrX:94345302
chrX:94345271(+) AGAGAATTAAGGACAAAAACTGTATGATAATTTCAATTAATGCTGAACAAACATTTAATAGAATTCAGCTTC
       Junction 2 ATATTATCTATAGATAATAGATATTATCTATTTCAATTAATGCTGAACAAACATTTAATAGAATTCAGCTTC
ChrX:94394842(-) ATAATATAGATAACAGATATTATCTATCTATATATAATATAGATAACAGATATTATCTATCTATATATAATA
                                  ChrX:94394748
```

**Haplotype Structure 1:**

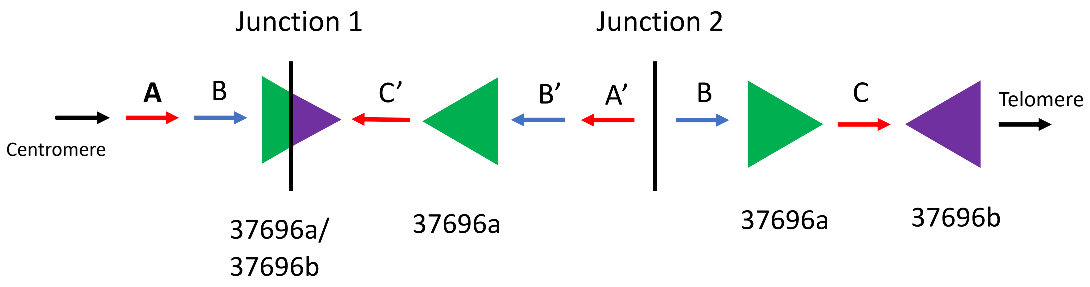


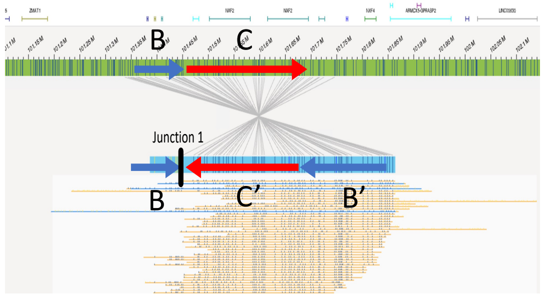


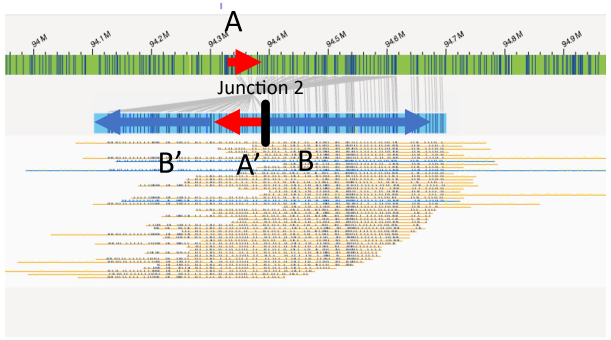

### BAB14392

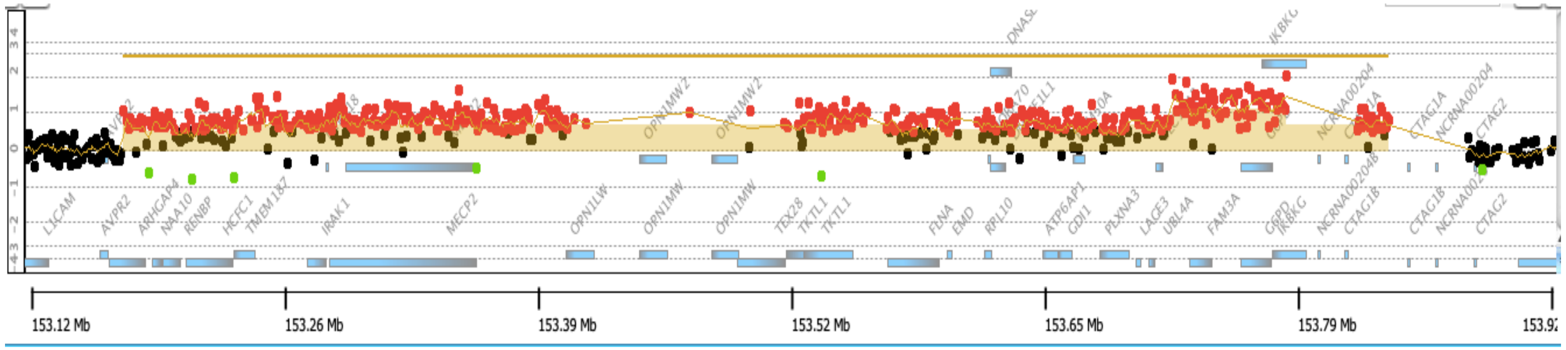


**Breakpoint Junction 2:**

chrX:153180047

chrX:153180013(+) CGGTGGCATGCGCCTGTAGTCCCAGCTACTTGGGAGGCTGAGGCAGGGGAATCGCTTGAACCCGGGAGGCGGA

Junction 2 CACCTCACCTGGCCTCTTTATTTATCAAATACTGAGGCTGAGGCAGGGGAATCGCTTGAACCCGGGAGGCGGA

chrX:153722563(-) CACCTCACCTGGCCTCTTTATTTATCAAATACTGTTTCTGGGCCAGGCACGGTGGCTTATGCCTGTAATCCCA

chrX:153722603

**Haplotype Structure:** ***Not Available***

### BAB14547

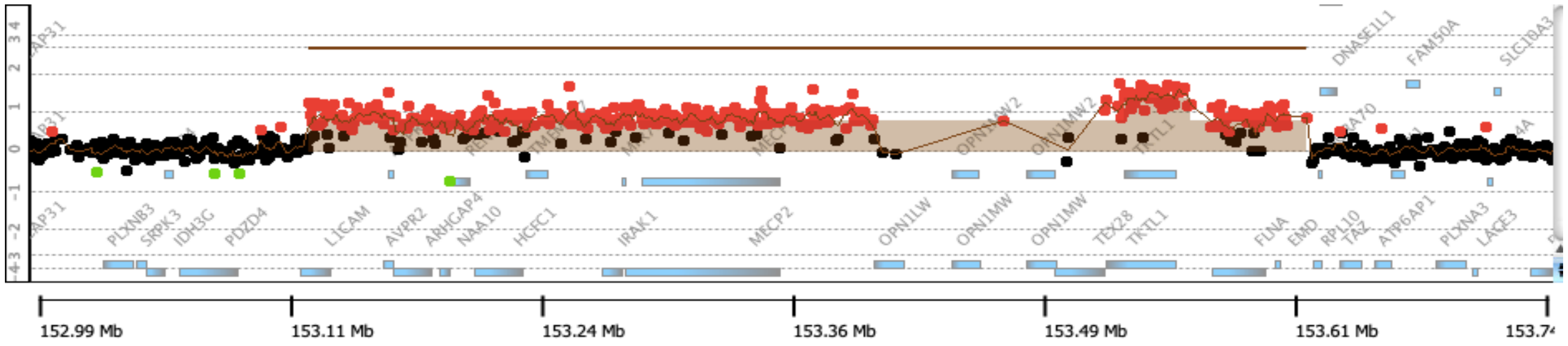


#### Breakpoint Junction 2:

chrX:153131087

chrX:153131049(+) TCAACCCAAGTCTTCCAGGAGAGCGGCTGGCAGGTGGCAAAGCCCCCTCACCATCCTGTCGCTTTACCTCAGTGATCA

Junction 2 CTCACTGCAACCTCTTGGGTTCAAGCGATTCTCTTGCCAAAGCCCCCTCACCATCCTGTCGCTTTACCTCAGTGATCA

chrX:153445956(-) CTCACTGCAACCTCTTGGGTTCAAGCGATTCTCTTGCCTCAGCCTCCCGAGTAGCTGGGATTACAGGCGCCCACCACA

chrX:153445920

#### Haplotype Structure 3:

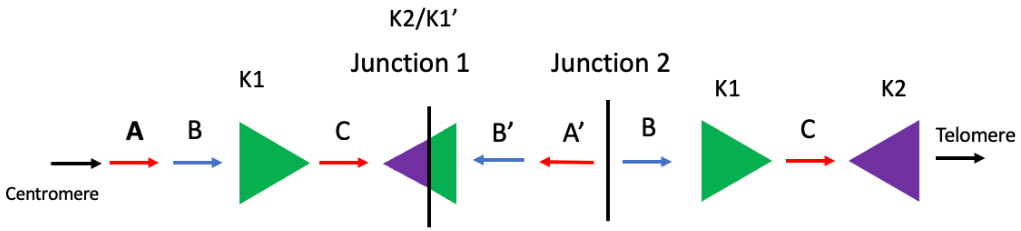


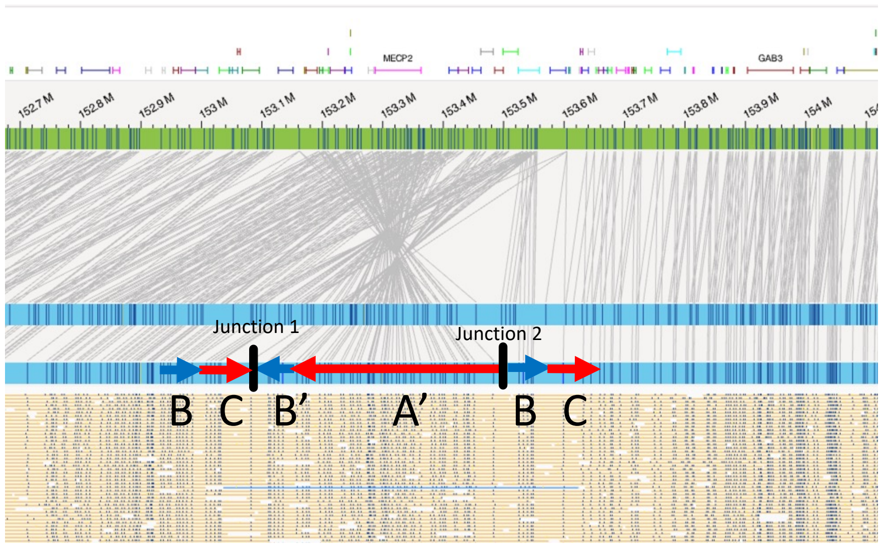

### BAB14604

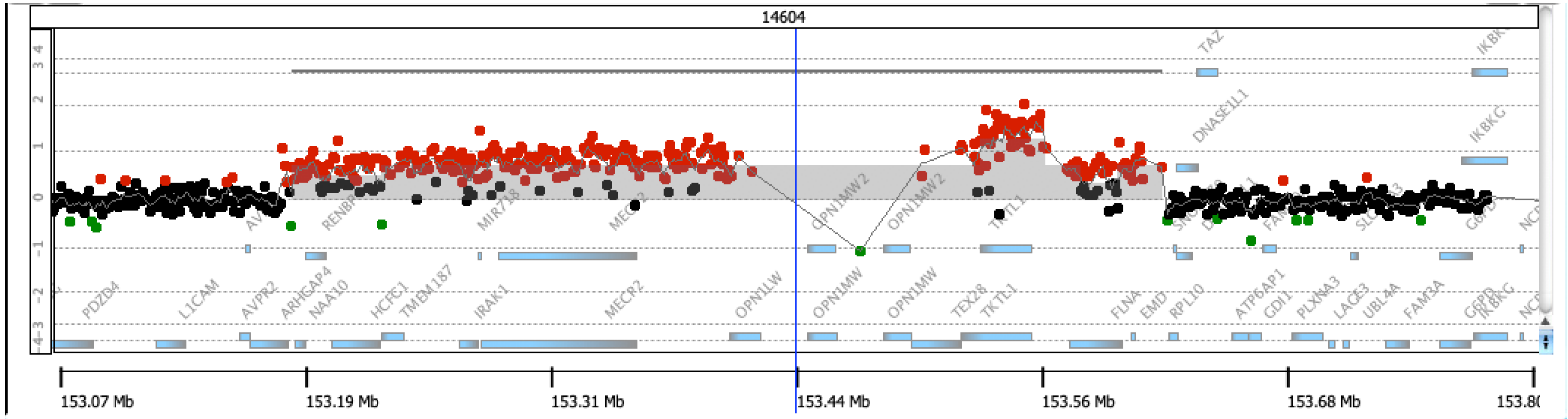


**Breakpoint Junction 2:**

chrX:153188686

chrX:152841838(+) CCAGGTGGCAGCTACTGTGCTAGTCCAGGCAAAAGACGGCAGGATTCAGACAAGCAGCAGCGAGGGAGGA

Junction 2 AGGGAGCGCCCTCTCAGAACCCCTCCTTAGACCTGACGGCAGGATTCAGACCAGCAGCAGCGAGGGAGGA

chrX:153078714(-) AGGGAGCGCCCTCTCAGAACCCCTCCTTAGACCTGGCTTCCCTCTCCTCACTTTTCCATTCCCATAGCGC

chrX:153499735

**Haplotype Structure 3:**

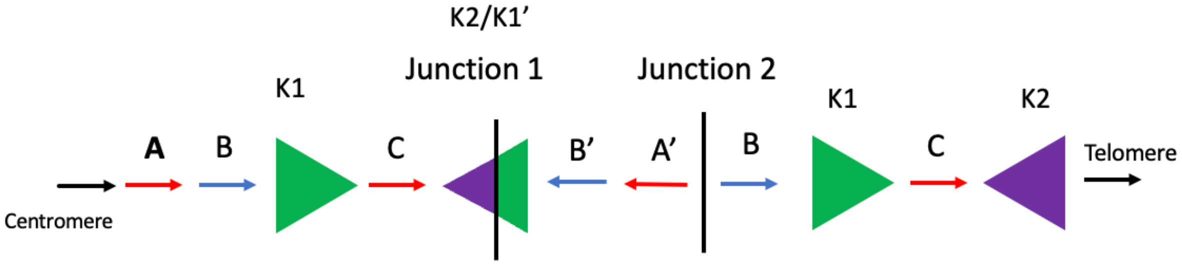


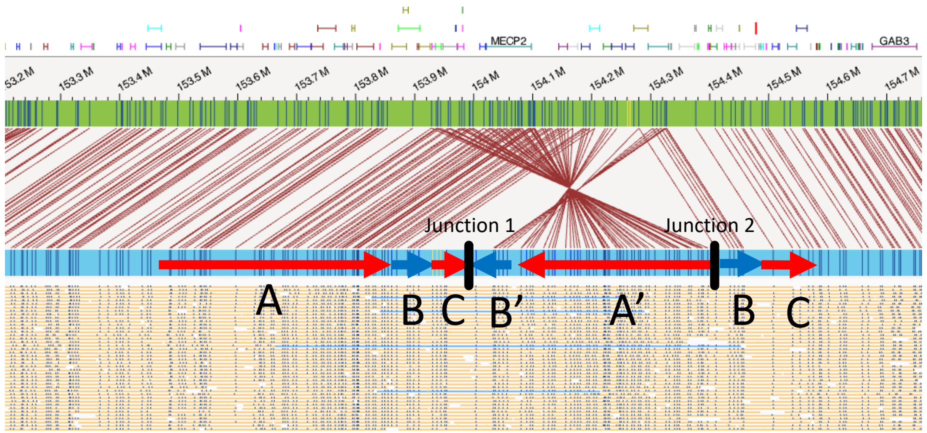

### BAB14686

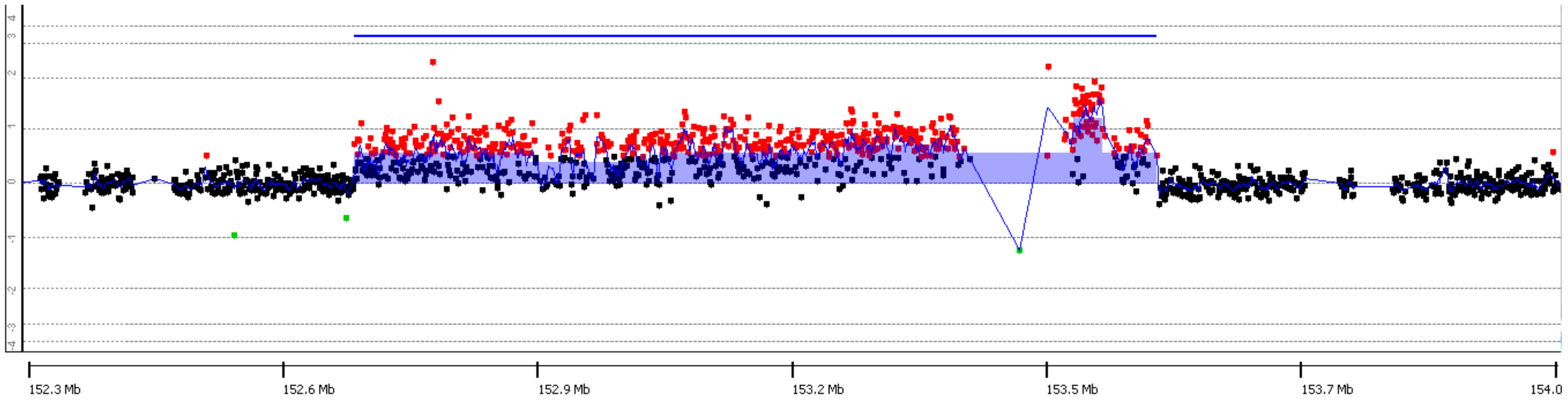


#### Breakpoint Junction 2:

chrX:152754451

chrX:152754420(+) CCTTCCCAACTGGGGAGGCAGGGGACTCGCACTTGGGAAATGTACCTGCGGGCTCTTGGGGTCGTCACCTGGC

Junction 2 GCAGAGCTGGAAGAAGCCAAGAGGTTCCACACTTGGGAAATGTACCTGCGGGCTCTTGGGGTCGTCACCTGGC

chrX:153441231(-) GCAGAGCTGGAAGAAGCCAAGAGGTTCCACATCAGCCTCCAGGAGTCCTATCACAGCCTAAAGGAGAGGTCTC

ChrX:153441203

#### Haplotype Structure 2:

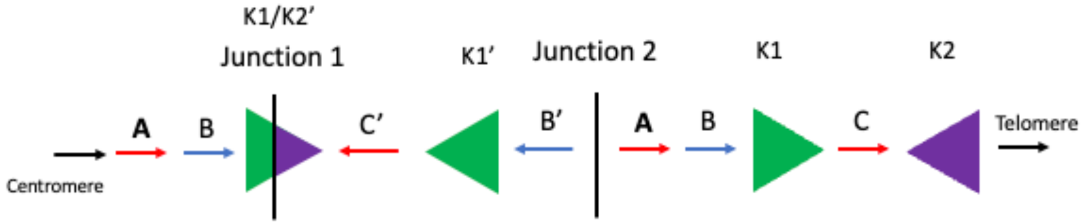


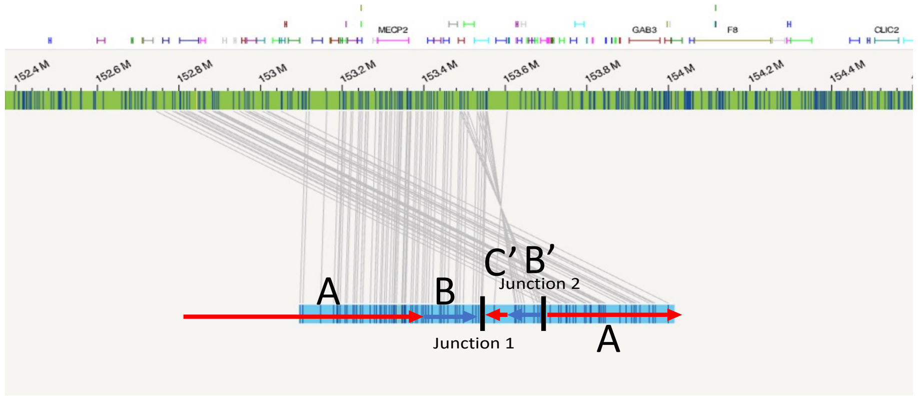

### BAB15418

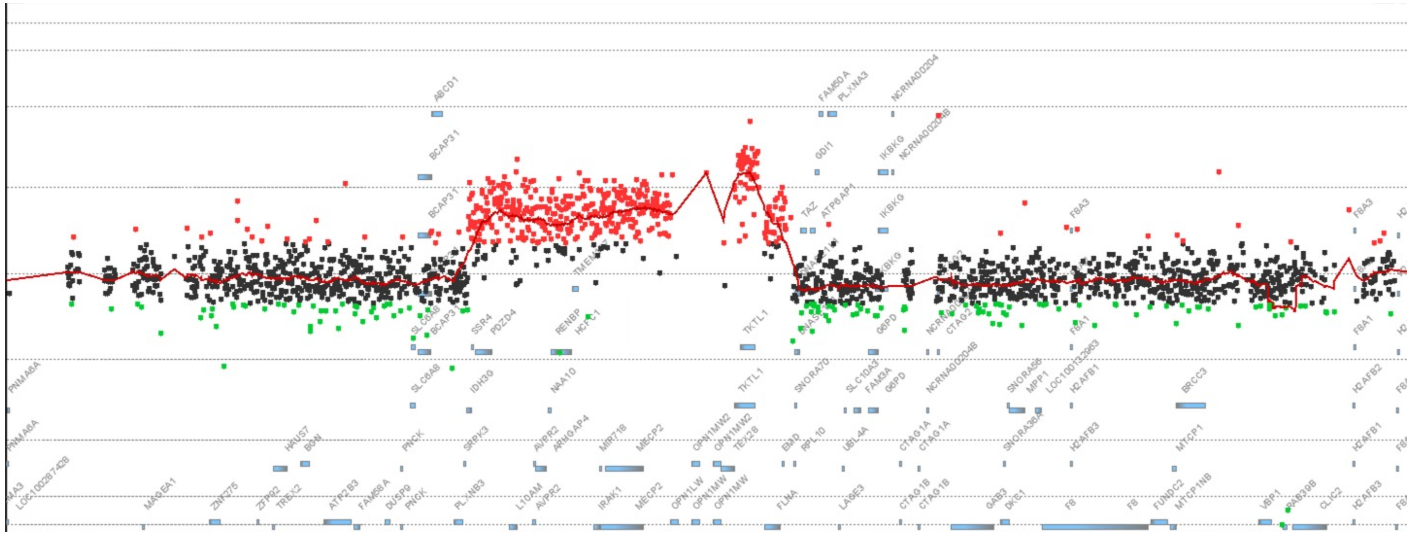

#### Breakpoint Junction 2:

```
                                                                  chrX:153054923
chrX:153054892(+) GGGAGGCTGAGGCAGGAGAATCGCTTGAAACTGGAAGGCAGAGGTTGCACTGAGCCGAGATCACACCGC
       Junction 2 CTGCCTCAGCCTCTCGAGTATCTGGGACTACTGGAAGGCAGAGGTTGCACTGAGCCGAGATCACACCGC
chrX:153413610(-) CTGCCTCAGCCTCTCGAGTATCTGGGACTACAAGCCGTGCATTACCACAATGGCAATTTTTTTTATTTC
                                        chrX:153413582
```

#### Haplotype Structure 4:

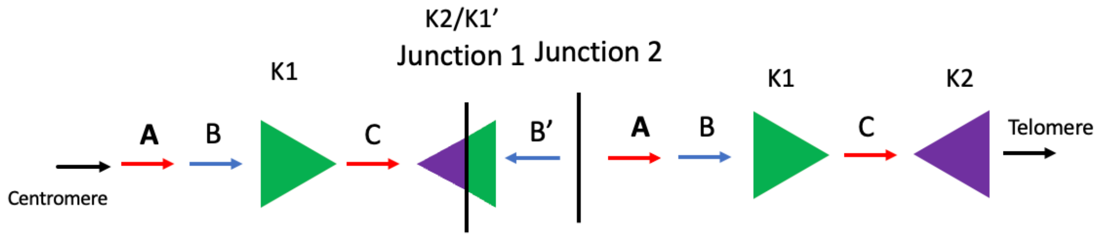


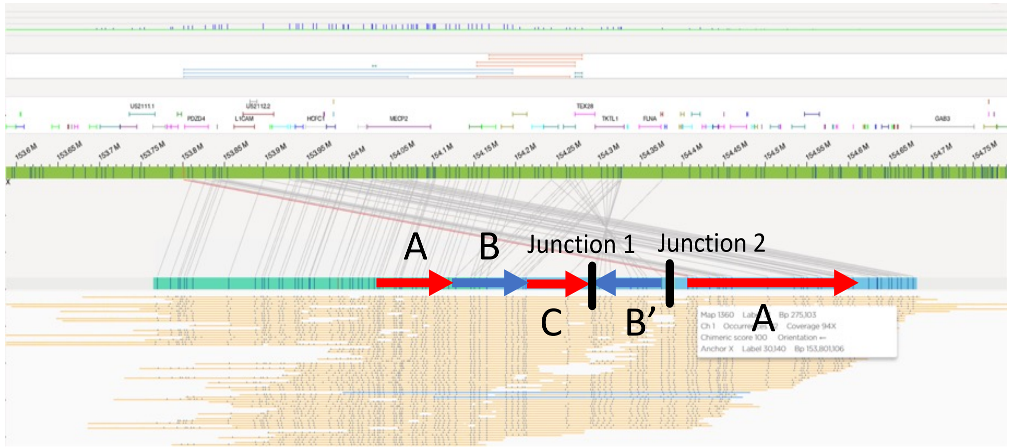

### BAB15428

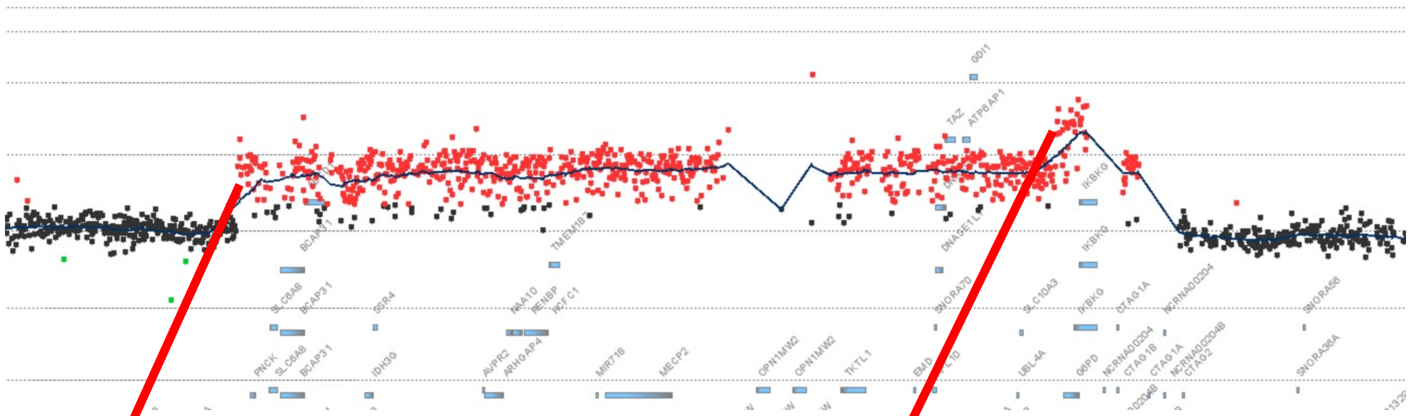

**Breakpoint Junction 2:**
Within *Alu* Repetitive Elements

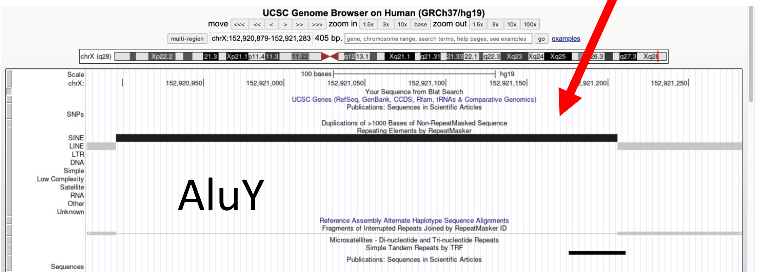


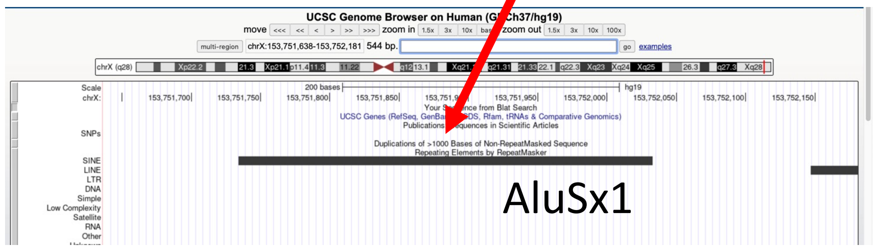


**Haplotype Structure 3**

*Or*

**Haplotype Structure 1**

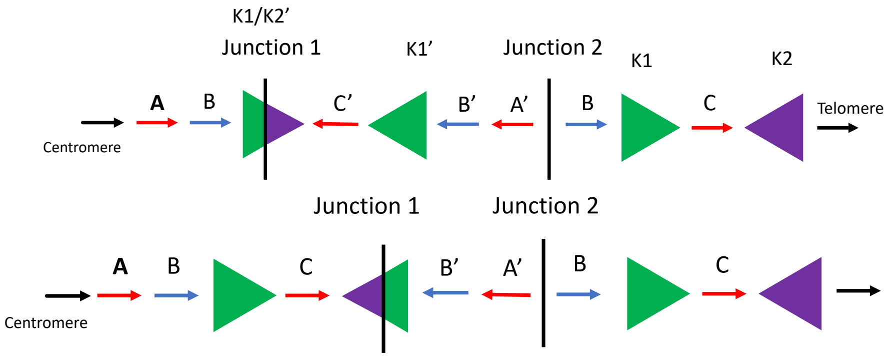


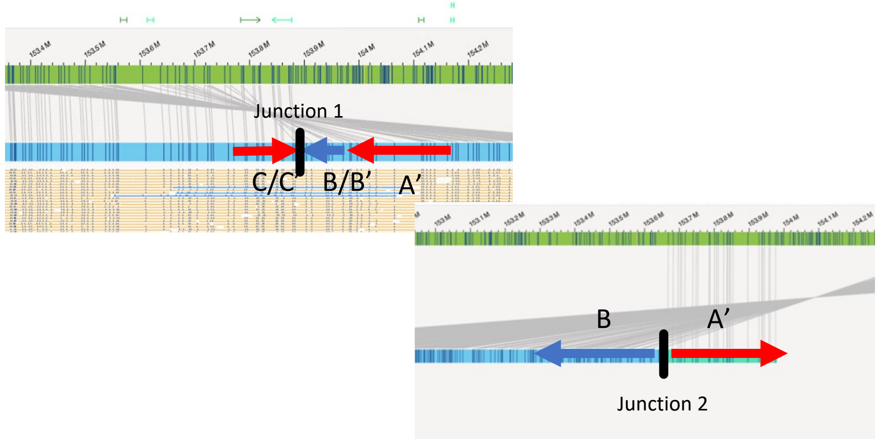

### BAB15702

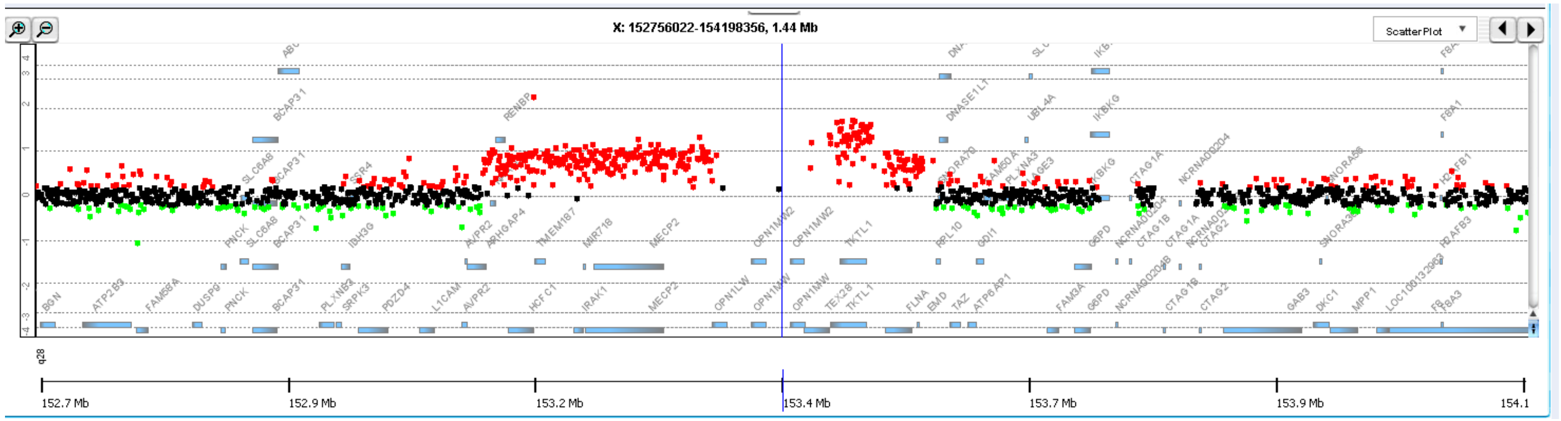


#### Breakpoint Junction 2:

```
                                                                 ChrX:153187761
chrX:153187715(+)  GATCACAGTGCTGCTCAGGGCAAGCCTGCAGCGATCTCGCAGTCTCAGTGGCTGGGGAAGCCTGCCCTCAGGCCCCAGGGGAAAGG
       Junction 2  CAATCTCACTCTTGTCAGTCAGGCTGGAGTGCAGTGGCGCAGTCTCAGTGGCTGGGGAAGCCTGCCCTCAGGCCCCAGGGGAAAGG
chrX:153446662(-)  CAATCTCACTCTTGTCAGTCAGGCTGGAGTGCAGTGGCGCAGTCTCGGCTCACTGCAACCTCCGTCTCCCCGGCTCAAGCAATTCT
                                                ChrX:153446626
```

#### Haplotype Structure 6:

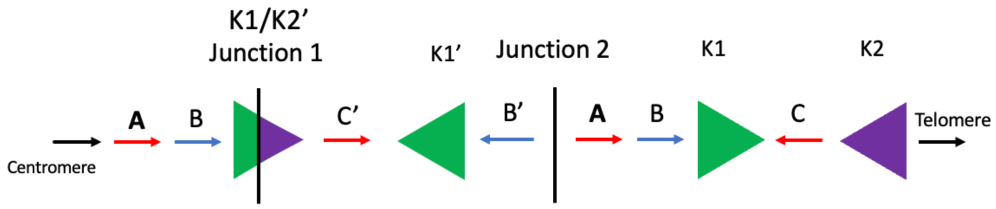


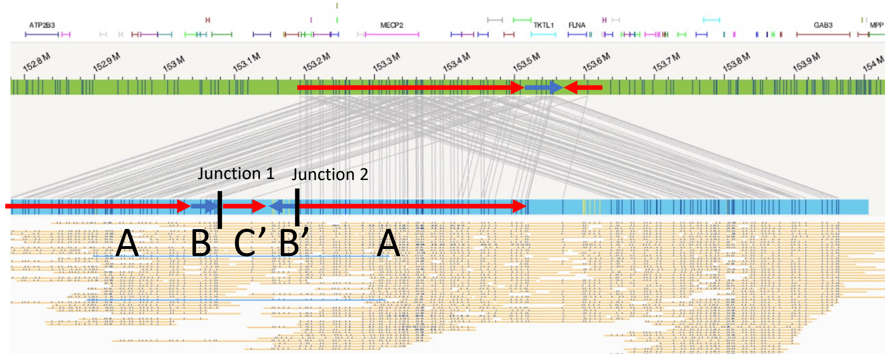

### BAB15705

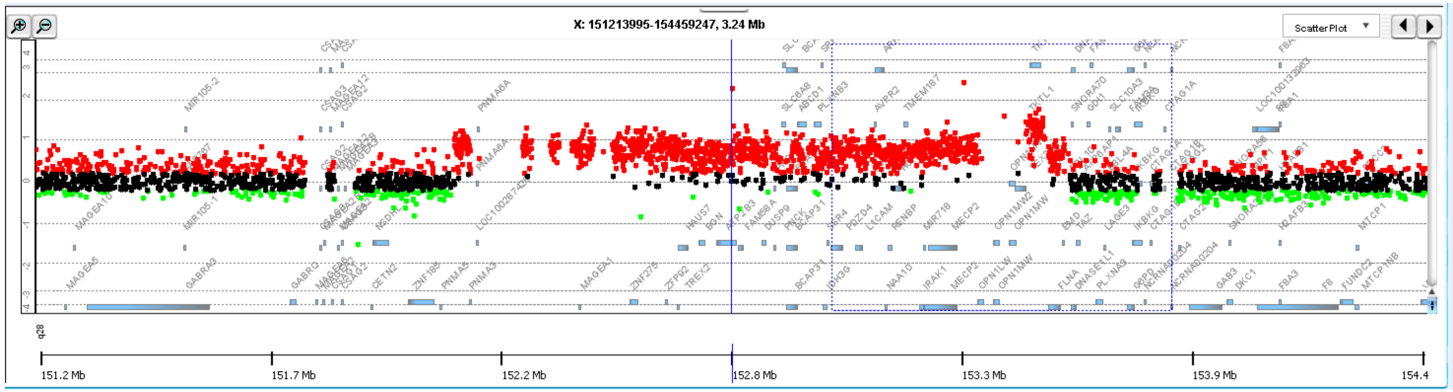


#### Breakpoint Junction 2:

ChrX:152190650

chrX:152190603(+) ACACAATCTTCGTCAAAATTTAAAGGTGTGGAGGCTGAGGCAGGAGAATGGCGTGAACCCGGGAGGCGGAGCTTGCA

Junction 2 GTGGCACATGCCTGTAGTCCCTGCTACATGGGAGGCTGAGACAGGAGAATGGCGTGAACCCGGGAGGCGGAGCTTGCA

chrX:153431409(-) GTGGCACATGCCTGTAGTCCCTGCTACATGGGAGGCTGAGACAGGAGGATCGCTTGAGCCCGAGAGTTTTATGTTGCA

ChrX:153468511

#### Haplotype Structure 2:

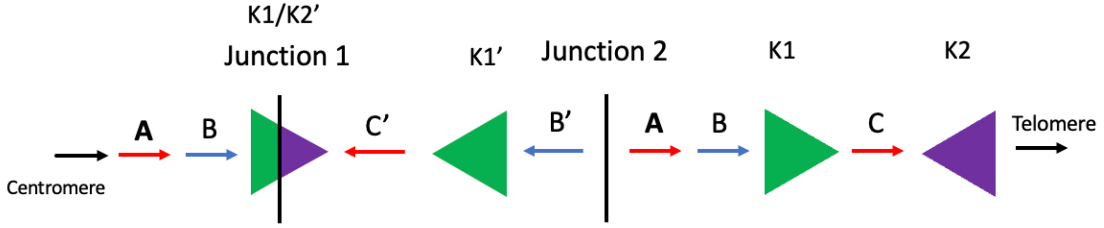


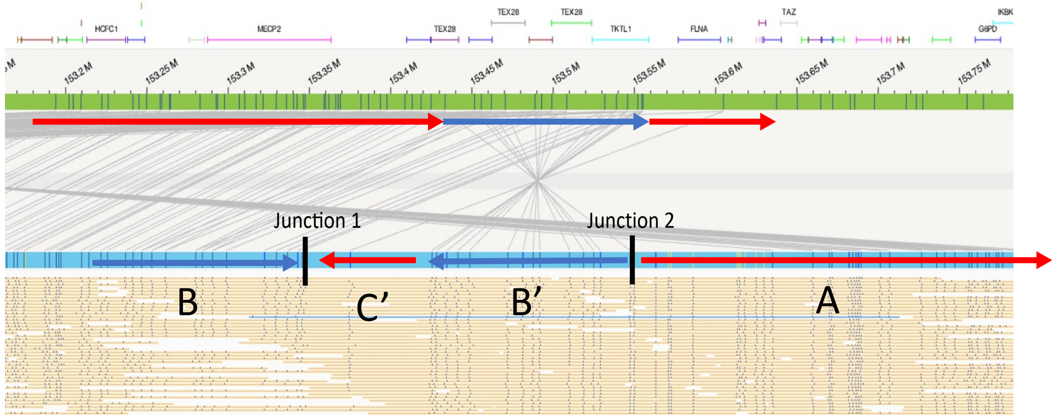

### BAB15740

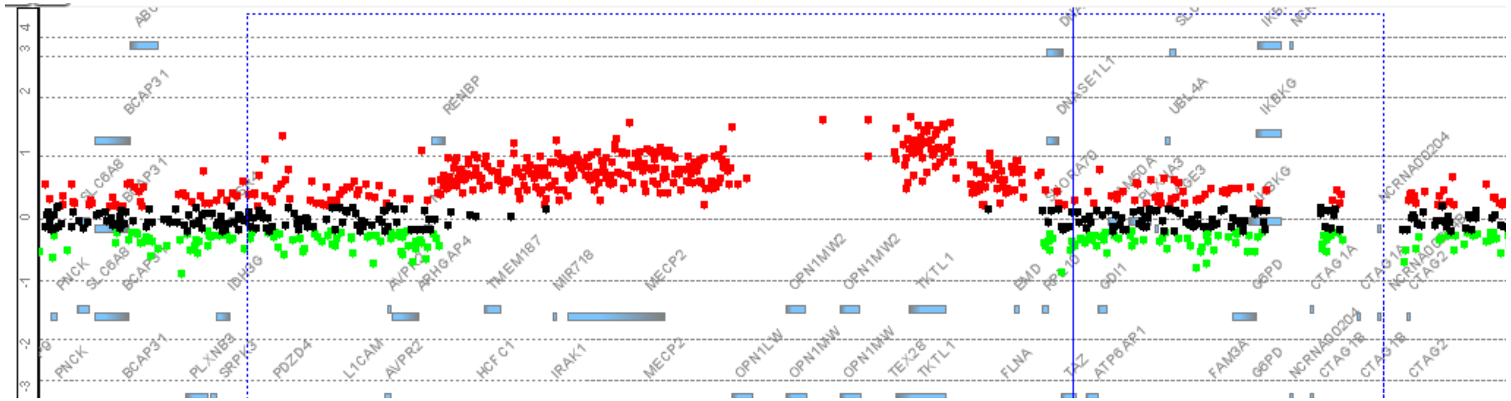

#### Breakpoint Junction 2:

```
                                                                                  ChrX:153205884
ChrX:153205843(+) CCTGTAGATTCTTGAATAGACACCAGGGCCCCAGACGAATGGCATATATGTCTGAGCATCAGCGTTTCGCCAGCTCCCTGG
       Junction 2 TTGTTTACTGACATCTGAAGAGACTCTTCAAGAATGCACTGGCATATATGTCTGAGCATCAGCGTTTCGCCAGCTCCCTGG
ChrX:153428346(-) TTGTTTACTGACATCTGAAGAGACTCTTCAAGAATGCACTGAAGAATATCTTCATGTATCTTTACCAATACATTGATGAAG
                                                              ChrX:153428308
```

#### Haplotype Structure 2:

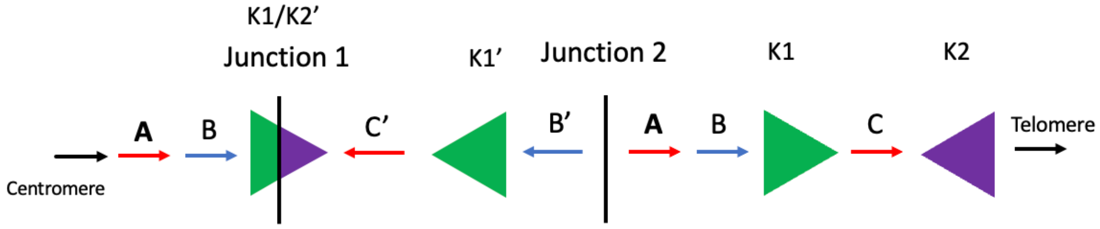


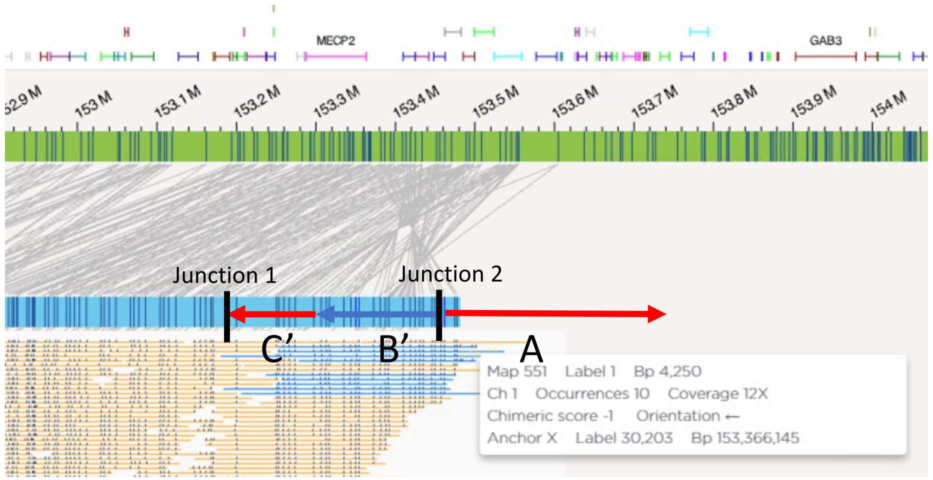

### BAB15789

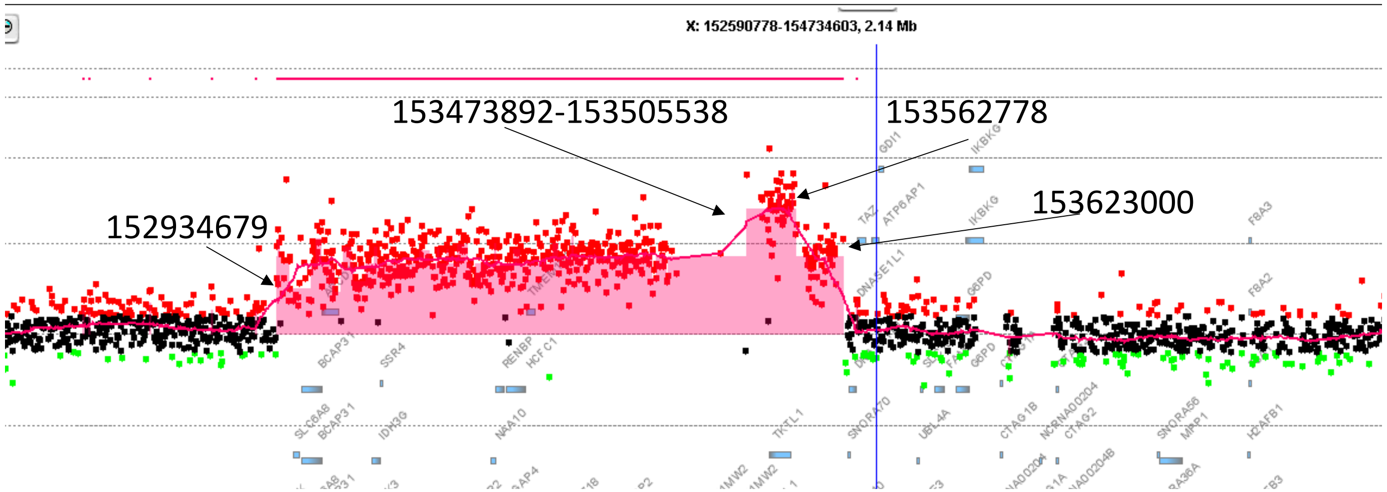


**Breakpoint Junction 2:**

ChrX:152934497(+)

| | |
|---|---|
| ChrX:152934457(+) | GTTGGTGGGGCTGGGGCCACTCAGACCAGAGGACAGGC**CA**CGGCTCTCCACTTCAGAAGCCACCGCCAAGCTGCCACTG |
| Junction 2 | TTCCAGCCCTGAGTGGAGCACAGCC**T**GCCCCAGGCAGA**CA**CGGCTCTCCACTTCAGAAGCCACCGCCAAGCTGCCACTG |
| ChrX:153421664(-) | TTCCAGCCCTGAGTGGAGCACAGCCGGCCCCAGGCAGA**CA**TGGCTGCCTCAGGCTGGCCCTGGGGAGTTCTAGAATTG |

ChrX:153421627(-)

**Haplotype Structure 2:**

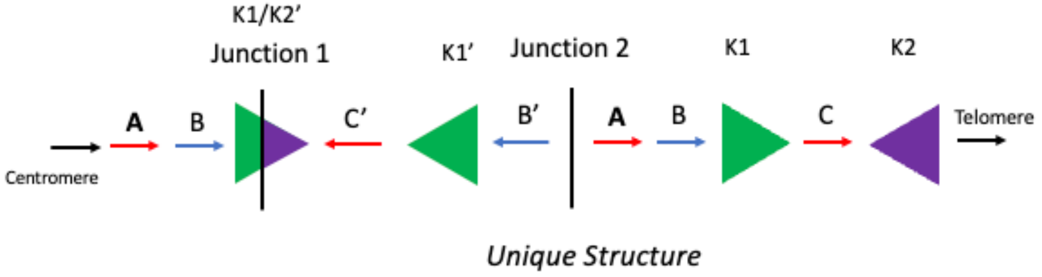


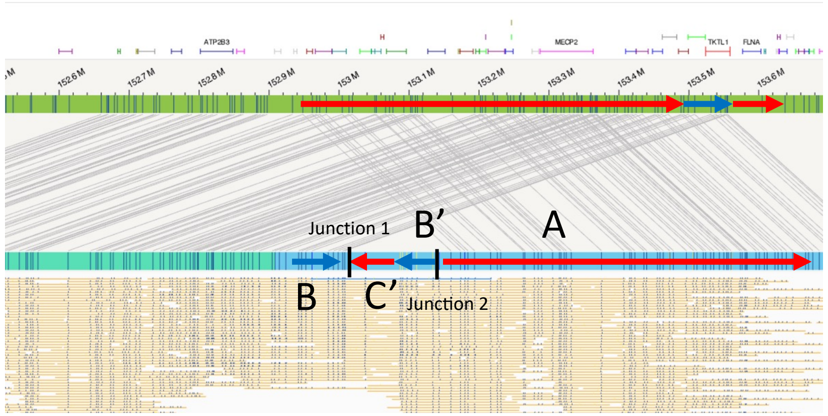

### BAB15421

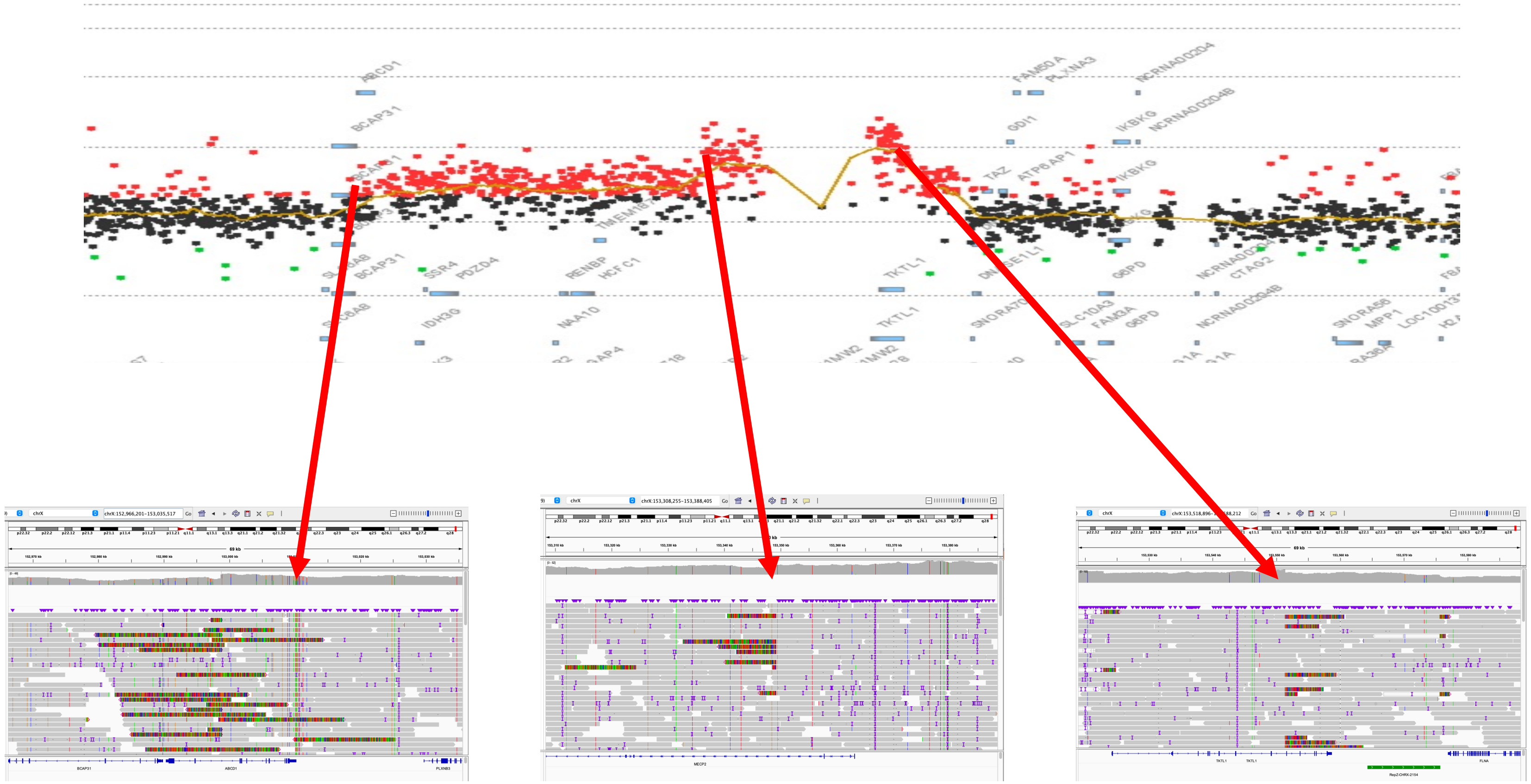

**Breakpoint Junction 2:**
***Complex Unresolved***

**Haplotype Structure:** ***Complex Unresolved***

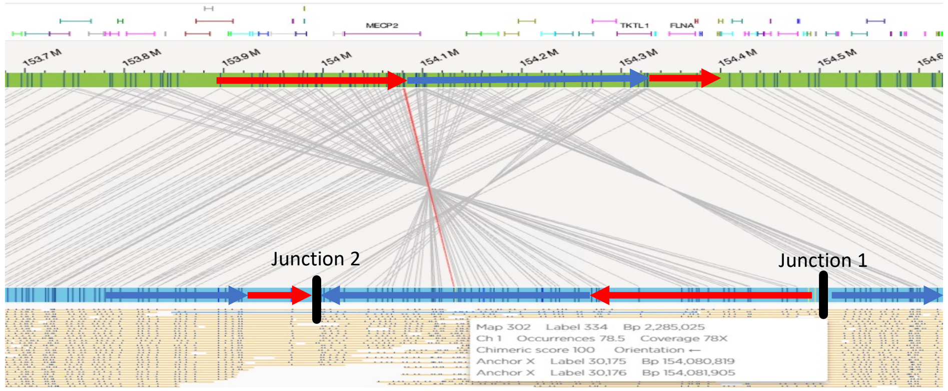
