## Additional File 3 for "Break-induced replication underlies formation of inverted triplications and generates unexpected diversity in haplotype structures"

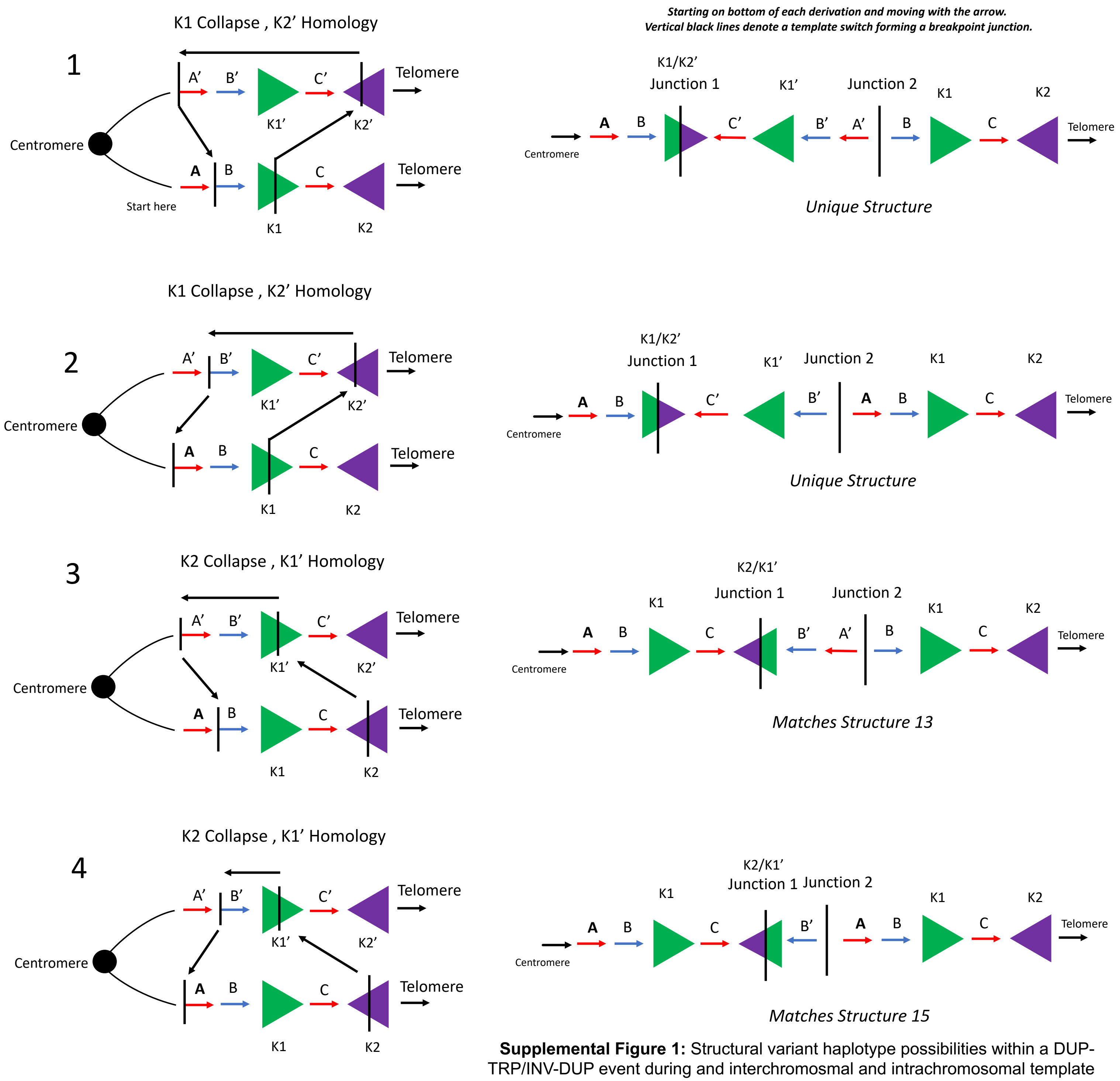


**Supplemental Figure 1:** Structural variant haplotype possibilities within a DUP-TRP/INV-DUP event during and interchromosmal and intrachromosomal template switch between inverted LCRs.

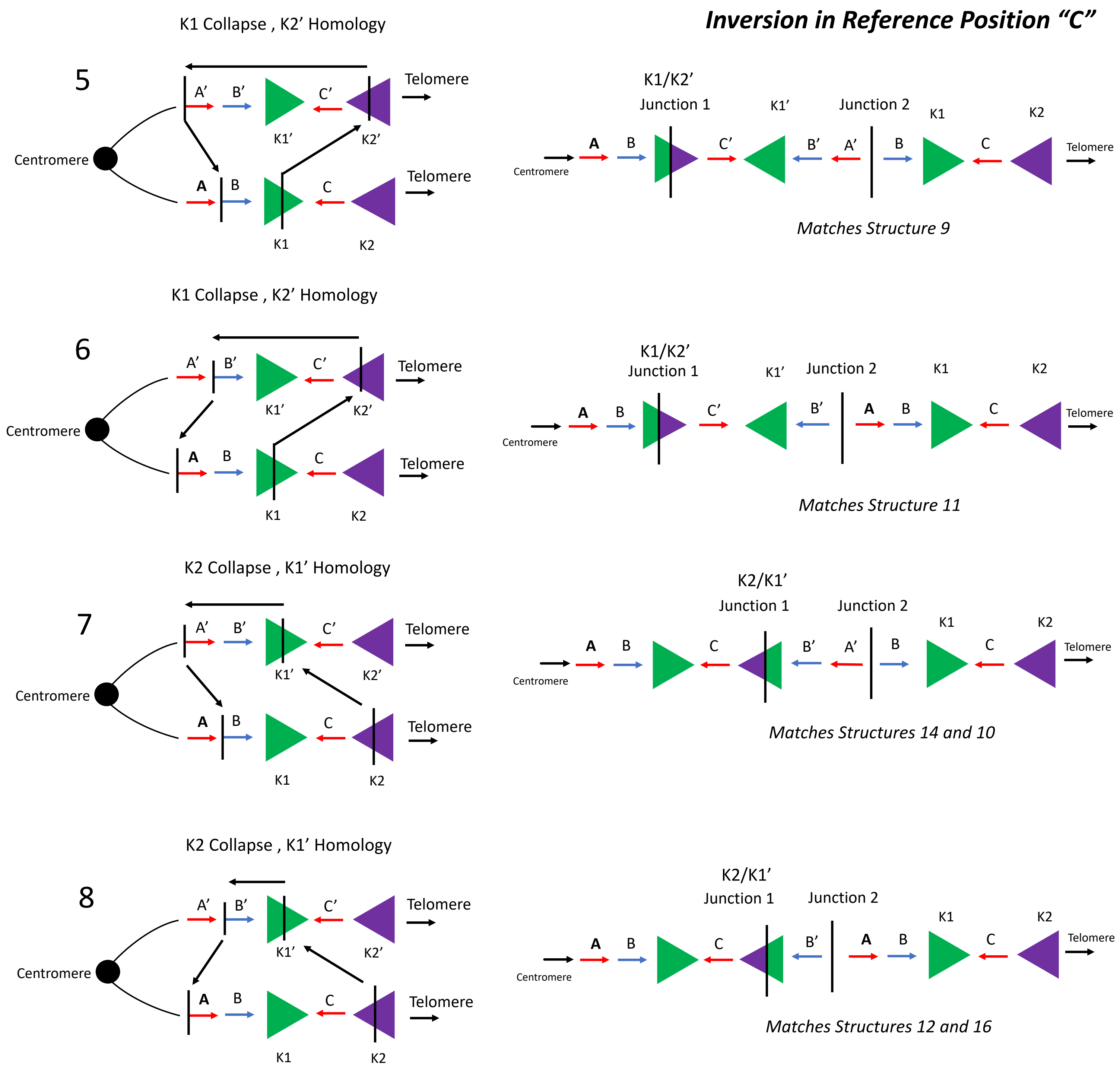

Inversion in Reference Position "C"
K1 Collapse , K2' Homology
5
A' B' C'
Telomere
K1' K2'
Centromere
A B C
Telomere
K1 K2
K1/K2'
Junction 1
K1'
Junction 2
K1
K2
Centromere
A B C' B' A' B C
Telomere
Matches Structure 9
K1 Collapse , K2' Homology
6
A' B' C'
Telomere
K1' K2'
Centromere
A B C
Telomere
K1 K2
K1/K2'
Junction 1
K1'
Junction 2
K1
K2
Centromere
A B C' B' A B C
Telomere
Matches Structure 11
K2 Collapse , K1' Homology
7
A' B' C'
Telomere
K1' K2'
Centromere
A B C
Telomere
K1 K2
K2/K1'
Junction 1
Junction 2
K1
K2
Centromere
A B C B' A' B C
Telomere
Matches Structures 14 and 10
K2 Collapse , K1' Homology
8
A' B' C'
Telomere
K1' K2'
Centromere
A B C
Telomere
K1 K2
K2/K1'
Junction 1
Junction 2
K1
K2
Centromere
A B C B' A B C
Telomere
Matches Structures 12 and 16

#### *Interchromosomal Event Mixture of Inversion "C" 1/2*

9

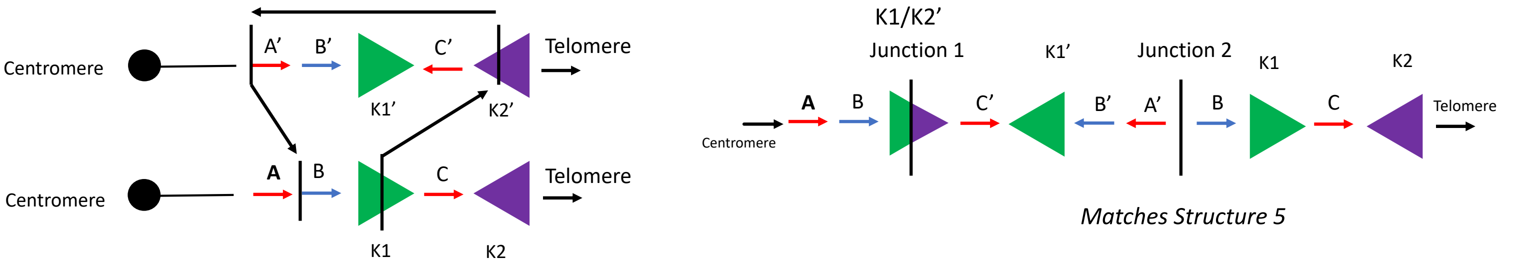


*Matches Structure 5*

10

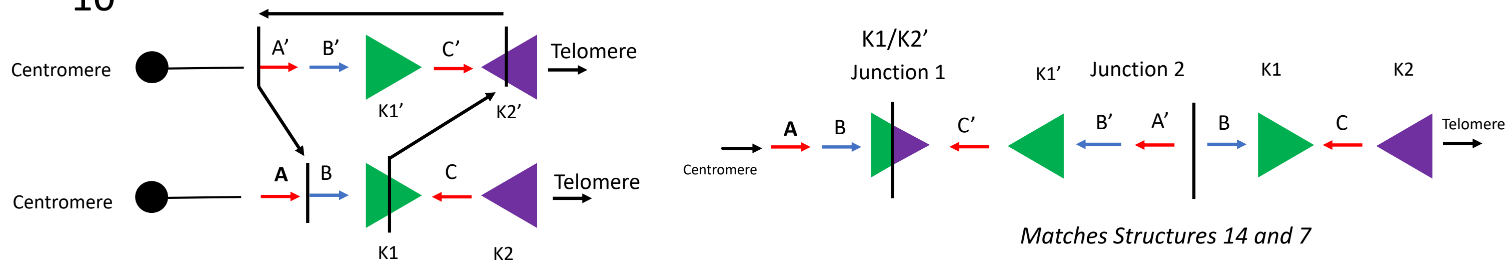


*Matches Structures 14 and 7*

11

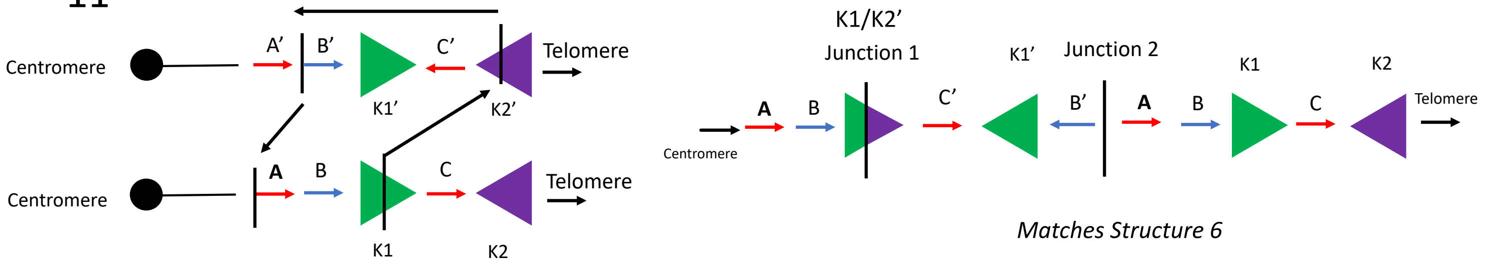


*Matches Structure 6*

12

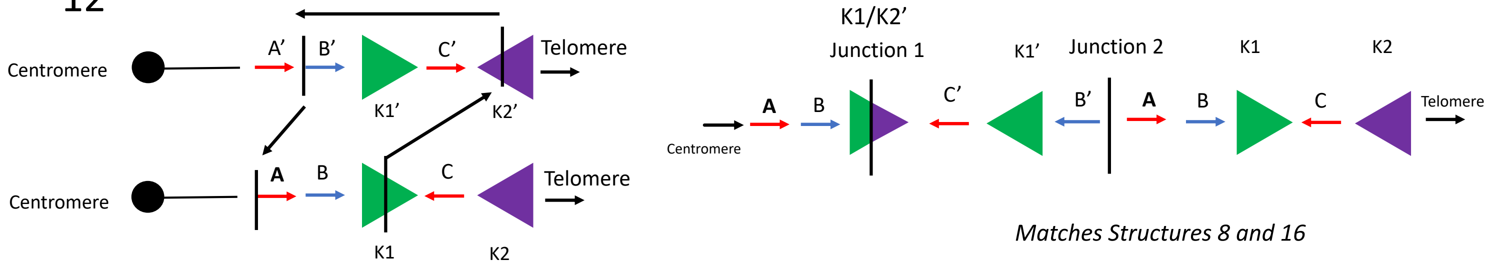


*Matches Structures 8 and 16*

### *Interchromosomal Event Mixture of Inversion "C" 2/2*

13

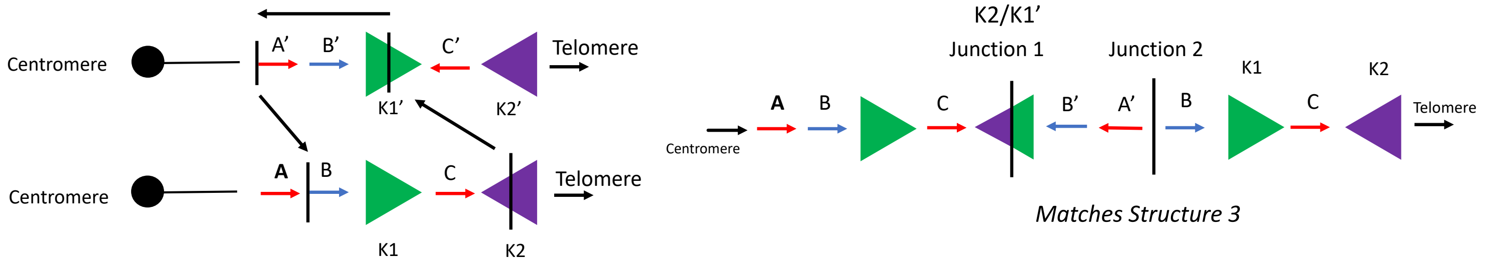


*Matches Structure 3*

14

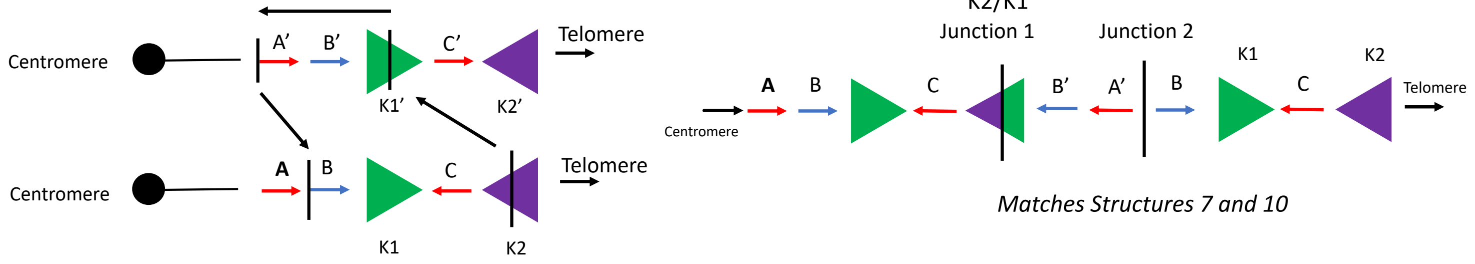


*Matches Structures 7 and 10*

15

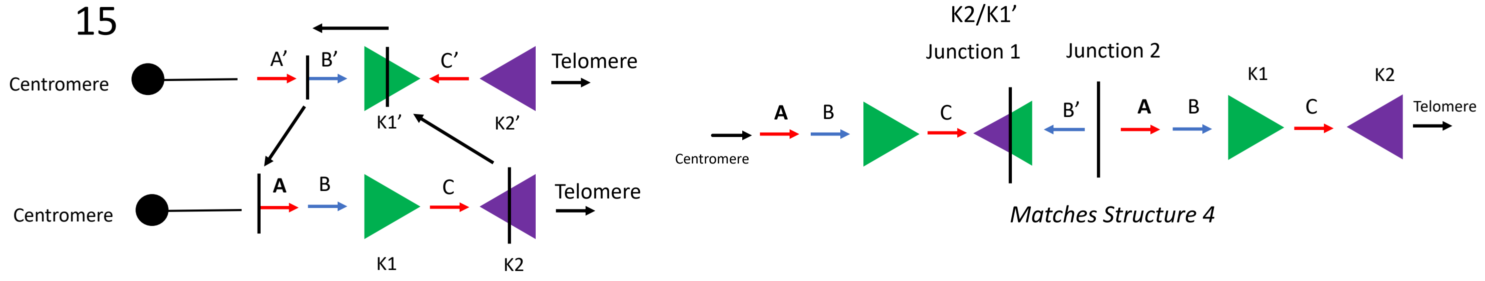


*Matches Structure 4*

16

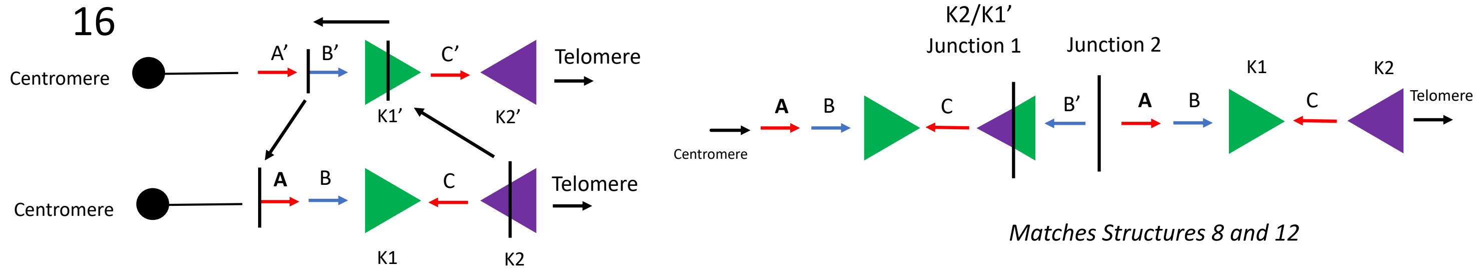


*Matches Structures 8 and 12*

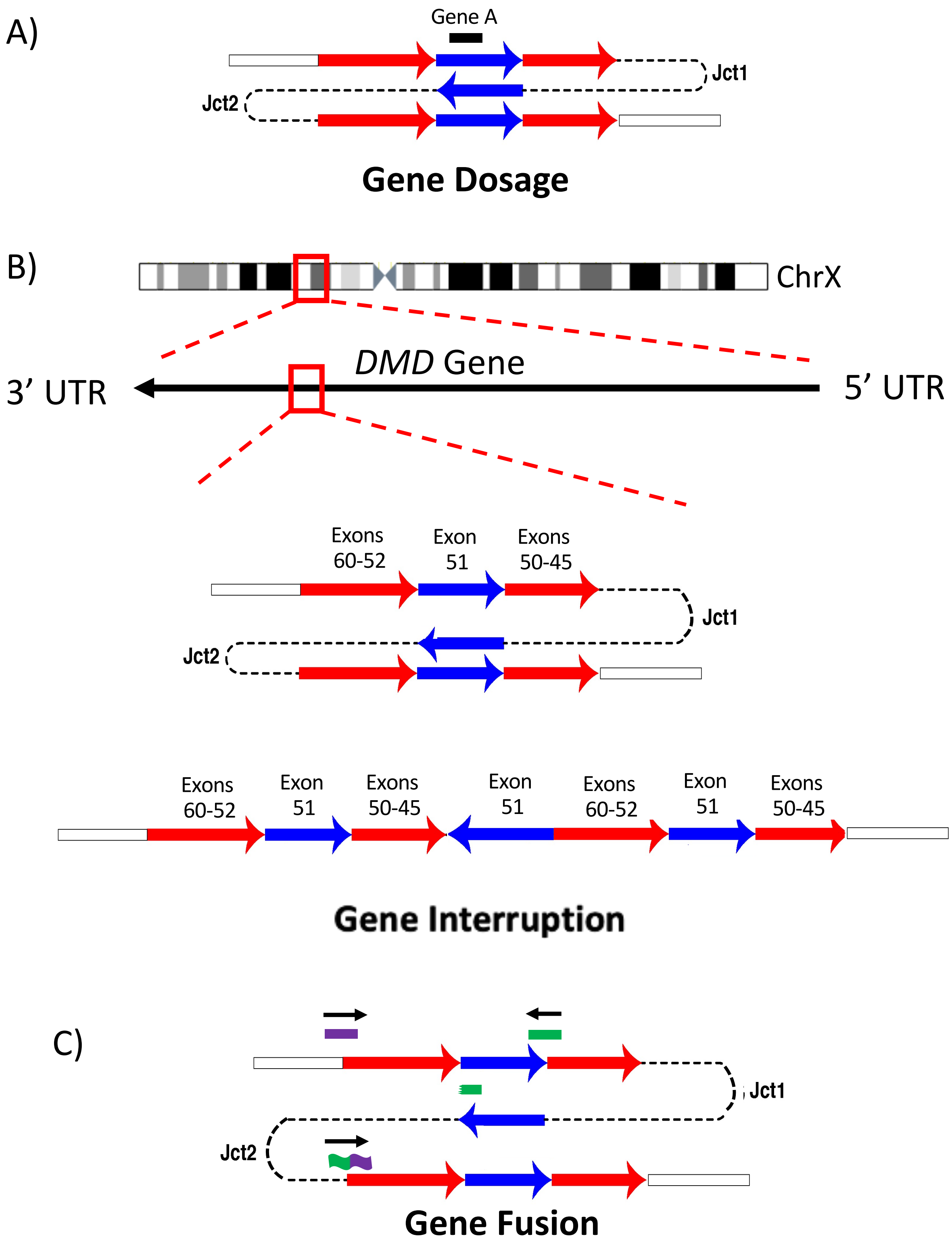


**Supplemental Figure 2:** Possible pathogenic effects of a DUP-TRP/INV-DUP event include gene dosage (A) gene interruption (B) as characterized by a disruption within the *DMD* gene [22] or gene fusion (C) events.

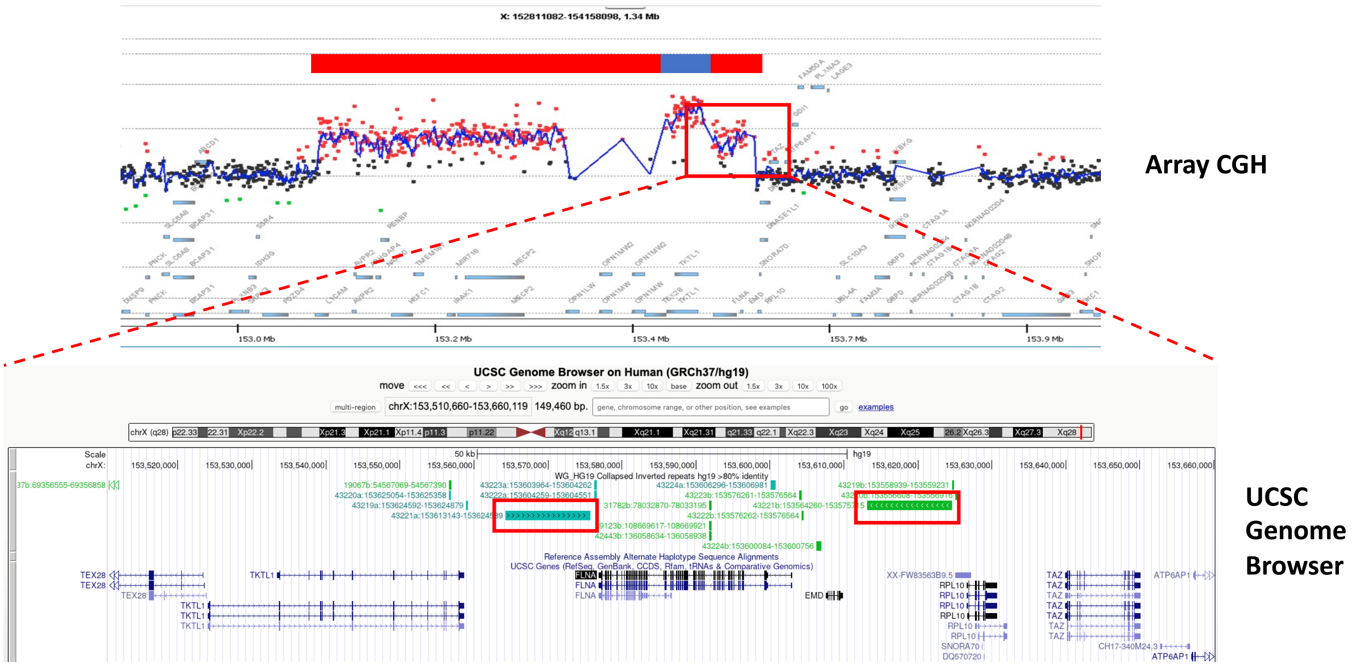


**Supplemental Figure 3:** Three pairs of inverted LCRs at Xq28 (43202a/43202a; 43221a/43221b (K1/K2); 43231a/43231b (L1/L2)) within the *MECP2* locus with the fourth pair being located at Xq22.1 (37696a/ 37696b). The corresponding location in the UCSC genome browser is shown.

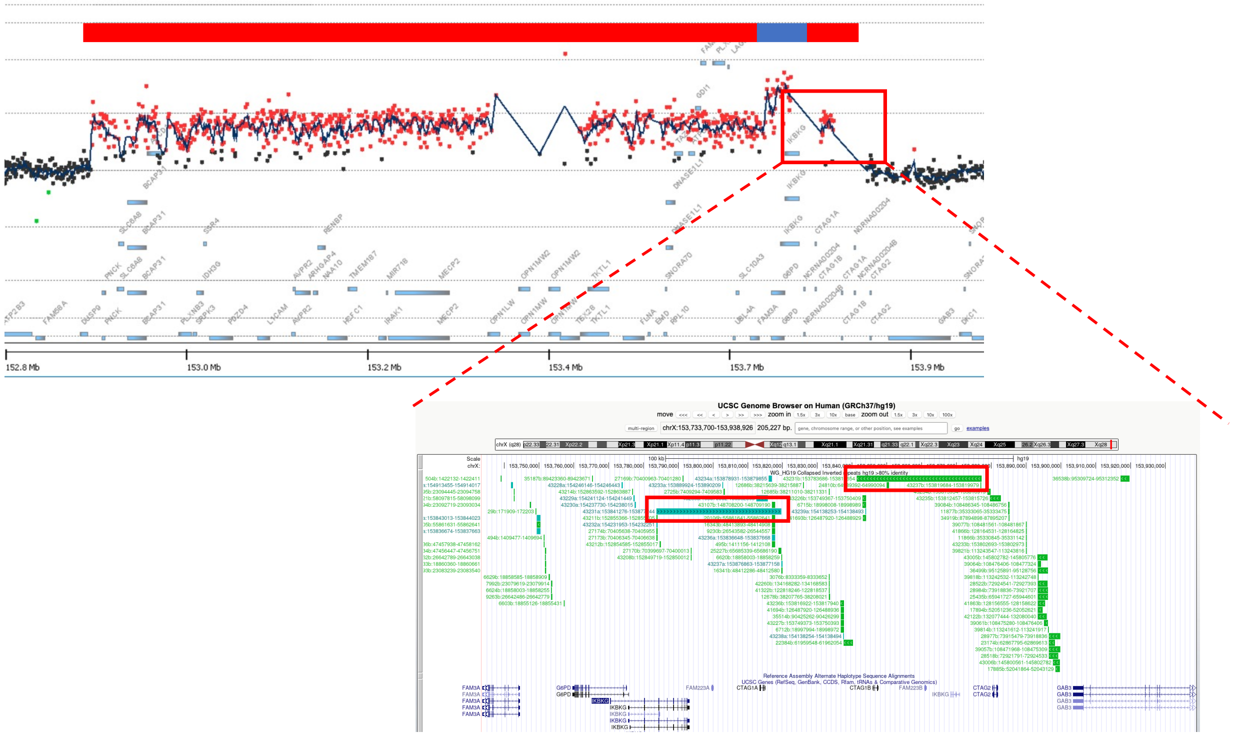

152.8 Mb
153.0 Mb
153.2 Mb
153.4 Mb
153.7 Mb
153.9 Mb
UCSC Genome Browser on Human (GRCh37/hg19)

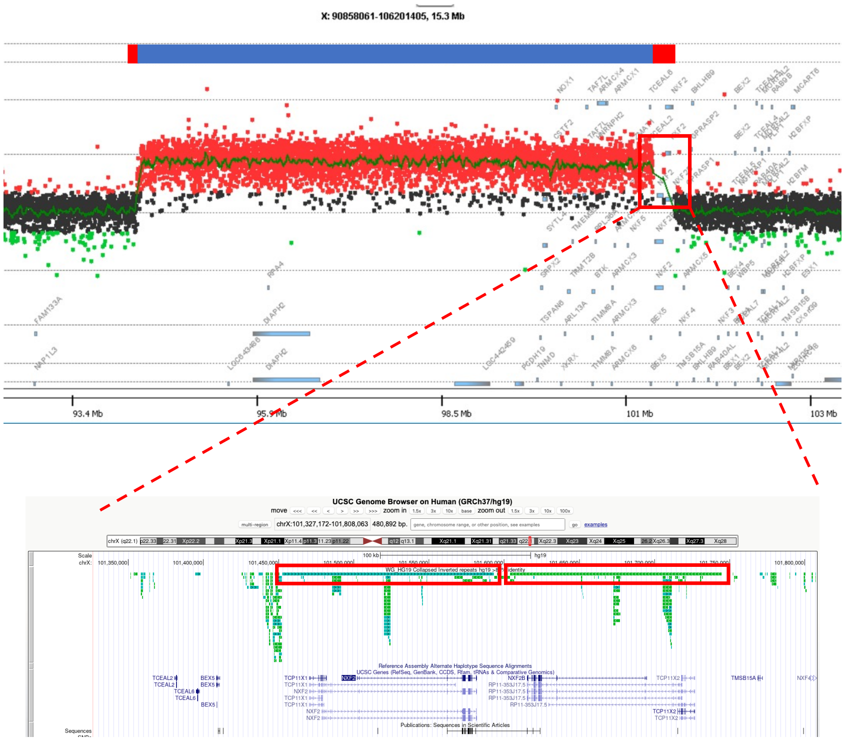
X: 90858061-106201405, 15.3 Mb
93.4 Mb
95.9 Mb
98.5 Mb
101 Mb
103 Mb
UCSC Genome Browser on Human (GRCh37/hg19)
chrX:101,327,172-101,808,063
480,892 bp.
WG_HG19 Collapsed Inverted repeats hg19
Reference Assembly Alternate Haplotype Sequence Alignments
UCSC Genes (RefSeq, GenBank, CCDS, Rfam, tRNAs & Comparative Genomics)
Publications: Sequences in Scientific Articles

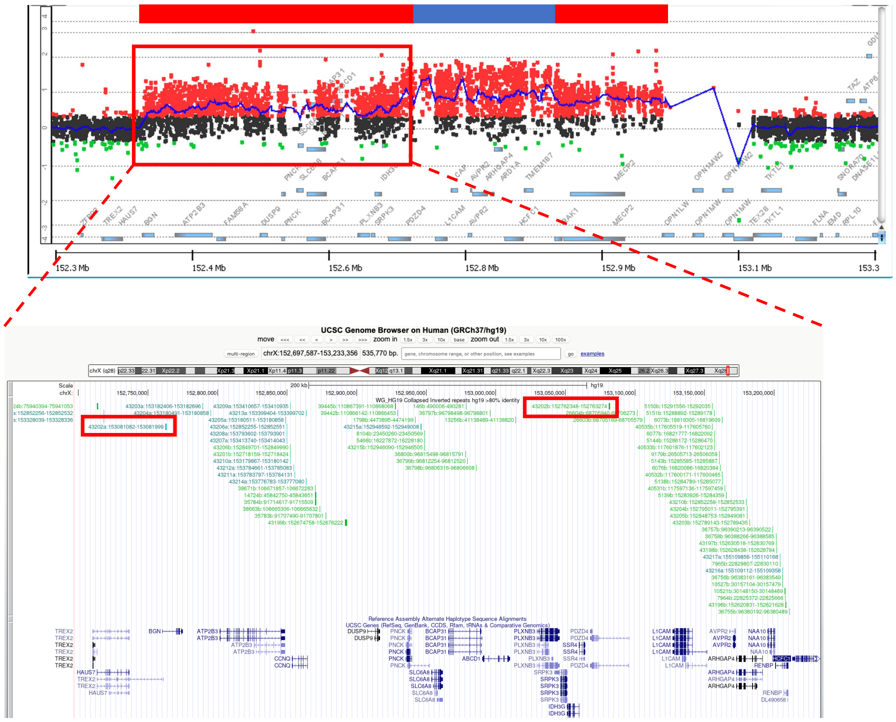
152.3 Mb
152.4 Mb
152.6 Mb
152.8 Mb
152.9 Mb
153.1 Mb
153.3
UCSC Genome Browser on Human (GRCh37/hg19)
chrX:152,697,587-153,233,356 535,770 bp.
WG_HG19 Collapsed Inverted repeats hg19 >80% identity
Reference Assembly Alternate Haplotype Sequence Alignments
UCSC Genes (RefSeq, GenBank, CCDS, Rfam, tRNAs & Comparative Genomics)
