## Additional File 4 for "Break-induced replication underlies formation of inverted triplications and generates unexpected diversity in haplotype structures"

**Additional File 4:** CRISPR-Cas9 targeted ONT data for BAB3114 family.

### COMPARING THE DUPLICATED FLNA INVERTED REPEATS

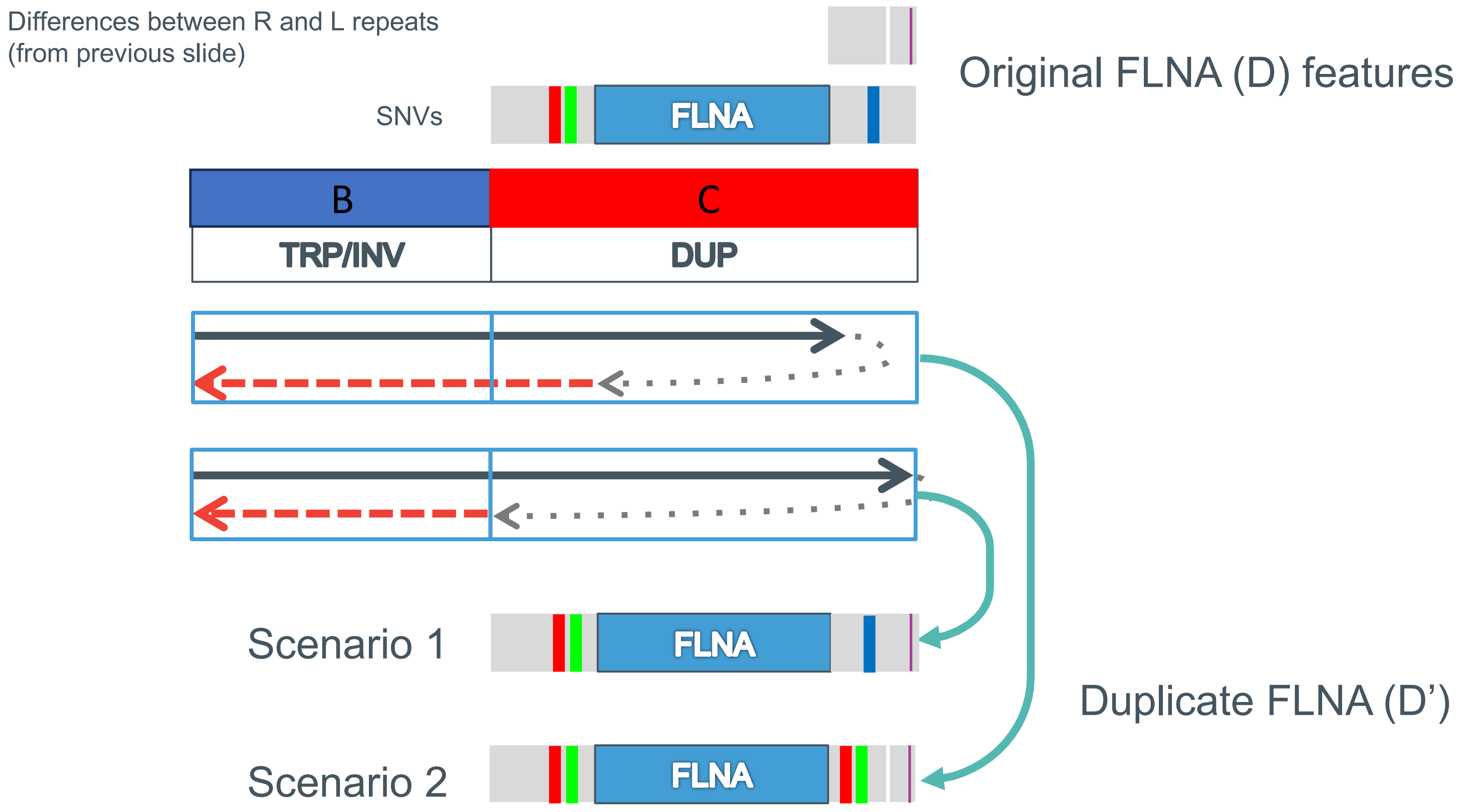

### GENERATE PHASED VARIANTS AND HAPLOTAG

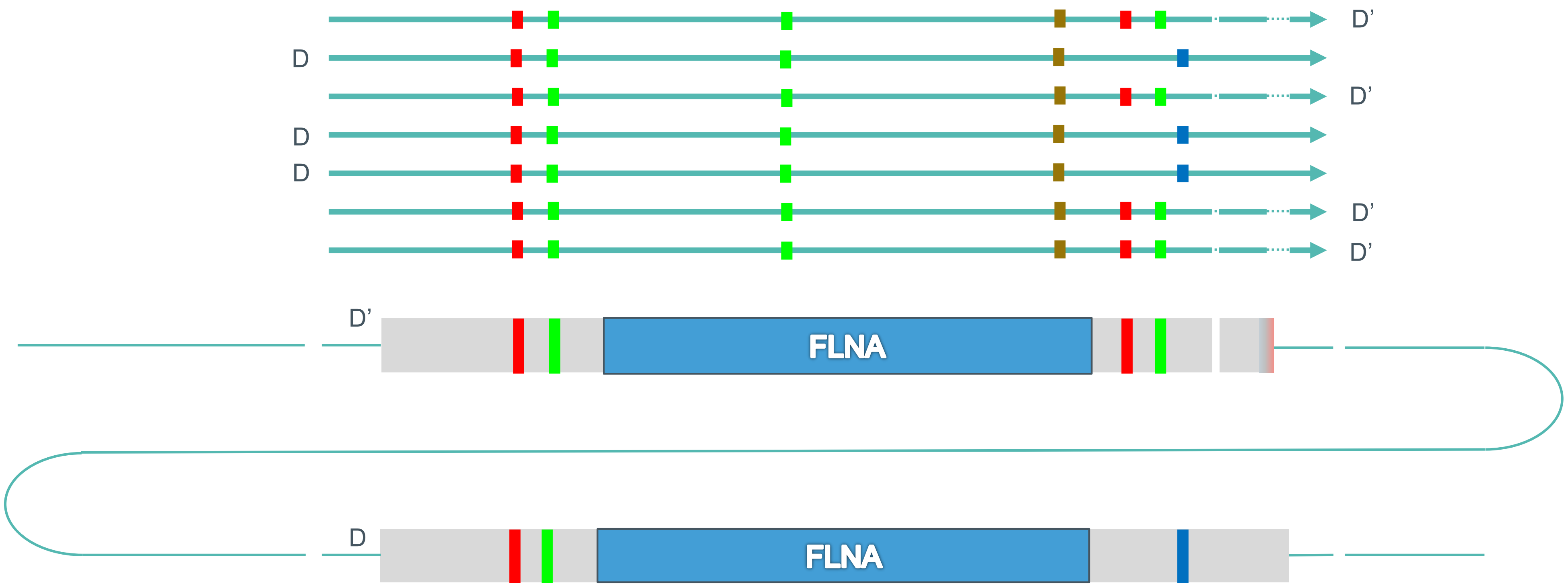

### BAB3114 HAPLOTYPING SEGREGATES READS INTO D AND D’

consensus

D reads

D’ reads

unassigned

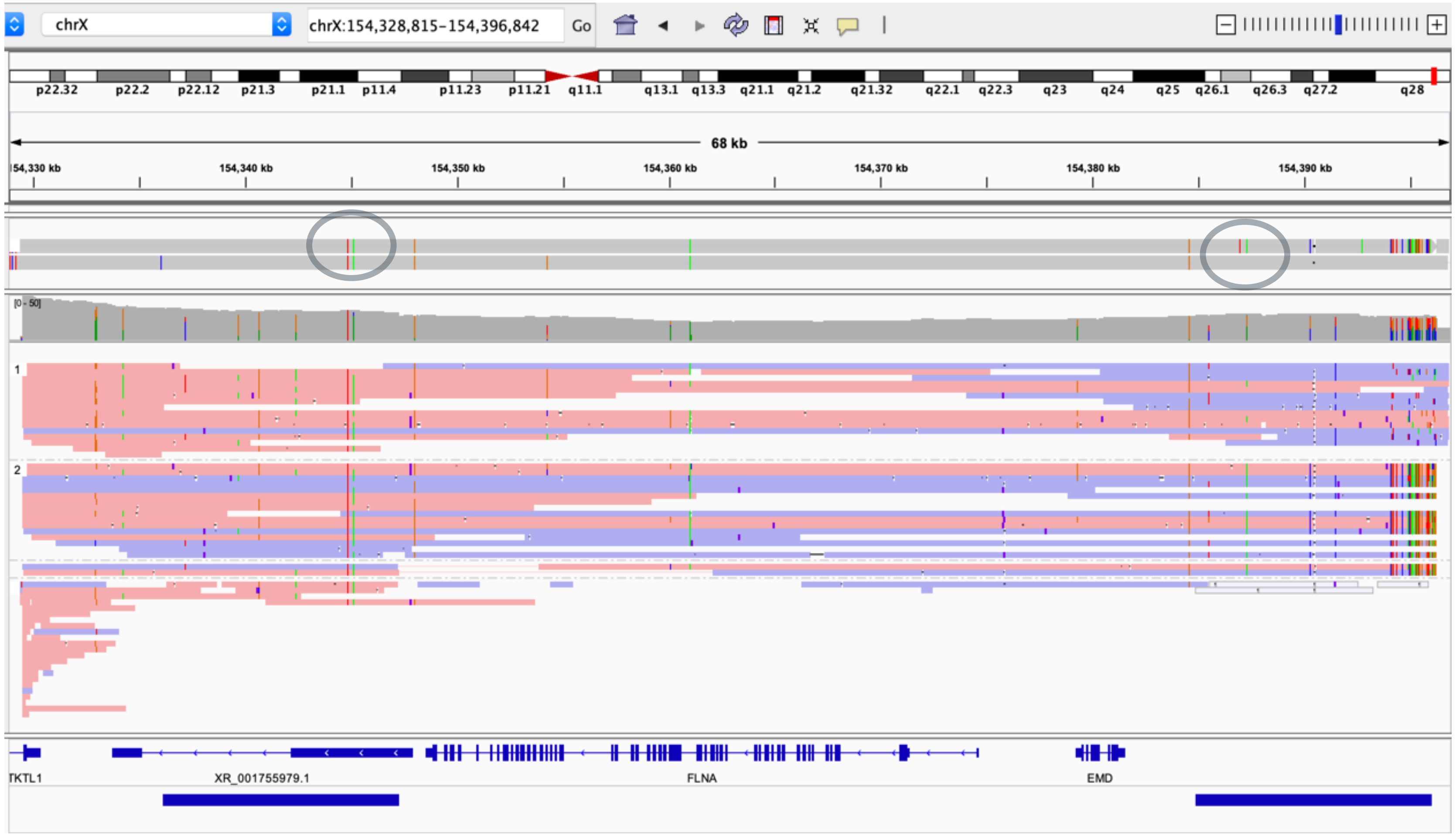

### BREAKPOINT RESOLUTION CONCLUSION

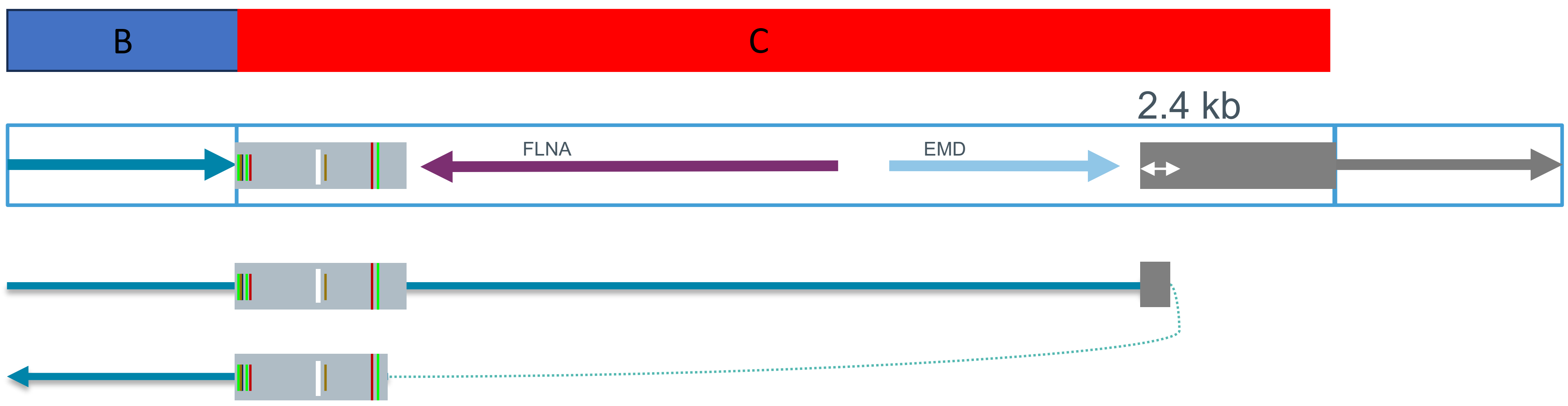


Resulting structure of the D’ element:

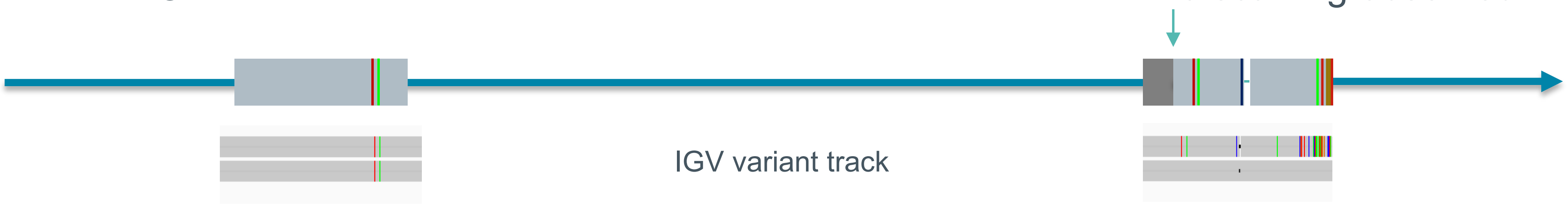

### A single 530 kb read resolves opsin cluster allele in BAB3114

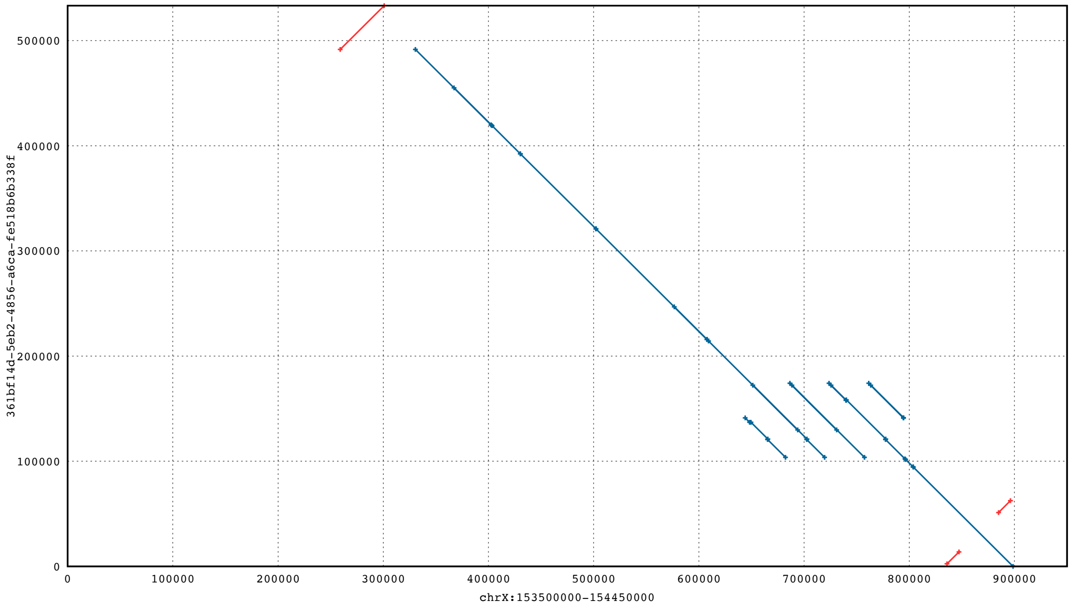


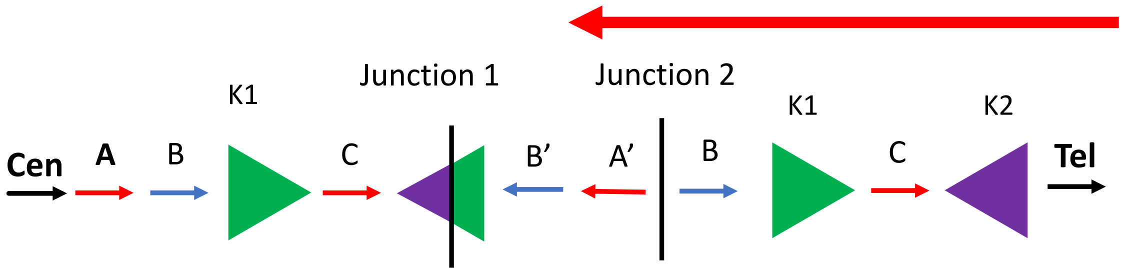


*Single 530 kb Nanopore read spanning the region*
