## Additional File 5 for "Break-induced replication underlies formation of inverted triplications and generates unexpected diversity in haplotype structures"

**Additional File 5:** StrandSeq data for samples BAB3114 and BAB14547.

### StrandSeq for BAB14547 and BAB3114

BAB3114

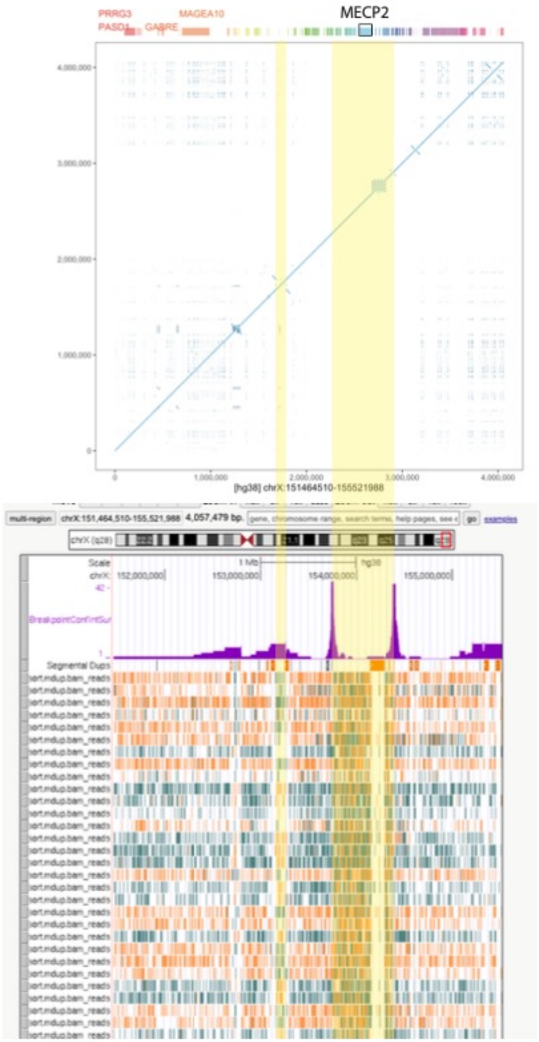


Inv InvDup

BAB14547

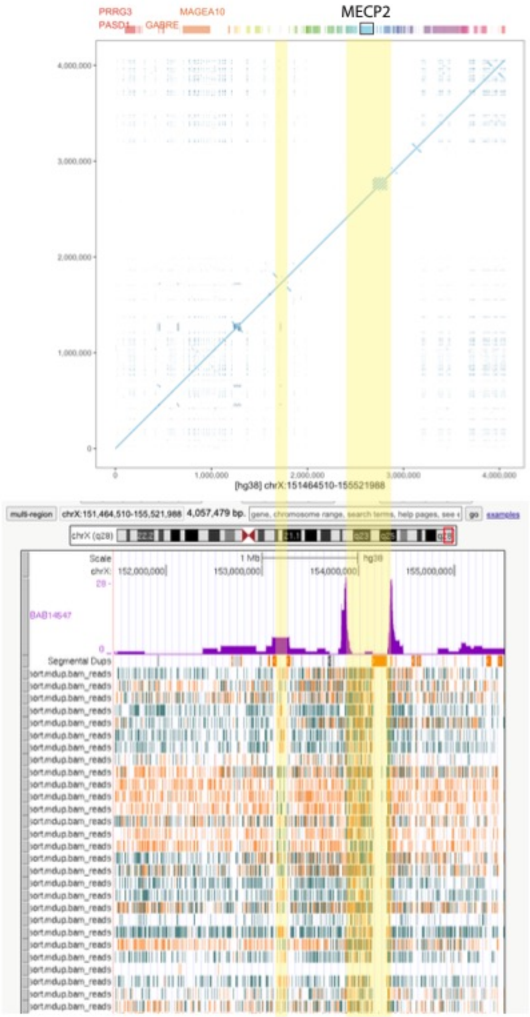


Inv InvDup

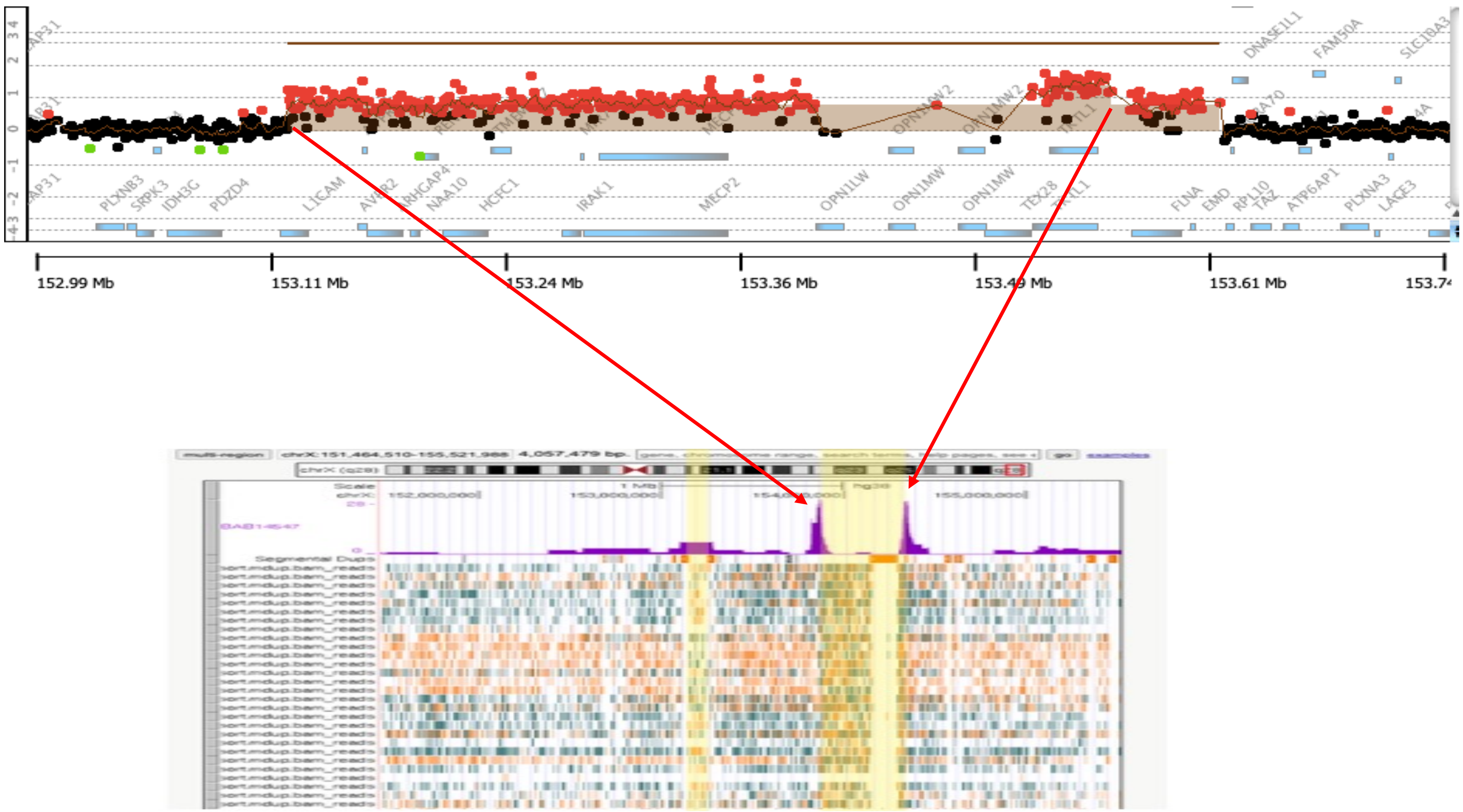

PLXNB3
SRPK3
IDH3G
PDZD4
L1CAM
AVPR2
ARHGAP4
NAA10
HCFC1
IRAK1
MECP2
OPN1LW
OPN1MW
OPN1MW
TEX28
FLNA
EMD
RPL10
TAZ
ATP6AP1
PLXNA3
LAGE3
DNASE1L1
FAM50A
152.99 Mb
153.11 Mb
153.24 Mb
153.36 Mb
153.49 Mb
153.61 Mb

### BAB14547

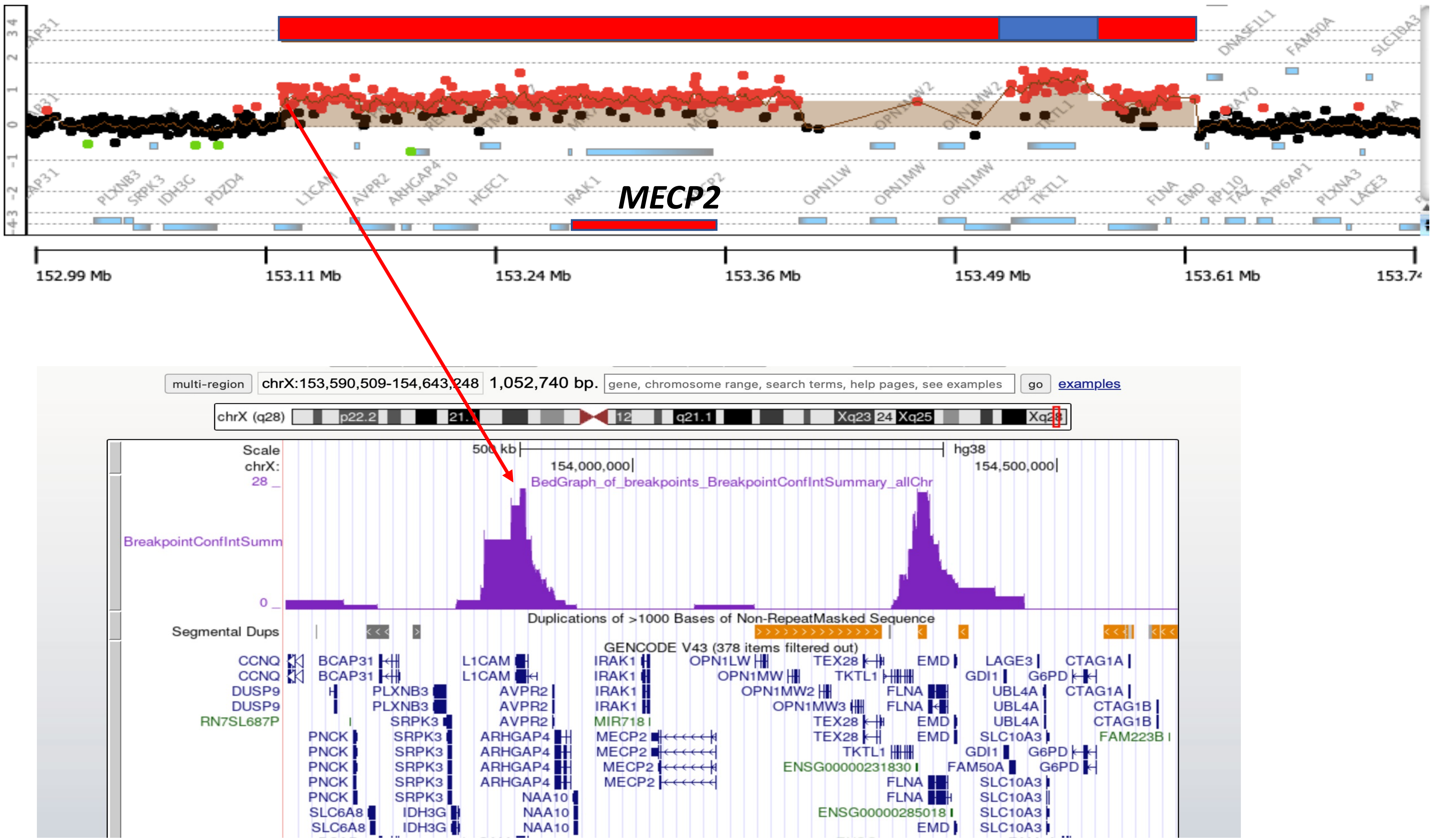

MECP2
152.99 Mb
153.11 Mb
153.24 Mb
153.36 Mb
153.49 Mb
153.61 Mb
multi-region
chrX:153,590,509-154,643,248
1,052,740 bp.
gene, chromosome range, search terms, help pages, see examples
go
examples
chrX (q28)
Scale
500 kb
hg38
chrX:
154,000,000
154,500,000
BedGraph_of_breakpoints_BreakpointConfIntSummary_allChr
BreakpointConfIntSumm
Duplications of >1000 Bases of Non-RepeatMasked Sequence
Segmental Dups
GENCODE V43 (378 items filtered out)

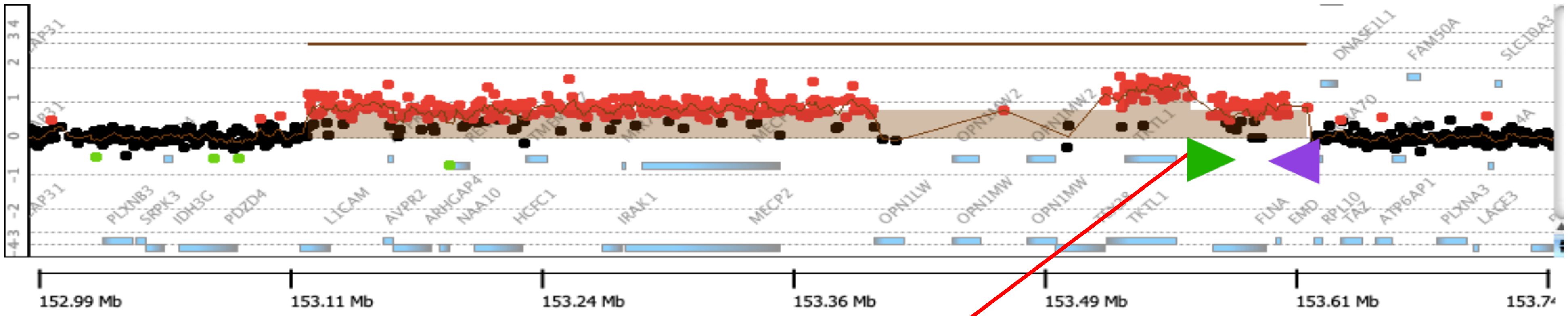

152.99 Mb
153.11 Mb
153.24 Mb
153.36 Mb
153.49 Mb
153.61 Mb


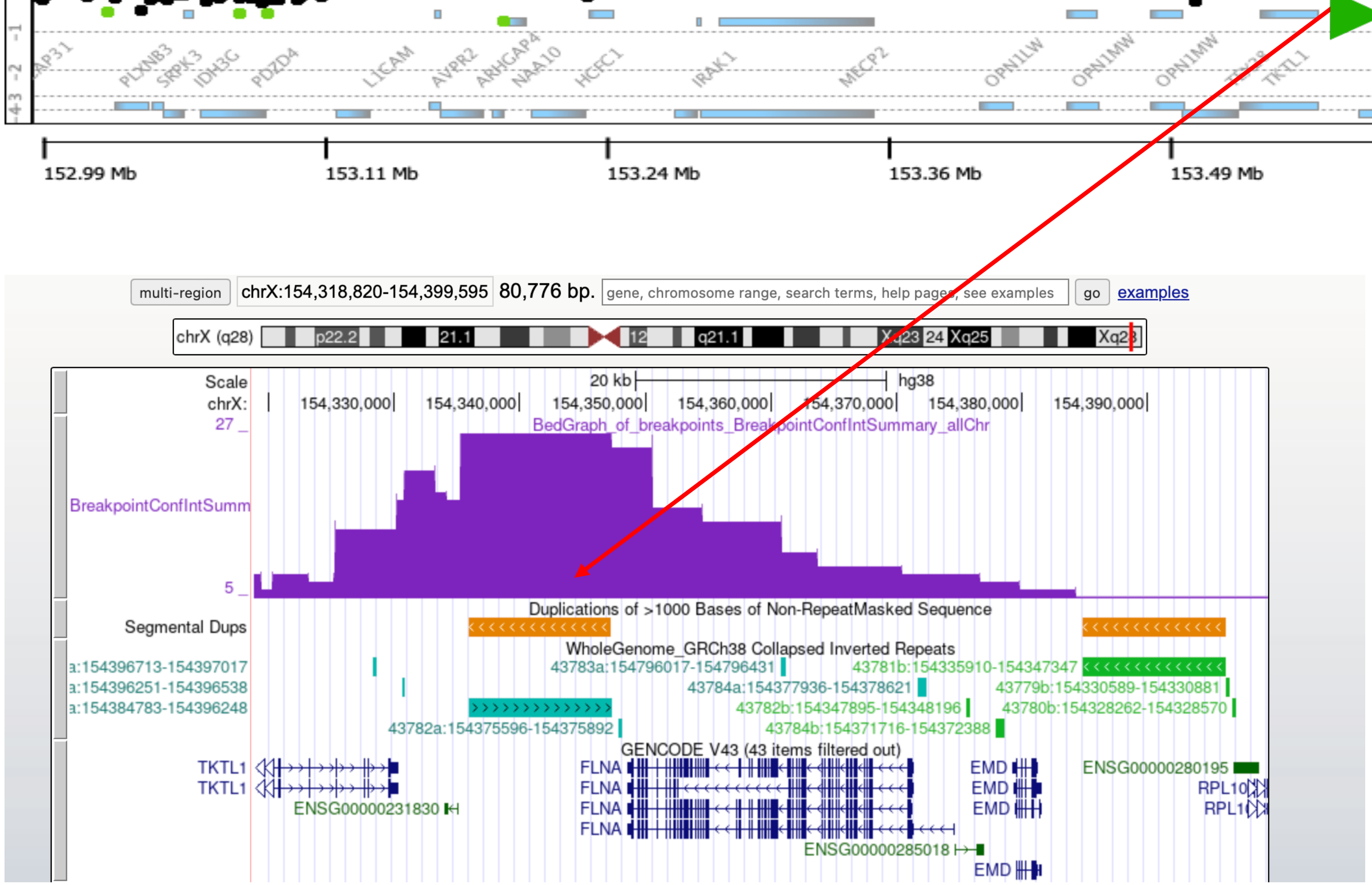

multi-region
chrX:154,318,820-154,399,595
80,776 bp.
go
examples
Scale
20 kb
hg38
BedGraph_of_breakpoints_BreakpointConfIntSummary_allChr
BreakpointConfIntSumm
Segmental Dups
Duplications of >1000 Bases of Non-RepeatMasked Sequence
WholeGenome_GRCh38 Collapsed Inverted Repeats
GENCODE V43 (43 items filtered out)
TKTL1
FLNA
EMD


### First LCR K1

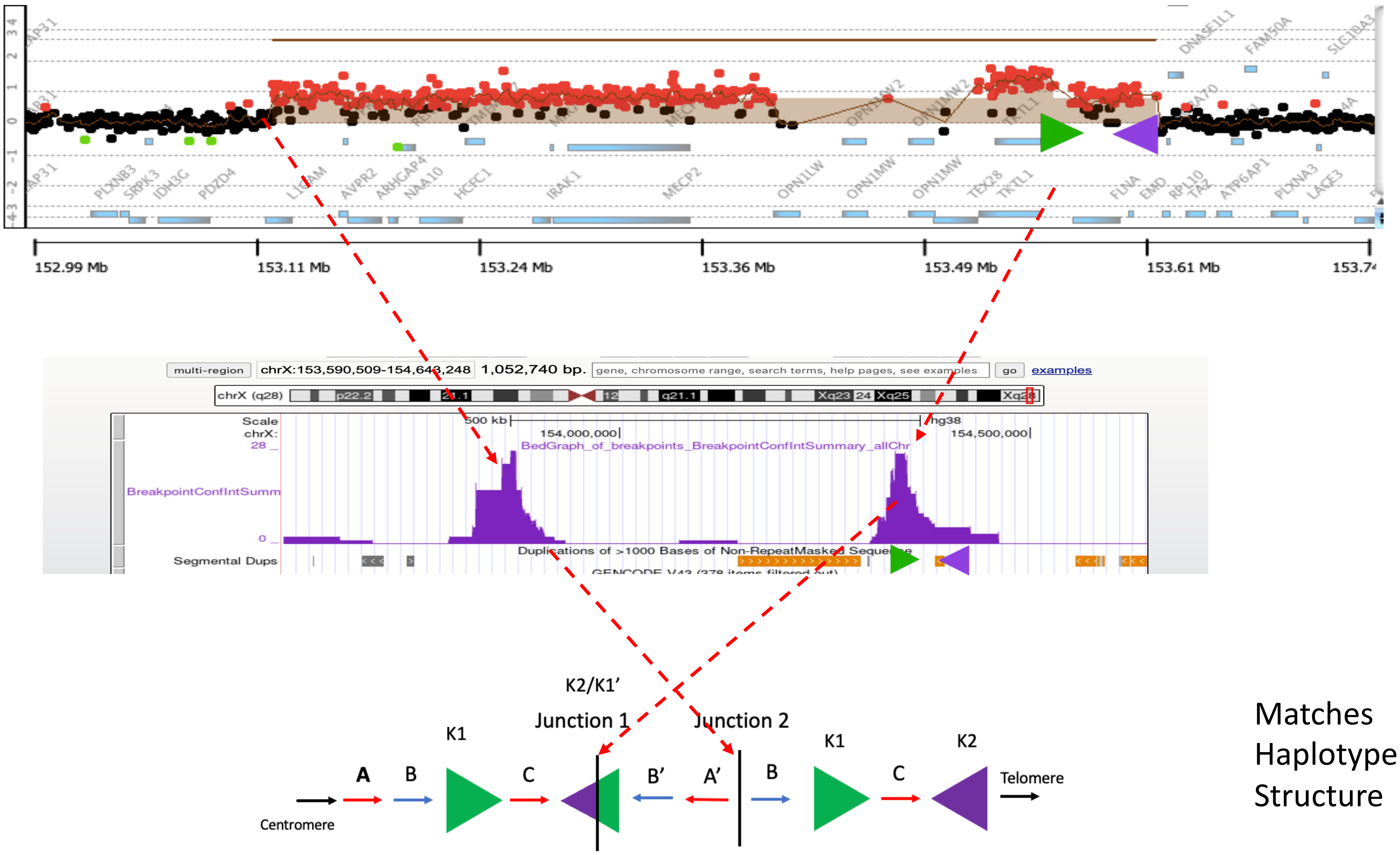


Matches Haplotype Structure

### BAB3114

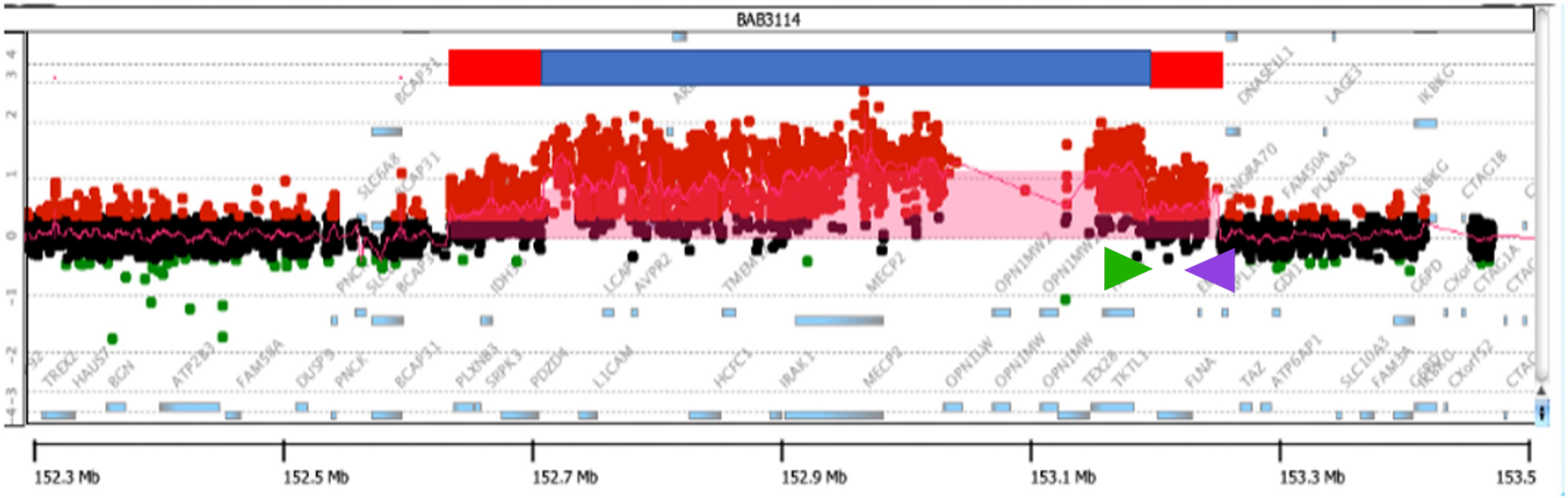


***Array Comparative Genomic Hybridization***

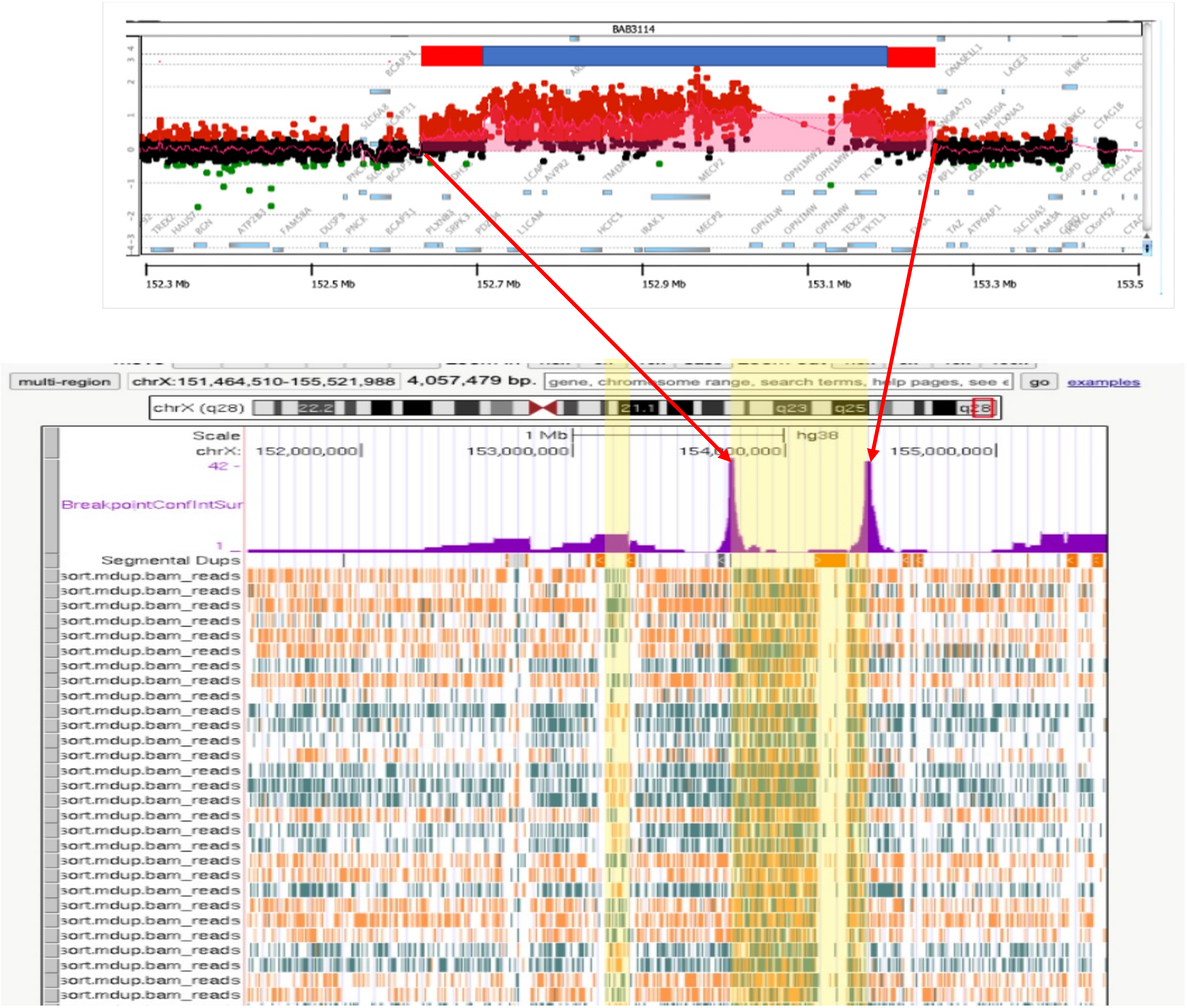

BAB3114
152.3 Mb
152.5 Mb
152.7 Mb
152.9 Mb
153.1 Mb
153.3 Mb
153.5
multi-region
chrX:151,464,510-155,521,988
4,057,479 bp.
go
examples
chrX (q28)
Scale
1 Mb
hg38
chrX:
152,000,000
153,000,000
154,000,000
155,000,000
BreakpointConfIntSur
Segmental Dups
sort.mdup.bam_reads

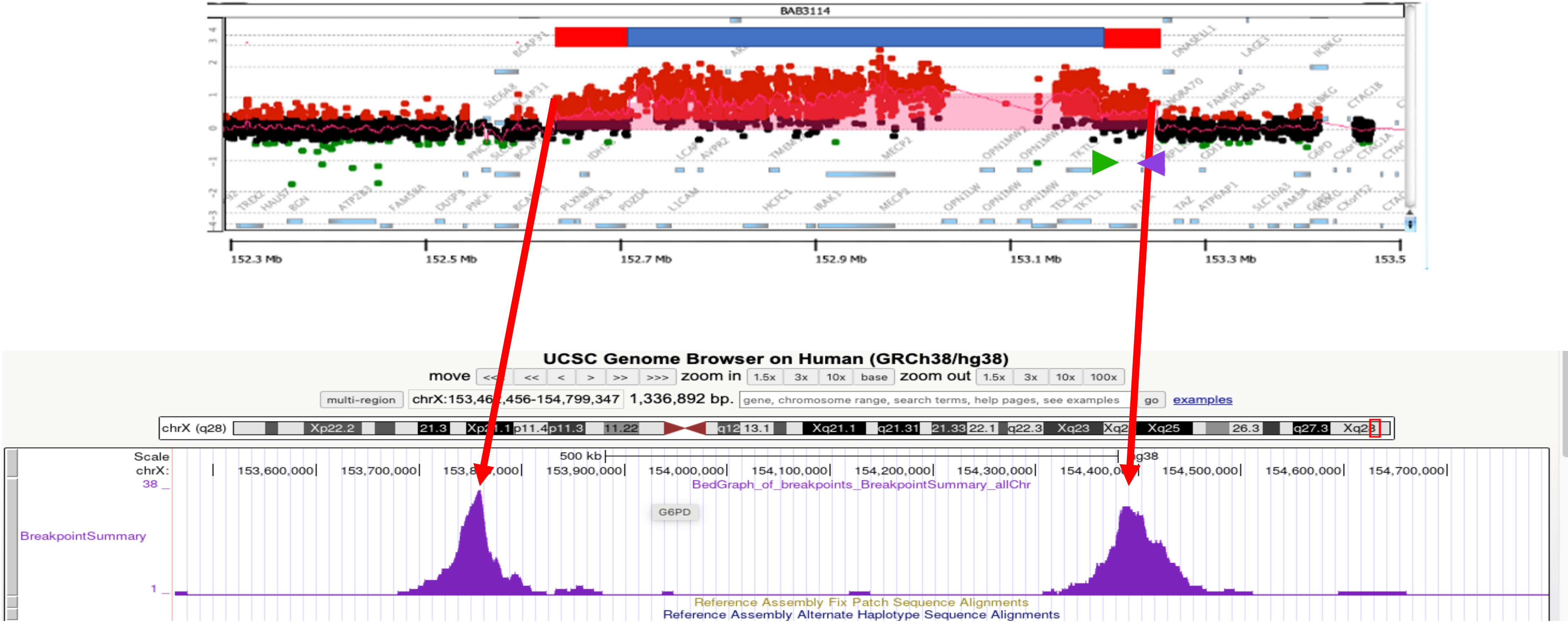
BAB3114
152.3 Mb
152.5 Mb
152.7 Mb
152.9 Mb
153.1 Mb
153.3 Mb
153.5
UCSC Genome Browser on Human (GRCh38/hg38)
move
zoom in
1.5x
3x
10x
base
zoom out
1.5x
3x
10x
100x
multi-region
1,336,892 bp.
gene, chromosome range, search terms, help pages, see examples
go
examples
chrX (q28)
Scale
chrX:
500 kb
153,600,000
153,700,000
153,900,000
154,000,000
154,100,000
154,200,000
154,300,000
154,500,000
154,600,000
154,700,000
38
BedGraph_of_breakpoints_BreakpointSummary_allChr
G6PD
BreakpointSummary
1
Reference Assembly Fix Patch Sequence Alignments
Reference Assembly Alternate Haplotype Sequence Alignments

### BAB3114

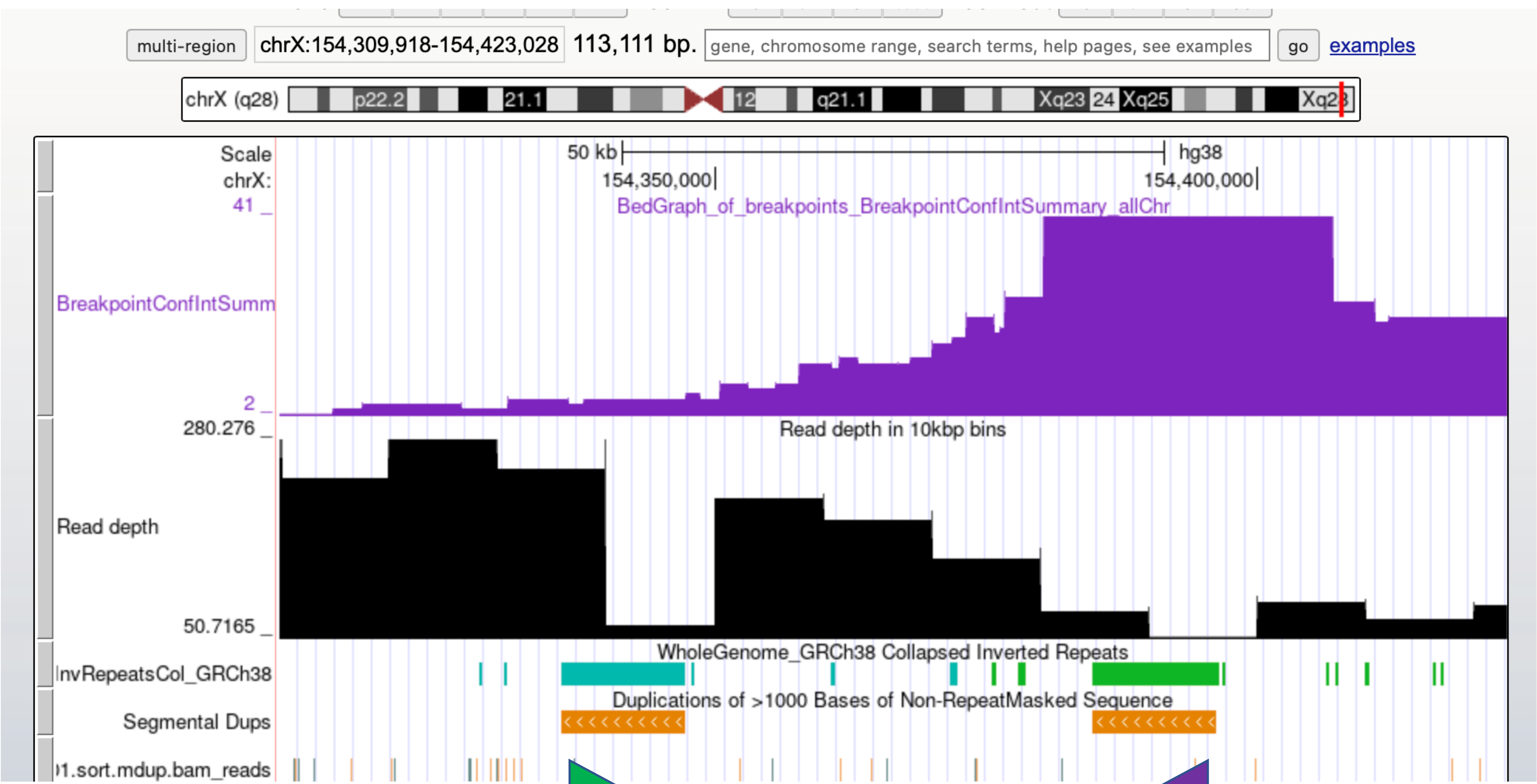

multi-region
chrX:154,309,918-154,423,028 113,111 bp.
gene, chromosome range, search terms, help pages, see examples
go
examples
chrX (q28)
p22.2
21.1
12
q21.1
Xq23
24
Xq25
Scale
50 kb
hg38
chrX:
154,350,000
154,400,000
41 _
BedGraph_of_breakpoints_BreakpointConfIntSummary_allChr
BreakpointConfIntSumm
2 _
280.276 _
Read depth in 10kbp bins
Read depth
50.7165 _
WholeGenome_GRCh38 Collapsed Inverted Repeats
InvRepeatsCol_GRCh38
Duplications of >1000 Bases of Non-RepeatMasked Sequence
Segmental Dups
1.sort.mdup.bam_reads


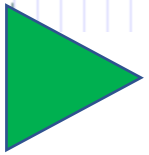

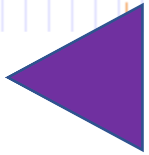

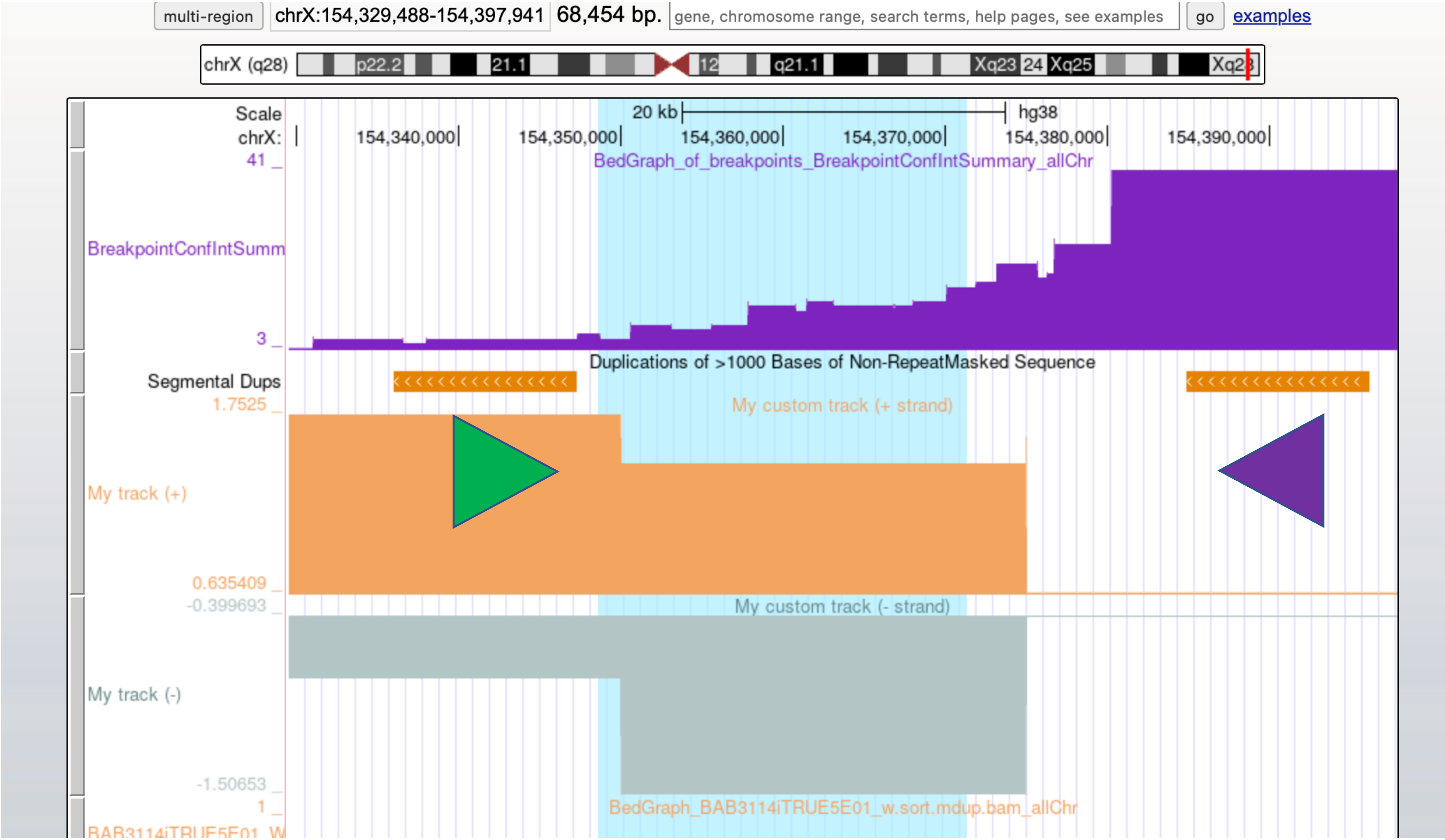
multi-region
chrX:154,329,488-154,397,941
68,454 bp.
gene, chromosome range, search terms, help pages, see examples
go
examples
chrX (q28)
p22.2
21.1
12
q21.1
Xq23
24
Xq25
Scale
20 kb
hg38
chrX:
154,340,000
154,350,000
154,360,000
154,370,000
154,380,000
154,390,000
41 _
BedGraph_of_breakpoints_BreakpointConfIntSummary_allChr
BreakpointConfIntSumm
3 _
Duplications of >1000 Bases of Non-RepeatMasked Sequence
Segmental Dups
1.7525 _
My custom track (+ strand)
My track (+)
0.635409 _
-0.399693 _
My custom track (- strand)
My track (-)
-1.50653 _
1 _
BedGraph_BAB3114iTRUE5E01_w.sort.mdup.bam_allChr

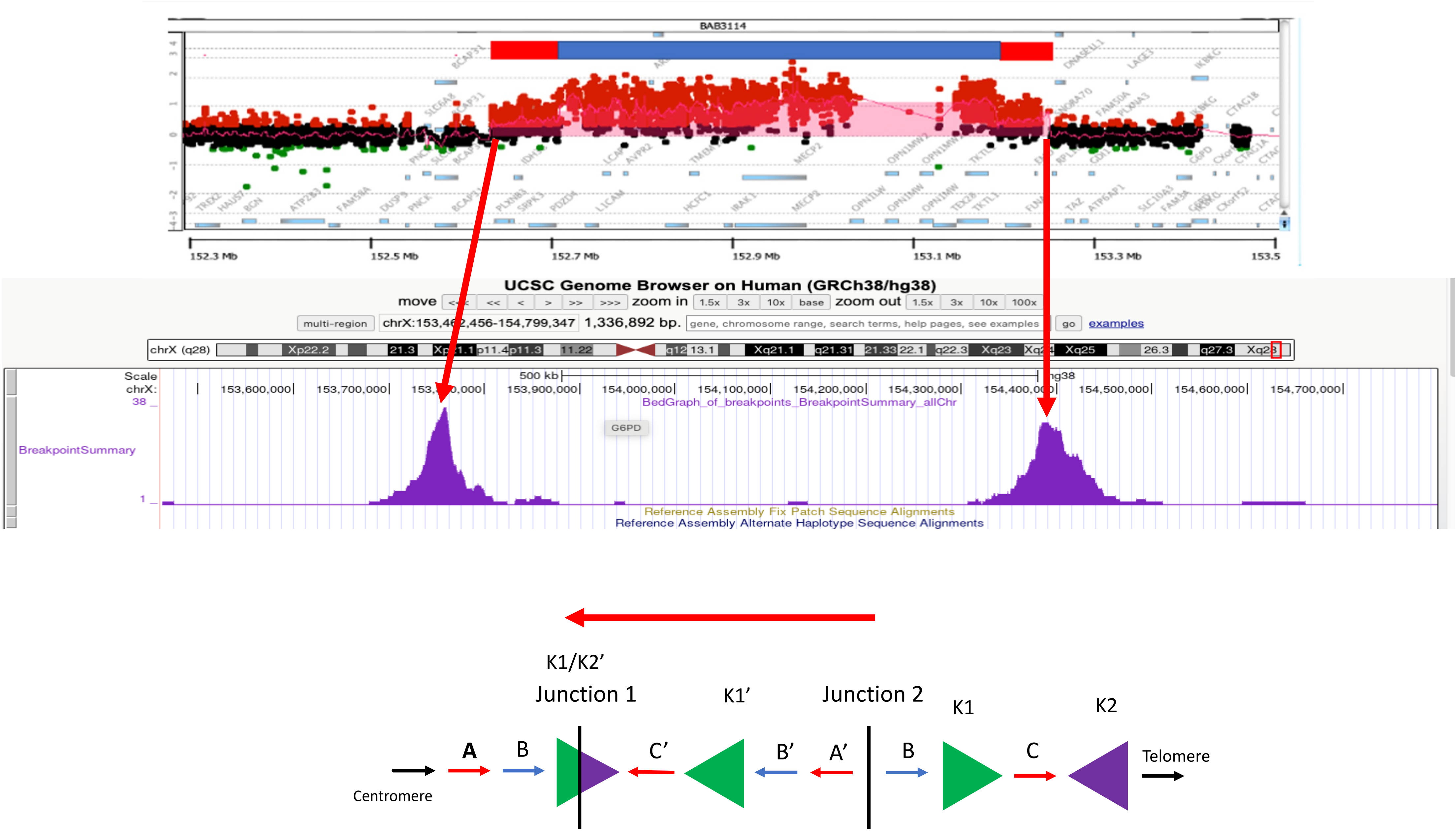

BAB3114
152.3 Mb
152.5 Mb
152.7 Mb
152.9 Mb
153.1 Mb
153.3 Mb
153.5
UCSC Genome Browser on Human (GRCh38/hg38)
chrX:153,462,456-154,799,347 1,336,892 bp.
BreakpointSummary
BedGraph_of_breakpoints_BreakpointSummary_allChr
G6PD
Reference Assembly Fix Patch Sequence Alignments
Reference Assembly Alternate Haplotype Sequence Alignments
K1/K2'
Junction 1
K1'
Junction 2
K1
K2
A
B
C'
B'
A'
B
C
Centromere
Telomere
